# Plant genetic bases explaining microbiota diversity shed light into a novel holobiont generalist gene theory

**DOI:** 10.1101/2023.12.22.572874

**Authors:** Loeiz Maillet, Manon Norest, Adam Kautsky, Anna Geraci, Elisabetta Oddo, Angelo Troia, Anne-Yvonne Guillerm-Erckelboudt, Cyril Falentin, Mathieu Rousseau-Gueutin, Anne-Marie Chèvre, Benjamin Istace, Corinne Cruaud, Caroline Belser, Jean-Marc Aury, Rosario Schicchi, Léa Frachon, Claudia Bartoli

**Author notes:** **Author Contributions:** LM & CB wrote the manuscript, LM, CB & LF did the bioinformatic and statistical analysis, A-MC, AK, A-Y G-E, CF, CB, AG. EO, AT & RS collected plants and ecological data, AK, CB & MN did the molecular biology experiments, BI, CC, CF, MR-G, CB & J-MA funded, sequenced and assembled the *Brassica napus* reference genome, CB found funds and directed the research. All the authors reviewed the manuscript. **Competing Interest Statement:** The authors declare no competing interest.

## Abstract

Plants as animals are strictly associated with a cortege of microbial communities influencing their health, fitness and evolution. Therefore, scientists refer to all living organisms as holobionts; complex genetic units that coevolve simultaneously. This is what has been recently proposed as the hologenome theory of evolution. This exciting and attractive theory has important implications on animal and plant health; however, it still needs consistent proof to be validated. Indeed, holobionts are still poorly studied in their natural habitats where coevolution and natural selective processes occur. Compared to animals and crops, wild plant populations are an excellent and unique model to explore the hologenome theory. These sessile holobionts have strictly coevolved with their microbiota for decades and natural selection and adaptive processes acting on wild plants are likely to regulate the plant-microbe interactions. Here we conducted for the first time a microbiota survey, plant genome sequencing and Genome-Environmental Analysis (GEA) of 26 natural populations of the non-model plant species *Brassica rapa*. We collected plants over two seasons in Italy and France, and analyzed the microbiota on two plant compartments (root and rhizosphere). We identified that plant compartment and season modulate *B. rapa* microbiota. More importantly, when conducting GEA we evidenced neat peaks of association correlating with both fungal and bacterial microbiota. Surprisingly, we found 13 common genes between fungal and bacterial diversity descriptors that we referred under the name of Holobiont Generalist Genes (HGG). These genes might strongly regulate the diversity and composition of plant microbiota at the inter-kingdom level.

**Significance Statement:** The novel hologenome concept claims that hosts and their associated microbes (considered as holobionts) are a unique evolutionary unit on which natural selection acts. Thus, the hologenome theory assumes that hosts and microbiomes simultaneously coevolve. This novel vision of universal evolutionary entities is promising for both animal and plant health purposes. However, it is still quietly controversial as it suffers from a lack of tangible evidence. How can we enrich the debate on holobionts? How can we translate this concept in discoveries that can change farming practices? Our study is filling the gaps of the hologenome theory by showing that certain genes under natural selection and regulating plant microbiota are generalist in response to fungal and bacterial communities.

## Introduction

With no exceptions, all living organisms are associated with microorganisms (viruses, bacteria, fungi, oomycetes) that are commonly referred to as the host microbiome. Recently, scientists coined the terminology of holobiont for describing hosts and their associated microbes as a unique entity on which natural selection operates. Therefore, there is a significant shift in the way that evolution is considered by evolutionary scientists; indeed, animals and plants are no longer a centric and sole protagonist of evolution but rather an entity much more complex and universal (1–3). More largely, the evolutionary core is not the individual genome anymore but what is called the hologenome, i.e. the genome of the host and of all associated microscopic organisms. Accepting the hologenome as a valid biological unit is equivalent to assuming that selection directly acts on holobionts as they are unique entities in constant coevolution (4). But how has this exciting and promising theory of evolution been studied so far? In animals, and particularly humans, the role of microbiota in regulating health has been extensively and deeply studied. However, we are still missing tangible evidence on the holobiont/hologenome theory.

Compared to animals, sessile organisms, such as plants, are an exceptional resource to study the hologenome concept as they are forced to adapt to the fluctuating environment where they evolve. Therefore, they strongly coevolve with soil microbes living in their ecological habitats. It has been proposed, but not entirely proved, that plants can modulate their microbiomes to dynamically adjust to their habitats (5). If confirmed, this observation can hugely change agriculture as we could breed crop varieties specifically selecting for beneficial microbes. As the host plant genome slowly evolves and the microbiota quickly change (6), the use of the holobiont concept in agriculture can improve crop systems in a short-time scale. However, the still unanswered question is: how plant holobionts can be studied and how information on the coevolution between hosts and microbiomes can improve agriculture? In the current genomic era, two main approaches can be adopted to improve knowledge on holobionts. Firstly, experimental evolution (7, 8) coupled with genomics can help identifying rapid evolutionary changes in both host and microbiome genomes. Experimental evolution is definitely a powerful tool, still underestimated and underused; however, it only allows the study of holobionts on a short ecological and evolutionary time-frame. Alternatively, Genome-Environmental-Analysis (GEA) allows to capture an entire picture of natural evolutionary processes in holobionts. Indeed, GEA can identify significant associations between the host genetic polymorphisms and environmental variables such as microbiome descriptors. As mentioned in Fitzpatrick (2020) “The next stage in plant microbiome research will require the integration of ecological and reductionist approaches to establish a general understanding of the assembly and function in both natural and managed environments” (9). In light of this, GEA is an alternative to identify the host genes involved in adaptive processes of the holobiont unit. Recently, studies showed the power of GEA to identify plant loci associated with biotic communities. This has been demonstrated both in the non-model plant species *B. incana* and its associated pollinator communities (10), and in the model plant species *Arabidopsis thaliana* and its plant-plant (11) and plant-microbiome (12) interactions. The latter study is the only describing adaptive loci associated with bacterial descriptors in natural plant populations. So far, GEA studies on plant-microbiome models are rare, especially on non-model plant species. *Brassica* species are a great genetic resource as wild *Brassica* populations - closely related to crops - still exist in a wide range of habitats. This fascinating genetic material is still poorly studied and can open the avenue into the elucidation of holobionts’ evolution (13).

Here we focus on the genetic architecture of the plant-microbiome interaction in wild *B. rapa* populations, a non-model plant and one of the two diploid progenitors of oilseed rape (*B. napus*). Wild *B. rapa* populations are closely related to various cultivated forms including turnips (*B. rapa* subsp. *rapa*). We ecologically and genetically characterized 26 wild *B. rapa* populations and produced information on the fungal and bacterial microbiome over two seasons and plant compartments (root and rhizosphere). We adopted a GEA approach and found that genetic bases controlling microbiota diversity and composition are shared between fungal and bacterial communities. Here we refer to microbiota shared genes as “Holobiont Generalist Genes (HGG)” since they regulate holobionts at different kingdoms levels.

## Results

### Genetic and ecological description of *B. rapa* natural populations showed highly genetically differentiated populations in the south of Italy

To explore the long-term coevolution between plants and their associated microbes, we chose to work on wild *B. rapa* populations as, compared to crops, they are locally adapted to their habitats and microbiomes. Twenty-six populations were collected during 2020 in both the south of Italy - Sicily (20 populations) - and the north of France - Brittany and Pays de la Loire (6 populations) - (Fig. 1A, B, C, Dataset S1). Plant habitats were selected to maximize the ecological variability (Fig. 1D) in terms of climatic, soil, land use, altitude, and plant community’s diversity (Dataset S1). Plant collection is part of a larger H2020 Prima project “BrasExplor” aiming at characterizing the genetic diversity of *Brassica* species in the Mediterranean basin (13). During fall 2020, we collected 30 plants for each of the 26 populations and sequenced each of them using a PoolSeq approach. We obtained 86,989,331.69 (average), 88,727,527.00 (median), 4,937,501.74 (standard deviation) reads per library (Dataset S2). Raw data were filtered based on sequences’ quality. After trimming, we obtained 86,964,161.23 (mean), 88,677,517.00 (median), 4,940,310.81 (SD) reads. Prior genome mapping, we sequenced the reference genome of *Brassica napus* RCC-S0 (SI Appendix), a synthetic line composed by *B. rapa* C1.3 A-subgenome. We adopted this method as *B. rapa* C1.3 is a variety genetically close to the wild *B. rapa* populations (SI Appendix, Dataset S3, Fig. S1). Wild *B. rapa* population genomes were mapped on the *B. rapa* C1.3 genome. After mapping and SNPs calling, we obtained a matrix of 12,518,450 SNPs that were trimmed. The final data matrix was composed by 1,138,924 SNPs.

**Figure 1.**
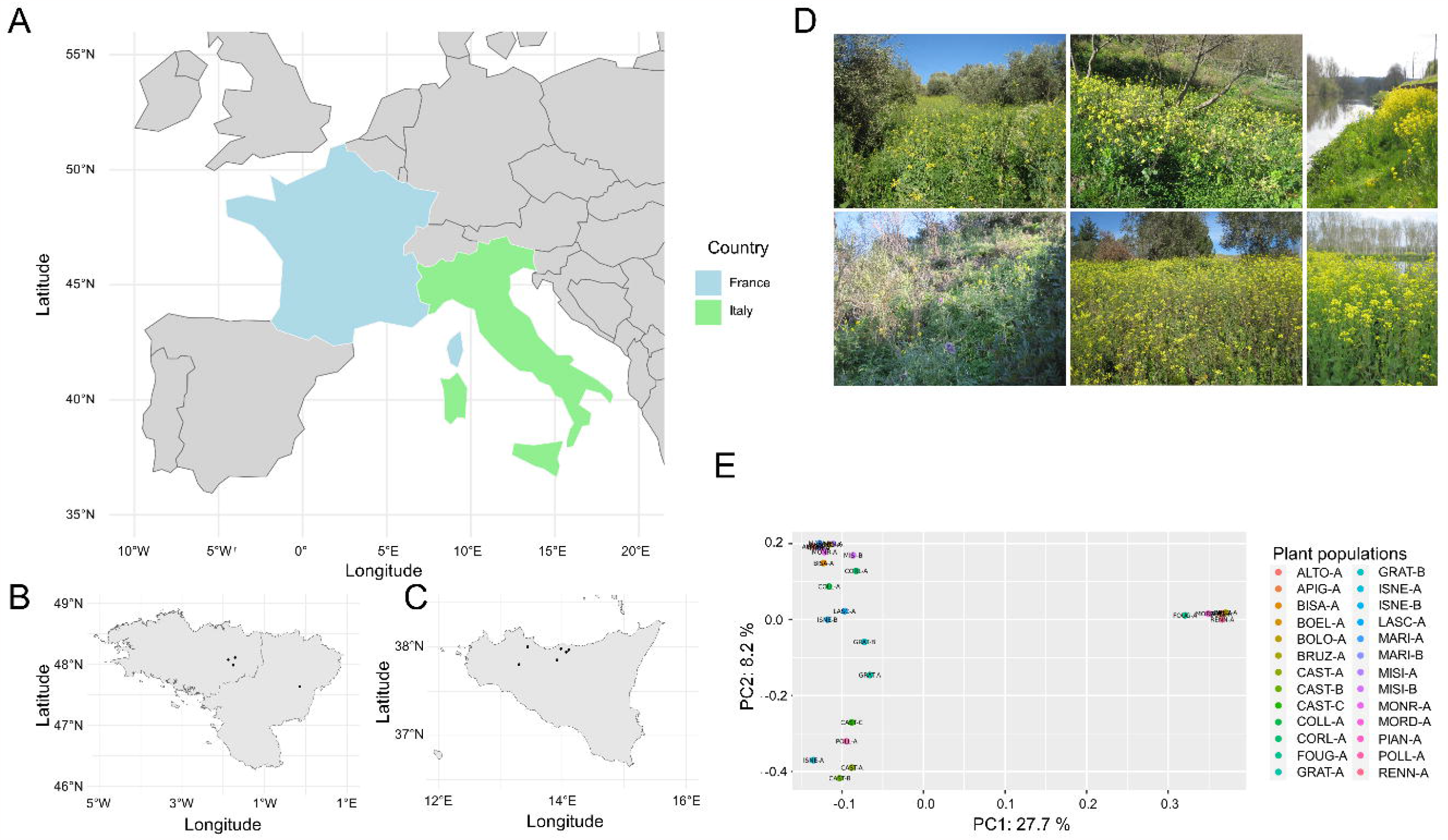
Ecological description of *Brassica rapa* natural *populations* collected in this study. **A**) Map of western Europe highlighting France (in light blue) and Italy (green). **B**) Map of Brittany and Pays de la Loire; dark plots represent the six French *B. rapa* populations. **C**) Map of Sicily; dark plots represent the 20 Italian *B. rapa* populations. **D**) Examples of the ecological habitats where *B. rapa* populations were collected in both Italy (two upper-left and two bottom-left panels) and France (upper-right and bottom-right panels). **E**) Principal-Component-Analysis (PCA) built on the SNPs matrix of the 26 plant populations; each population is represented with a different colored dot. Percentage of the explained variance for the two first principal components (PC) (*x-axis* indicates the first principal component, and *y-axis* indicates the second principal component) is reported. The Italian populations are all distributed on the left side of the PCA panel, otherwise French populations are all distributed on the right side of the PCA panel.

The genomic Principal Component Analysis (PCA) on the SNPs matrix of the 26 populations (Fig. 1E) showed that principal component 1 partly explains the variation among the countries, and principal component 2 the variation within each country (mostly for the Italian populations). We observed that the Italian populations were more scattered in the genomic space compared to those collected in France (Fig. 1E). This is probably related to the ecological diversification of the habitats where plants were collected. Indeed, populations from Sicily were collected in highly diversified habitats ranging from abandoned fields to olive orchards (Fig. 1D, Dataset S1) and showing a larger diversity in terms of plant communities (Fig. S2, Dataset S1). On the other hand, French populations were all collected closer to rivers (Fig. 1D) and habitat characterization showed, as expected, that plant communities were extremely different to those observed in Sicily (Fig. S2, Dataset S1). Nevertheless, other factors, e.g. the *B. rapa* populations demography history, might influenced the genomic differentiations of the Sicilian populations.

### *Brassica rapa* microbiota descriptors are influenced by plant compartment, season and plant genotype

To study the interactions within the *B. rapa* populations and their associated microbiomes, we conducted an *in situ* microbiome survey over two seasons: i) spring (SP), corresponding to February 2020 in Sicily, and May 2020 in France - plants were at the flowering stage – and ii) fall (FA), corresponding to December 2020 in Italy, and January 2021 in France; all plant populations were at the 4-leaves stage in fall. For microbiome description we collected 6 plants per population and we separated root (R) and rhizosphere (RH) compartments. We amplified a portion of ITS1 and *rpoB* (14) for fungal and bacterial characterization communities respectively. For bacterial communities, root samples were removed because of their poor raw data quality. Concerning the fungal communities, we observed a strong plant compartment effect on diversity *α*-descriptors but not on the fungal composition (Fig. 2A, Fig. S3, Table S1). Not surprisingly, fungal diversity was higher in the rhizosphere than in the root compartment. We also evidenced a significant effect of the season variable on fungal diversity but not on composition. This effect was significant in the root compartment but not in the rhizosphere (Fig. 2A, 2B, Table S2). As both season and plant compartment strongly influenced fungal communities, analyses were run on four datasets to test whether plant genotype impacts on microbiome descriptors: root in fall (R-FA), root in spring (R-SP), rhizosphere in fall (RH-FA), and rhizosphere in spring (RH-SP). By running a transformation-based Canonical Redundancy Analysis (tb-RDA) and linear-mixed models on the Principal Component Analysis (PCoA) axis, we identified a strong plant population effect on the fungal composition on both R and RH compartments and over the two seasons (Fig. 2C, Table S3, Table S4). The plant population variable also strongly influenced *α*-diversity descriptors (Fig. 2D, Fig. S4, Table S4).

**Figure 2.**
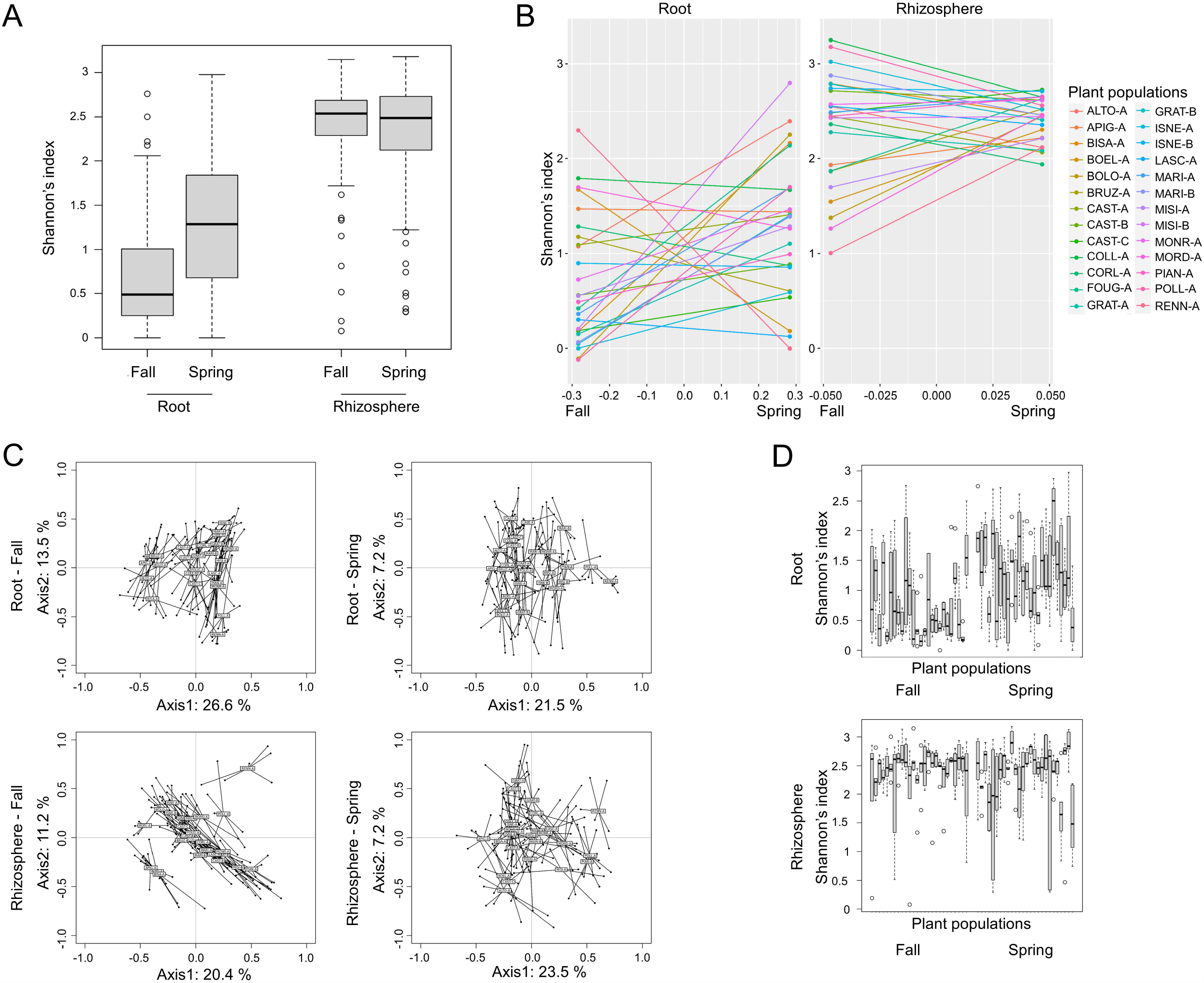
Fungal microbiota descriptor analysis. **A**) Box plots on the Shannon’s index (*y*-axis) showing the variation of fungal *α*-diversity within plant compartments (root and rhizosphere, *x*-axis) and seasons (fall and spring, *y*-axis). **B**) Finlay Wilkinson’s regression for the Shannon’s index of the 26 *Brassica rapa* plant populations over two seasons (fall and spring) and for each plant compartment (left: root, right: rhizosphere). **C**) Transformation based-Redundancy Analysis (tbRDA) plot representing the fungal composition variation within the 26 *B. rapa* populations run on four datasets: top-left: root-fall, top-right: root-spring, bottom-left: rhizosphere-fall, bottom-right: rhizosphere-spring. **D**) Box plots illustrating the seasonal effect on the Shannon’s index (*y*-axis) estimated on root (top panel) and rhizosphere (bottom panel) compartments over two seasons and for each plant population. Each plot represents the Shannon’s index of a specific plant population (*x*-axis).

For bacterial communities we only described the rhizosphere microbiome as a large majority of root samples were characterized by less than 1000 reads after sequencing. As for the fungal microbiome, we did not observe a seasonal effect on rhizosphere samples (Fig. 3A, 3B, Fig. S5, Table S5). On the other hand, we observed a significant effect of the plant population variable on both bacterial community composition (Fig. 3C, Table S4, Table S6) and α-diversity (Fig. 3D, Table S4). Globally, for both fungal and bacterial communities we observed a significant plant population effect on both *α*-diversity descriptors and composition. We therefore estimated the broad-sense heritability (H^2^) on microbiota descriptors (Shannon’s index, species richness and the two first axis of the PCoA) and observed high values of H^2^ for most of the fungal and bacterial descriptors (Table S7). These results claim a crucial role of plant genetic diversity in modulating both fungal and bacterial communities. We then coupled information on plant population genetic diversity with microbiome descriptor analysis to better define the genetic architecture of *B. rapa* holobiont.

**Figure 3.**
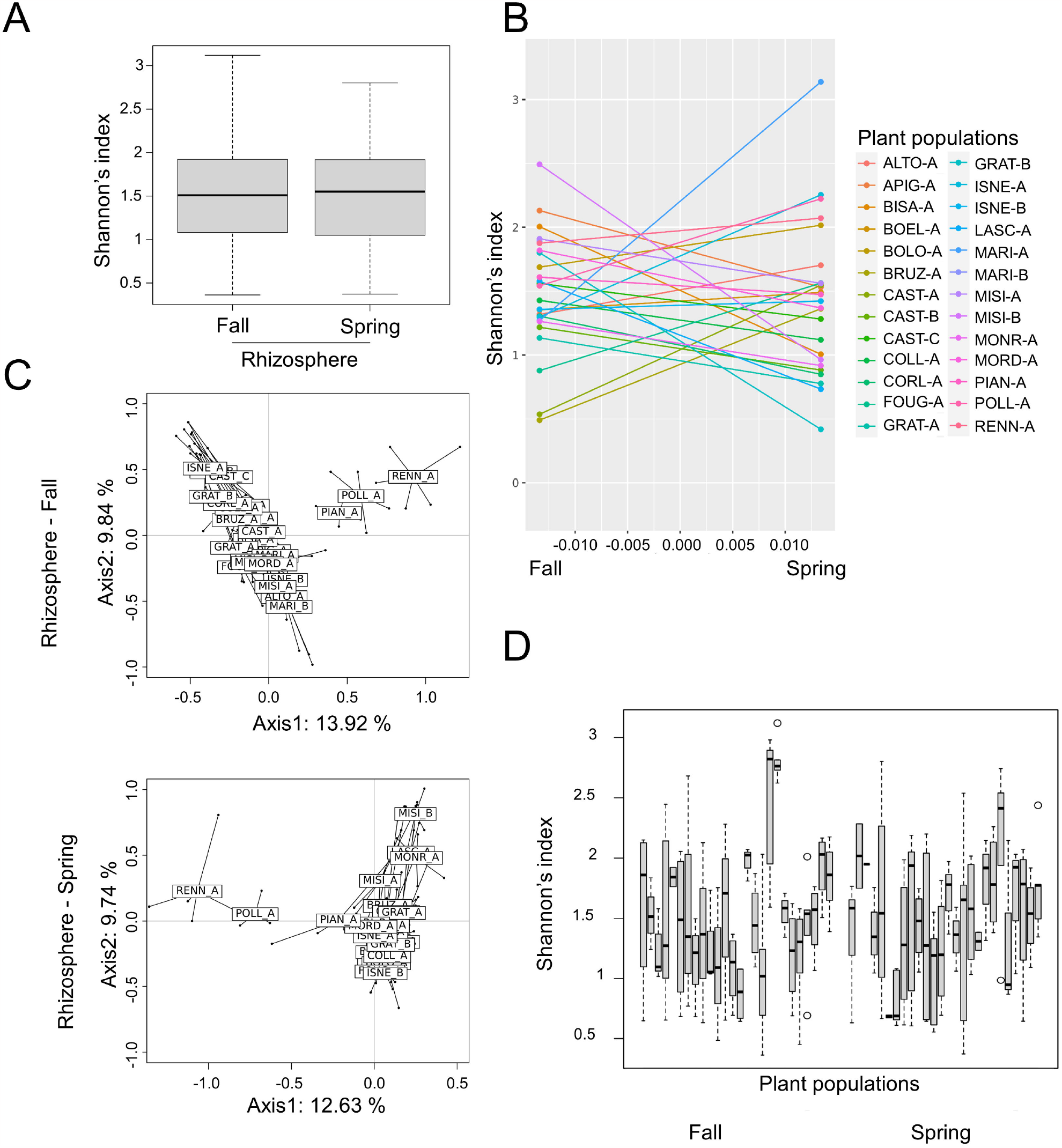
Bacterial microbiota descriptor analysis. **A**) Box plots on the Shannon’s index (*y*-axis) showing the variation of bacterial *α*-diversity within two seasons (fall and spring *x*-axis). **B**) Finlay Wilkinson’s regression of the 26 *Brassica rapa* plant populations over two seasons (fall and spring) **C**) Transformation based-Redundancy Analysis (tbRDA) representing the bacterial composition variation within the 26 *B. rapa* populations run on two datasets: fall (top panel) and spring (bottom panel). **D**) Box plots illustrating the seasonal effect on the Shannon’s index (*y*-axis) estimated over two seasons and for each plant population. Each plot represents the Shannon’s index of a specific plant population (*x*-axis).

### Genome-environmental-analysis showed a highly diversified plant genetic architecture within plant compartments and seasons as association with plant resistance genes

To identify the genetic bases associated with both fungal and bacterial community descriptors, we adopted a GEA approach by combining a Bayesian Hierarchical Model (BHM) that decreases the rate of false genotype-ecology positive associations (15), with a Local Score (LS) approach allowing the detection of QTLs with small effects (16). Best-Linear-Unbiased-Predictions (BLUPs) were estimated on *α*-microbiota descriptors, community composition descriptors, and on the relative abundance of the most prevalent OTUs (Dataset S5-S10). The plant trimmed SNPs matrix was then integrated in the BHM model that estimates a covariance matrix minimizing for population structure; BLUPs were integrated as covariables to estimate the associated SNPs. After running the LS approach, we selected only QTLs that had a significant local score value (Lindley score), and we retrieved genes falling in these QTLs. For both fungal and bacterial communities, we found that the genetic architecture is highly variable among seasons and plant compartments, as few genes were found to be shared within root and rhizosphere samples, and within fall and spring when analyzing fungal and bacterial communities separately (Fig. S6, S7).

Regarding fungal microbial descriptors, we found neat peaks of association (i.e. carrying high LS) falling in genome regions corresponding to annotated *B. rapa* genes (Dataset S11, S12). As very little information is available on *B. rapa* gene function, we searched their orthologs in the closely related plant model species *Arabidopsis thaliana*. We then retrieved the known functions on TAIR website https://www.arabidopsis.org/. Among genes falling into the most significant association peaks, we found the A01p01520.1_BnaRCC gene associated with the relative abundance of the fungal OTU000080 that corresponds to the well-known plant-protecting fungal species *Trichoderma* sp. We also found that the A02p35100.1_BnaRCC gene was associated with the fungal PCoA axis 4 (Dataset S11, S12). Both genes correspond to members of the cytochrome P450 (CYP) in *A. thaliana*. CYP is an ancient superfamily of enzymes involved in multiple catalytic pathways and identified in all domains of living organisms (17). We evidenced a neat peak of association on chromosome 7 to be associated with the OTU000048 belonging to the fungal species *Preussia flanaganii*. We retrieved the gene A07p21630.1_BnaRCC (Dataset S11, S12) corresponding in *A. thaliana* to the Basic Helix-Loop-Helix protein 100 (bHLH100), which is one of the largest transcription factor gene families in *Arabidopsis* and which is involved in the resistance to abiotic stresses (18). We identified several SNPs within genes known to be related to plant resistance pathways. For instance, A01p08480.1_BnaRCC is associated with the relative abundance of the fungal OTU000050 *Fusicolla* sp. (Dataset S11, S12). Its *A. thaliana* orthologous gene is the IBI1 receptor, which is responsible for BABA-induced resistance. Also, we found A09p17430.1_BnaRCC to be associated with the relative abundance of the fungal OTU000079 belonging to the *Vishniacozyma* (Dataset S11, S12) species that are known to induce resistance to fungal pathogens. Its *A. thaliana* ortholog corresponds to the BURNOUT1 gene that is related to several plant stress responses.

We also found several neat peaks associated with bacterial descriptors (Dataset S11). For example, we identify a significant peak of association in the A07p36280.1_BnaRCC gene (encoding a deoxyribonucleoside kinase: ATDNK) to be associated with bacterial species richness (Dataset S11, S12). Interestingly, we found a significant peak on chromosome 9 to be associated with the bacterial species *Bacillus simplex* (Dataset S11, S12) described to be a biocontrol agent against *Fusarium* fungal pathogens and other plant pathogens as most *Bacillus* species (19, 20). The peak fell in the A09p41890.1_BnaRCC gene, whose *A. thaliana* ortholog correspond to the TIP41-like protein that was shown to be involved in the plant defense against bacterial pathogens (21).

### Common genetic bases associated with both fungal and bacterial communities drive into a novel theory of Holobiont Generalist Genes (HGG)

One of the aims of our study was to explore whether genetic bases associated with microbiome descriptors are shared among the bacterial and fungal communities. We therefore intersect genes found to be associated with fungal and bacterial descriptors. Surprisingly, when merging all compartments and seasons, we found that 13 genes were shared among microbiome descriptors (Fig. 4). For instance, we found a neat peak within the A04p30810.1_BnaRCC gene associated with both relative abundance of the fungal species *Cystofilobasidium macerans* and the bacterial species richness (Fig. 5). This latter gene was also associated with the relative abundance of *Curvularia spicifera* and the ITS PCoA axis4 (Dataset S11). Its Arabidopsis ortholog is AT2G43610, which belongs to a chitinase family protein and that is involved in root *Arabidopsis* morphogenesis (22). A neat peak of association was found within A06p13350.1_BnaRCC gene associated with both relative abundance of the fungal species *Pseudeurotium* sp. and the relative abundance of the bacterium *Arthrobacter* sp (Fig. 5). The *A. thaliana* homologous AT1G18160 belongs to the M3KDELTA7 family that is involved in complex osmotic-stress and abscisic acid (ABA) signal transduction network (23). Interestingly, we observed that the A09p41890.1_BnaRCC gene, that is the homologous of a TIP41-like, was shared among fungal (*Preussia flanaganii* relative abundance) and bacterial (species richness and *Bacillus simplex* abundance) microbiome descriptors (Fig. 4, Dataset S11).

**Figure 4.**
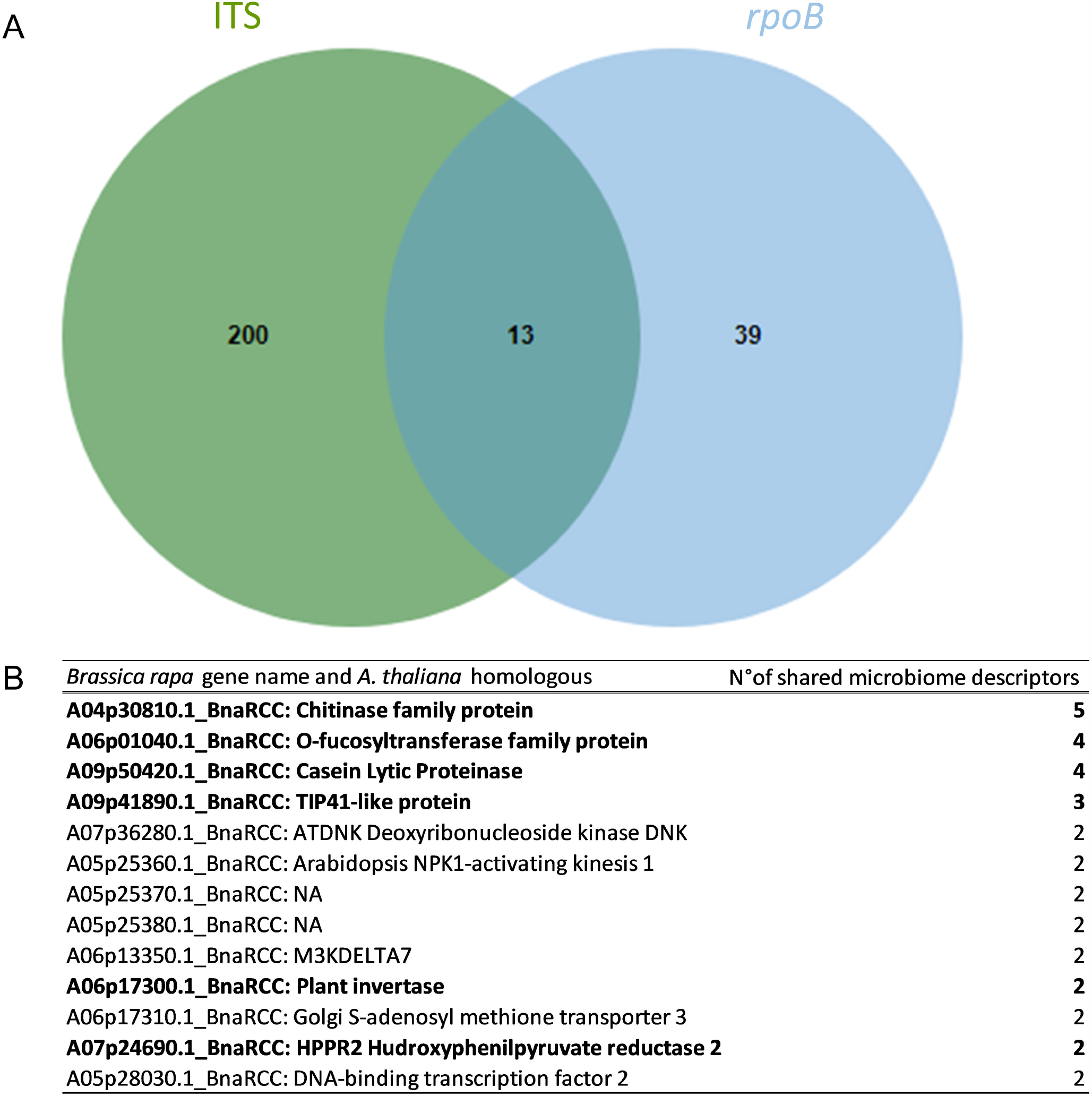
Analysis on the Holobiont Generalist Genes (HGG) shared between fungal and bacterial descriptors. **A**) Venn Diagram showing the 13 genes shared among fungal (in green) and bacterial (in blue) genes that were found to be significantly associated with microbiota descriptors. **B**) Function of the 13 shared genes inferred from *Arabidopsis thaliana*. Genes in bold are those associated with significant XtX values (i.e. genes under natural selection). NA = not attributed, indicates that gene functions were not found for the *A. thaliana* homologous.

**Figure 5.**
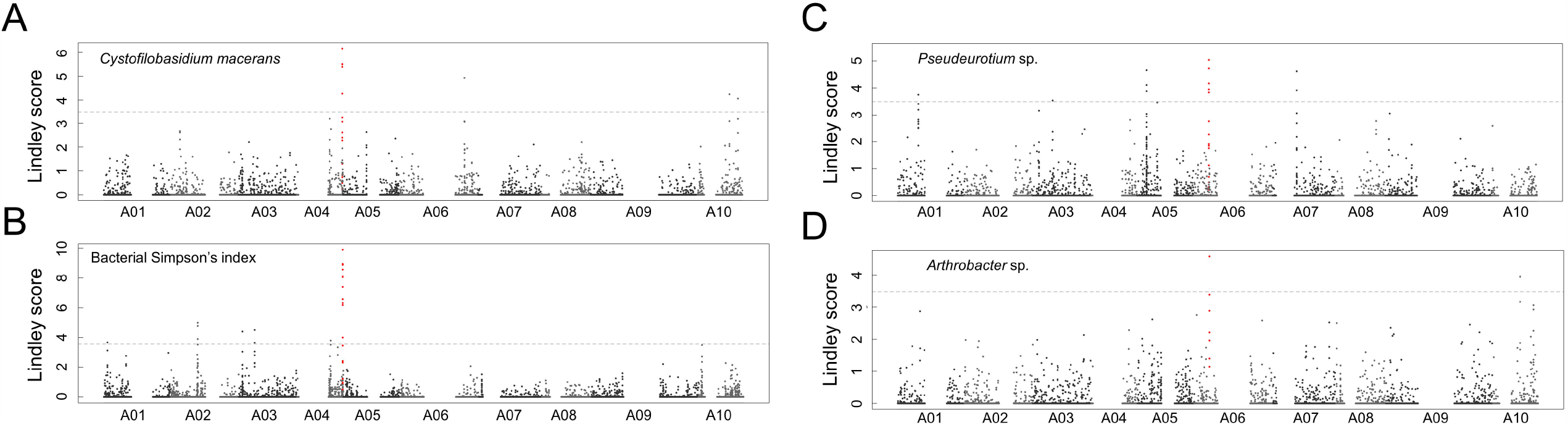
Manhattan plots illustrating Genome-Environmental-Analysis (GEA) for Holobiont Generalist Genes (HGG) found to be associated with both fungal and bacterial descriptors. **A**) Manhattan plot for the fungal OTU *Cystofilobasidium macerans* relative abundance estimated in the rhizosphere compartment in spring. The red peak of association on chromosome 4 falls in the A04p30810.1_BraC13 gene coding for a chitinase. **B**) Manhattan plot for the Simpson’s index estimated on the bacterial dataset in fall. The red peak of association in chromosome four falls in the A04p30810.1_BraC13 gene. **C**) Manhattan plot for the fungal OTU *Pseudeurotium sp*. relative abundance estimated in the root compartment in spring. The red peak of association on chromosome 6 falls in the A06p13350.1_BraC13 gene also called M3KDELTA7. **D**) Manhattan plot for the bacterial OTU *Arthrobacter sp*. relative abundance estimated in the rhizosphere compartment in fall. The red peak of association on chromosome 6 falls in the A06p13350.1_BraC13 gene corresponding to M3KDELTA7. For all Manhattan plots, the *y*-axis corresponds to the values of the Lindley process corresponding to the −log10 (p) of the tuning parameter ξ = 3. The dashed line indicates the significance threshold as described in Bonhomme *et al*., 2019 (16). Blank spaces indicate centromeric positions that were eliminated from the analysis.

Intriguingly, on chromosome 5 three successive genes A05p25360.1_BnaRCC, A05p25370.1_BnaRCC and A05p25380.1_BnaRCC were all associated with bacterial composition (PCoA axis 3) and with the relative abundance of the fungal species *Cladosporium delicatulum* (Fig. 4, Fig. S8, Dataset S11). We identified that the Arabidopsis homologous of the A05p25360.1_BnaRCC gene belongs to a NPK1-activating kinesin-like protein. However, no gene function was found for both A05p25370.1_BnaRCC and A05p25380.1_BnaRCC to evaluate whether the three genes participate to the same metabolic pathways. As gene regulation acts in cis (within one protein), structural interactions between proximally located genes generate genetic interactions of what is called causal genes (24, 25). These causal genes are likely to be detected physically close in GEA as we observed for detected genes associated to the same microbiome variables. In our study, we referred to alleles found to be common between fungal and bacterial descriptors as Holobiont Generalist Genes (HGG). We found that HGG are likely to regulate species richness, composition and abundance of the host microbiome and therefore they are expected to play several roles in modulating plant microbiomes over holobionts evolution. Additionally, our results suggest that HGG not only act at a global microbiome level but they can also physically interact within themselves.

### Holobiont Generalist Genes (HGG) and specific plant-microbiota regulating alleles are under natural selection

A question worth investigating on HGG, consists in identifying whether these microbiota-associated loci have been shaped by natural selection. To answer this question, we additionally tested whether the identified SNPs were enriched in a set of SNPs subjected to adaptive spatial differentiation. This was done by estimating a Bayesian measure of spatial genetic differentiation (XtX) (15). We then intersected the top 0.05% SNPs carrying the highest XtX with the top 0.05% SNPs carrying the highest LD values and we retrieved genes on which these SNPs were located. Among genes shared between the fungal and bacterial descriptors, we found that 6 out 13 genes shaped signatures of selection; i.e., they were associated with high XtX values (Fig 4, Dataset S13). Within the genes with strong signature of selection we found A09p41890.1_BnaRCC (TIP41-like protein *A. thaliana* homologous), A04p30810.1_BnaRCC (chitinase protein *A. thaliana* homologous) and A06p01040.1_BnaRCC (O-fucosyltransferase family protein *A. thaliana* homologous) that are the alleles shared among the highest number of microbiota descriptors (Fig. 4, Fig. 5, Dataset S13). Funding selection signatures on alleles shared among several microbiome descriptors reinforce the hypothesis that strong selective pressures are acting on plant populations to maintain HGG as they are crucial in modulating the host microbiome.

Beside HGG, we found strong signatures of selection on alleles that are specific for fungal or bacterial descriptors. Indeed, we identified strong selection signals on A01p08480.1_BnaRCC (DataSet S13) that is associated with the relative abundance of two fungal species: *Fusicolla* sp. and *Periconia prolifica*, a cobalt-producing fungus with antibacterial activities (26). The *A. thaliana* ortholog of A01p08480.1_BnaRCC is IBI1 (Impaired in BABA-induced disease Immunity) (DataSet S13) and acts as receptor protein activating plant defense through beta-aminobutryric acid (BABA). Interestingly, we found that three *B. rapa* genes (A07p24690.1_BnaRCC, A09p50420.1_BnaRCC and A09p07970.1_BnaRCC) displaying a strong signature of selection were associated with the bacterial biocontrol species *Bacillus simplex* abundance (DataSet S13). This is coherent with the hypothesis that alleles regulating the presence of beneficial plant microbes are likely to be positively selected.

## Discussion

The holobiont concept is now spreading from the scientific and academic environment to the whole society. On June 14th 2023 “The Economist” published an article on how holobionts are shifting the biological meaning of an individual https://www.economist.com/science-and-technology/2023/06/14/the-idea-of-holobionts-represents-a-paradigm-shift-in-biology. But, how much do we know about the evolution of holobionts? What is obvious is that studying the coevolution between wild plants and the associated microbiomes is an important step in capturing the genetic bases influencing the evolutionary trajectories of holobionts. Here, we take advantage of ecologically and genomically differentiated wild populations of the non-model plant species *B. rapa* to study evolutionary traits of holobionts. Wild *B. rapa* populations (AA, 2n=2x=20) are closely related to cultivated forms, (e.g. turnip, leafy and oilseed crops). They are a unique and understudied natural genetic material, providing invaluable insights into the genetic interactions among holobionts in natural habitats. *B. rapa* wild-populations used in our study, were identified by flow cytometry, chloroplast genomic regions sequencing and cytogenetic analysis (13) in the context of the H2020 PRIMA project ‘BrasExplor’.

Not surprisingly, we found that the *B. rapa* fungal and bacterial microbiome is highly variable among plant compartments (roots and rhizospheres) with a highest diversity in the rhizosphere (Fig. 2). This is in accordance with studies conducted on natural *A. thaliana* populations (27) and several crop species such as maize (28) and other cereals (29), or in less-studied plant species like switchgrass (30) or *Agave* (31). As observed for the bacterial leaf and root microbiome in *A. thaliana* (27), we found a seasonal effect on the fungal communities that was mostly significant in the root compartment but not in the rhizosphere (Fig. 2). On perennial crops, similar results were observed on plant leaves (32) with a strong bacterial communities’ variation over seasons. It is therefore evident that studying holobionts over several seasons/years and host compartments is crucial to dissect the genomic architecture regulating the host-microbiome interactions. Also, we observed that microbiome variation is not equivalent among fungal and bacterial communities. In addition, cross-kingdom interactions can influence the evolutionary trajectories of holobionts (33). For example, competition or cooperation among fungal and bacterial communities can define the host microbiome diversity and composition with direct consequences on the evolutionary interactions within a given holobiont (34). Therefore, there is an urgent need to take into consideration both prokaryotic and eukaryotic communities when studying holobionts in their native habitats. A genome-wide-association study on leaf samples of *A. thaliana* accessions showed a highly differentiated host genetic architecture among fungal and bacterial communities (35). Similarly, two studies on *A. thaliana* roots and leaves found almost no overlaps among genes structuring the bacterial and fungal microbiota (34, 36). Nevertheless, the cited studies were conducted in common gardens or experimental fields. The hologenome theory postulates that the evolution of plants and animals is defined by their interactions with the hosted microbiomes in the native ecological context where holobionts coevolve (37). However, studies of holobionts in their native habitats are still underrepresented. To our knowledge, Roux *et al*., 2023 is the only large plant *in situ* survey describing a highly variable genetic architecture between plant compartments (leaves and roots) and seasons (fall and spring) for the bacterial *A. thaliana* microbiome (12). As mentioned by the authors, the next step is to compare fungal and bacterial microbiomes *in situ* to identify common plant genetic loci structuring inter-kingdom interactions. By controlling for plant population structure (15), our approach on wild *B. rapa* populations allowed the identification of the non-negligent number of 13 Holobiont Generalist Genes (HGG) –

i.e. genes regulating the holobionts at several layers among prokaryotes and eukaryotes – shared among fungal and bacterial descriptors. As the bacterial communities were not described in the root compartment, it might be possible that this number is underestimated. Also, underestimation of HGG can be related to important population plant structure within the French and Italian populations. As the Bayesian model used in our study corrects for population structure, true positive association might be removed by decreasing the number of HGG on the overall analysis. Similarly, our study focuses on 26 wild populations, and including more plant genetic diversity could increase the chance to individuate a larger number of HGG. However, “actual” *B. rapa* wild populations, are scarcely distributed and hard to detect. Therefore, increasing the number of plant populations is not as easy as for the worldwide distributed and amply studied *A. thaliana*.

Among the HGG, a large part was associated with fungal or bacterial OTUs reported to be biocontrol agents or Plant-Growth-Promoting (PGP) microbes. For instance, A06p13350.1_BnaRCC was associated with: i) the fungal species *Pseudeurotium* sp. known to be a tomato endophyte with both PGP and biocontrol activity (38) and ii) with the promising biocontrol *Arhtrobacter* bacterial species (Fig. 5) enhancing plant growth and known to suppress mold disease in tomato (39, 40). The A06p13350.1_BnaRCC *A. thaliana* homologous (AT1G18160) belonging to the M3KDELTA7 family was described to be required for the abscisic acid (ABA) signal transduction via SnRK2 (snf1-related protein kinase2s) activation. ABA is known to play a crucial role in plant development as well as in stress tolerance (41). Therefore, in natural plant populations, beneficial fungal and bacterial communities are expected to trigger ABA host responses to confer resistance to both biotic (as pathogens’ attacks) and abiotic stresses. Alleles recognizing beneficial microbes are then likely to be positively selected in natural plant populations. Nevertheless, the A06p13350.1_BnaRCC was not found in our study to be under selection (Fig. 4). On the other hand, the A09p41890.1_BnaRCC gene associated with both *Preussia* fungus and *Bacillus simplex* biocontrol bacterium (20), was enriched for XtX suggesting that this gene is under natural selection in the *B. rapa* populations described here. *Preussia* sp. is producing nitric oxide (NO) and phytohormones like indole-3-acetic acid (IAA) and several gibberellins, and was showed to improve rice growth under greenhouse conditions (42). As for the M3KDELTA7 family, the A06p13350.1_BnaRCC *A. thaliana* ortholog is coding a TIP41 like-protein inducing ABA responses (21). This suggests that HGG not only influence several plant beneficial microbes, but they also trigger generalist plant stress defense pathways. Holobionts are therefore likely to share common biological mechanisms that concretely act to protect the host by positively selecting for microbes that can enhance immune defenses or stimulating host growth. By constituting fungal and bacterial collections from natural plant populations, the next step is validating the role of HGG when interacting with native microbes or when modulating the host defense responses.

## Materials and Methods

### Plant sampling and genome sequencing and analysis

Ecological descriptions of the 26 *B. rapa* populations are described in Falentin *et al*., 2023 (13). For plant genomic characterization, twenty-six populations were collected during February 2020 in the south of Italy (Sicily) and during May 2020 in the north of France (Brittany and Pays de Loire, Supplementary Data Set 1). For plant PoolSeq sequencing, leaves from 30 individuals within each plant population were collected directly *in situ*, and an equal leaf stab was pooled into a sterilized 50ml Falcon tube that was stored at -80°C for 24h prior 2 days of lyophilization. Twenty-five mg of lyophilized leaves were then placed into a 2ml Eppendorf tube and homogenized for 1 min at 35 Hz with the mixer mill RETSCH MM 400. DNA of the lyophilized leaves was extracted with the Qiagen DNeasy Plant Pro Kit by following the manufacturer’s instructions. Integrity, purity and quantity of the DNA was assessed by running 5μl of the extracted DNA on a 1% agarose gel and by Thermo Scientific NanoDrop spectrophotometer. Plant DNA was sequenced at the BGI platform (Hong-Kong, China). For this, 1μg of genomic DNA was randomly fragmented with Covaris. DNA fragments were selected by Agencourt AMPure XP-Medium kit to an average size of 200-400bp. Fragments were end repaired and then 3’ adenylated. Adaptors were ligated to the ends of these 3’ adenylated fragments. Fragments were then amplified by PCR followed by a purification with Agencourt AMPure XP-Medium kit. The double stranded PCR products were heat denatured and circularized by the splint oligosequence. The single strand circle DNA (ssCir DNA) was formatted as the final library. Libraries were qualified by QC. The qualified libraries were sequenced by BGISEQ-500: ssCir DNA molecule formed a DNA nanoball (DNB) containing more than 300 copies through rolling-circle replication. The DNBs were loaded into the patterned nanoarray by using high density DNA nanochip technology. Finally, 100 bp paired end reads were obtained by combinatorial Probe-Anchor Synthesis (cPAS). Raw plant reads were trimmed using Fastp v0.20.1 (44). Clean reads were then mapped to *B. rapa* C1.3 genome assembly (SI Appendix) using bwa mem v0.7.17-r1188 (45) and SNPs calling was conducted with varscan mpileup2cns v2.4.4 https://dkoboldt.github.io/varscan/. The plant allele frequencies matrix was trimmed according to four successive criteria: i) removing SNPs with missing values in more than two populations, ii) only sites with mean depth values (over all populations) greater than or equal to the mean depth minus the standard deviation for depth and less than the mean depth plus the standard deviation for depth were kept iii) sites with a Minor Allele Frequency (MAF) under 0.05 were removed, iv) SNPs falling into centromeric regions were deleted.

### In-situ *microbiota characterization*

Detailed information on the fungal and bacterial microbiota characterization and related bioinformatic and statistical analysis is described in the SI Appendix. The list of fungal and bacterial OTUs obtained after data trimming and the corresponding taxonomy and sequences are available in Supplementary Data Set 14 and 15 respectively.

### Genome-Environmental-Analysis (GEA) to finely map plant loci to microbiota descriptors

GEA analysis was performed on microbiota descriptors (species richness, Shannon’s diversity, PCoA first four axes and abundance of the most prevalent fungal and bacterial OTUs) as described in Roux *et al*., 2023 and Frachon *et al*., 2023 (10, 12). Detailed information about GEA analysis and related scripts are provided in the SI Appendix. Microbiota descriptors used for GEA analysis are available in Supplementary Data Set 4 to 10.

## Data availability

Raw FastQ reads corresponding to the microbiota ITS and *rpoB* characterization were deposited in the Sequence Read Archive (SRA) of NCBI and are available at https://www.ncbi.nlm.nih.gov/sra/PRJNA1049637. Plant raw reads were deposited on NCBI and are available under the BioProject number PRJNA1049637 and the BioSample accession listed in Dataset S16. Reads, genome assembly and gene prediction of *Brassica napus* RCC-S0 are available in the European Nucleotide Archive under the following project PRJEB71115.OTU taxonomic affiliations are reported in Dataset S14 for fungal communities and Dataset S15 for bacterial communities. Scripts used for bioinformatic and statistical analysis are reported in SI Appendix.

## Acknowledgments

This work has been funded by the ERC Starting Grant HoloE^2^Plant (Project N° 101039541) and by the National French INRAE Project ECOSYMB. The RCC-S0 genome was supported by Genoscope and by “Commissariat à l’Energie Atomique et aux Energies Alternatives (CEA)” and “France Génomique (ANR-10-INBS-09-08)”. We kindly thank the H2020 PRIMA BrasExplor consortium for providing us information about plant GPS coordinates. A warm thanks to Fabrice Roux for the exciting discussion on GEA. Many thanks to Gogepp INRAE platform (in particular to Fabrice Legeai) for their technical support in data management and storing. And thanks to all students and colleagues that contribute every day to the ERC HoloE^2^Plant project, without you nothing would be possible.

