## Supplementary material for "Plant genetic bases explaining microbiota diversity shed light into a novel holobiont generalist gene theory": SI Appendix

#### TABLE OF CONTENTS


| | |
|---|---|
| **SUPPLEMENTARY TEXT** | **3** |
| **Supplementary methods** | **3** |
| **Supplementary references** | **7** |
| **SUPPLEMENTARY TABLES** | **9** |
| **SUPPLEMENTARY FIGURES** | **13** |
| **SCRIPTS FOR DATA ANALYSIS** | **22** |

### SUPPLEMENTARY TEXT

#### SUPPLEMENTARY METHODS

##### Brassica napus *RCC-S0 reference genome data production and analysis*

*Illumina PCR-Free library preparation and sequencing.* Genomic DNA of RCC-S0 *Brassica napus* was extracted from 2.5g of fresh plant material as described before (1). For DNA sequencing, 1.5 µg was sonicated to a 100–1500-bp size range using a Covaris E220 sonicator (Covaris, Woburn, MA, USA). The fragments were end-repaired and 3′-adenylated, and Illumina adapters (Nextflex™ PCR Free Barcodes (Perkin Elmer, Waltham, MA, USA)) were added using the Kapa Hyper Prep Kit (Roche, Basel, Switzerland). The ligation products were purified twice with 1X AMPure XP beads (Beckman Coulter Genomics, Danvers, MA, USA). The library was quantified by qPCR using the KAPA Library Quantification Kit for Illumina Libraries (Roche), and its profile was assessed using a High Sensitivity DNA kit on an Agilent Bioanalyzer (Agilent Technologies, Santa Clara, CA, USA). The library was sequenced on an Illumina HiSeq2500 instrument (Illumina, San Diego, CA, USA) using 250 base-length read chemistry in a paired-end mode.

After the Illumina sequencing, an in-house quality control process was applied to the reads that passed the Illumina quality filters. The first step discards low-quality nucleotides (Q < 20) from both ends of the reads. Next, Illumina sequencing adapters and primer sequences were removed from the reads. Then, reads shorter than 30 nucleotides after trimming were discarded. These trimming and removal steps were achieved using in-house-designed software based on the FastX package https://github.com/institut-de-genomique/fastxtend. The last step identifies and discards read pairs that are mapped to the phage phiX genome, using SOAP aligner (2) and the Enterobacteria phage PhiX174 reference sequence (GenBank: NC_001422.1). This processing, described in Alberti (2017) (3), resulted in high-quality data.

*MinION and PromethION library preparation and sequencing.* Libraries were prepared following the protocol provided by Oxford Nanopore Technologies (Oxford Nanopore Technologies Ltd, Oxford, UK) « 1D gDNA long reads without Blue Pippin » with the Ligation Sequencing Kit SQK-LSK108. Genomic DNA (0.8 to 2.5 µg) was repaired using the FFPE DNA repair Mix (New England Biolabs, Ipswich, MA, USA). DNA fragments were then end-prepped with the NEBNext Ultra II End Repair/dA-Tailing Module (NEB) with incubation times of 20 minutes (instead of 5 minutes) at 20°C and 65°C. Sequencing adapters were ligated using the NEB Blunt/TA Ligase Master Mix (NEB) and ligated fragments were purified with AMPure XP beads (Beckmann Coulter, Brea, CA, USA) diluted with a AMPure Dilution Buffer (25mM Tris-HCl (pH 8.0 at 4°C), 400mM NaCl). Beads were pelleted and then washed twice with a 0.6x salty ABB solution (mix of 360 µl ABB from the Ligation Sequencing Kit and 240 µl 5 M NaCl) and once with the TE Buffer provided in the Ligation Sequencing Kit. DNA was eluted in Elution Buffer (incubation at 37°C), then mixed with the Running Buffer with Fuel Mix (RBF) and the Library Loading Beads (LLB) and loaded on MinION R9.4.1 flow cells. Reads were base-called using Albacore version 2.1.10. One library was prepared following the same protocol, but starting with 4µg genomic DNA and a size-selection with a Blue Pippin in order to recover the largest genomic DNA fragments prior to the first step.

Libraries that were loaded on PromethION R9.4.1 flow cells were prepared following the protocol « 1D Genomic DNA by Ligation (SQK-LSK109) – PromethION » with some exceptions : incubation times for the DNA repair and End Prep step were increased to 20 minutes as described above, the ligation temperature was 25°C instead of room temperature, all the incubation times during the beads purification steps were increased from 5 to 10 minutes and the final elution was performed at 37°C. MinION and PromethION reads were respectively basecalled using Albacore 2.1.10 and Guppy 1.4.3. The nanopore long reads were not cleaned and raw reads were used for genome assembly.

*Optical mapping for RCC-S0.* The Direct label and stain (DLS) labelling (using the DLE-1 enzyme) and the Nick Label Repair and Stain (NLRS) labelling (using the BspQI enzyme) protocols were performed according to Bionano Genomics with 750ng and 300ng of DNA respectively. The Chip loadings were performed as recommended by Bionano Genomics.

*Long reads-based genome assembly.* First the genome size of *Brassica napus* RCC-S0 was estimated, using Genomescope (4) and Illumina short-reads, to 860Mb with a low heterozygosity rate. From the raw Nanopore reads, we generated a sample of high-quality reads, representing 30X of coverage, using Filtlong https://github.com/rrwick/Filtlong. The genome of *Brassica napus* RCC-S0 was assembled by giving this subset as input to the SMARTdenovo (5) assembler with default parameters, except '-k 17' and '-c 1' to generate a consensus sequence. The assembly was then polished three times using Pilon[8] (version 1.22) with Illumina reads. The BspQI and DLE-1 preparations were run on a single flow cell each. The generated molecules were assembled to produce optical maps using software provided by Bionano Genomics with the following two options: "add pre-assembly" and "non-haplotype without extend and split" (bionano solve and tools Version: 3.2.2_08222018). The two optical maps and the ONT contigs were subjected to the 2-enzyme hybrid scaffolding pipeline to generate the hybrid scaffolds. BisCoT[9] was used to correct artefactual duplications (negative gaps) introduced during the scaffolding process. A final step of polishing was performed using Pilon and Illumina sequencing data.

*Use of dense genetic maps for scaffold anchoring and psedo-chromosome assembling.* In order to achieve the accurate anchoring of assembled scaffolds to genetic maps and perform pseudochromosome assembly, we used three different segregating populations with a large number of mapped SNP markers:

1) BnaDYDH ('DY'): an established population consisting of 356 doubled haploid (DH) individuals derived from a cross between 'Darmor-bzh' and the cultivar 'Yudal', originating from South-Asia (6)

2) BnaDSDH ('DS'): an established population consisting of 129 doubled haploid (DS) individuals derived from a cross between 'Darmor' and the cultivar 'Samouraï' (6)

3) BnaWYDH ('WY'): a new population consisting of 181 doubled haploid (WY) individuals developed from a cross between 'Westar' and the cultivar 'Yudal', originating from South-Asia (Supplementary Data Set 17).

These populations were genotyped using the Illumina 8K, 20K, and 60K arrays (7) and genetic maps were constructed using CarthaGene software (8). A total of 30576, 11725 and 23368 markers deriving from the Illumina 8K, 20K, and 60K arrays were previously genetically mapped on 'DY', 'DS' and 'WY' maps respectively. The sequence contexts of all single-nucleotide polymorphism markers that were genetically mapped were blasted (Evalue 1E-6, word size 7, reward 1, penalty -3, gapopen 1, gapextend 2, dust yes) against the sequenced genome *Brassica napus* RCC-S0 assembly to validate the quality of our assembly and to help order and orient the scaffolds. After applying a filter of 98% identity and 90% length, 4122, 3219 and 7323 markers ('DY', 'DS' and 'WY' maps respectively) were kept for the final *B. napus* RCC-S0 anchoring (Fig. S9, S10, S11). The genetic and physical positions were discordant for only 425, 59 and 395 markers (7.3% for 'DY', 1.8% for 'DS' and 5.4% fo 'WY') due to an inaccurate position on the genetic map (variation of a few centimorgans in almost all cases).

*Gene prediction.* Repeats in the genome assembly were masked using Tandem Repeat Finder (9) for tandem repeats and RepeatMasker http://repeatmasker.org/ for simple repeats as well as known repeats included in RepBase (10). In addition, known *Brassica* transposable elements (11) were detected using RepeatMasker. Gene prediction was performed using several proteomes: 8 from other genotypes of *B. napus* (12) (Westar, Zs11, QuintaA, Zheyou73, N02127, GanganF73, Tapidor3, and Shengli3), *Arabidopsis thaliana* (UP000006548), the *B. napus* pan-annotation, resistance gene analogues (RGAs) from Brassica (13), and the 2020 annotation of Darmor-bzh (1). Proteomes were aligned on the genome in a 2-step strategy. First, BLAT (14) (version 36 with default parameter) was used to quickly localize corresponding putative regions of these proteins on the genome. The best match and the matches with a score ≥90% of the best match score have been retained. Second, alignments were refined using Genewise[18] (version 2.2.0 with default parameters), which is more accurate for detecting intron/exon boundaries. Alignments were kept if >75% of the length of the protein was aligned on the genome.

All the protein alignments were combined using Gmove (15), which is an easy-to-use predictor with no need of a pre-calibration step. Briefly, putative exons and introns, extracted from alignments, were used to build a graph, where nodes and edges represent, respectively, exons and introns. Gmove extracts all paths from the graph and searches open reading frames that are consistent with the protein evidence. Finally, we decided to exclude genes having both >50% of untranslated regions and a CDS length smaller than 300bp. Following this pipeline, we predicted 106,447 genes with 4.41 exons per gene on average.

***Microbiota characterization and bioinformatic analysis***

*Sampling and generation of ITS1 and* rpoB *amplicons.* For *B. rapa* microbiota characterization, roots and rhizosphere samples were collected and separated *in situ* and stored in a cold box at 4°C during travelling. In total, we collected 6 plants for each population during two seasons: i) spring, corresponding to February 2020 in Sicily and May 2020 in France, and ii) fall, corresponding to December 2020 in Italy and January 2021 in France. Each root rhizosphere sample was placed in a 2ml Eppendorf tube containing 3 mm metallic beads and it was lyophilized for 2 days. After lyophilization, all samples were then homogenized into individual tubes for 1 min at 35 Hz with the mixer mill RETSCH MM 400. The obtained powders were placed in 1.2 ml S-block 96-well plates by using a metallic spoon. At each sample the spoon was cleaned with pure alcohol. DNA was extracted by using the Qiagen DNeasy 96 PowerSoil Pro Kit (reference: 47017) by following kit instructions. The extracted DNA was then quantified with the NanoDrop spectrophotometer. All DNAs were adjusted to 20ng/µl prior amplicon generation. Specifically, bacterial communities were characterized through amplification of a fraction of the *rpoB* gene with primers and conditions described before (16) by using internal TAGS for multiplexing as described in Bartoli *et al*., 2018 (17). Fungal communities were characterized by amplifying a portion of the ITS1 with a protocol developed in this study and adapted to *B. rapa*. For this the following primers have been used: ITS1Fngs 5' - GGTCATTTAGAGGAAGTAA - 3' and ITS2ngs 5' - TTYRCKRCGTTCTTCATCG - 3'. PCR program consisted of: initial denaturation at 95°C for 5 min, 35 cycles at 95°C for 5 sec, 58°C for 30 sec, 72°C for 30 sec and a final elongation at 72°C for 10 min. All PCRs for both markers were performed in a final volume of 50 µl with the MTP Taq DNA Polymerase (Sigma-Aldrich, reference: D7442). PCR amplicons were then purified by using Agencourt®AMPure® magnetic beads following the manufacturer’s instructions. Purified amplicons were quantified by spectrometry and appropriately diluted to obtain an equimolar concentration. Two microliters of equimolar PCR-purified products were used for a second PCR containing the 288 Illumina indexes. The second PCR amplicons were then purified and quantified as described above to obtain a unique equimolar pool that was quantified by real-time quantitative reverse transcriptase-polymerase chain reaction and sequenced with Illumina MiSeq 2 × 250 v3 (Illumina Inc., San Diego, CA, USA) in the GeT-PlaGe Platform (Toulouse, France).

Bioinformatics analysis and data filtering

Raw reads were demultiplexed by considering the Illumina indexes and the three internal TAGS with the Flexbar software (18) available at https://github.com/seqan/flexbar. For the fungal communities, the total number of reads and OTUs before data trimming was 11, 345, 800 and 14, 142 respectively. The average number of complete pairs sequences per sample was 18, 182 (SD = 5, 022). For the bacterial data set, the total number of reads and OTUs before data trimming was 3, 631, 685 and 6, 838 respectively. The average number of complete pairs per sample was 1, 125 (SD = 963). For both *rpoB* and ITS1 datasets we removed samples with less than 1,000 reads to keep only samples represented by a correct number of reads. This led to the elimination of the entire *rpoB* roots dataset. Taxonomic affiliation was performed by using a Bayesian classifier implemented in the classify.seqs command of MOTHUR (19) against the UNITE ITS database v 8.2 (fungi-only) and the *rpoB* database available at FROGS https://frogs.toulouse.inra.fr/. Clustering of sequences into OTUs was performed with Swarm (20) https://github.com/torognes/swarm by using a clustering threshold (d) = 1. Only OTUs with a minimum of five sequences across all samples were kept. After data filtering, for the ITS we obtained 602 samples and 1227 OTUs and for *rpoB* 465 samples and 1237 OTUs. Script for bioinformatic analysis are provided below in the script section.

Analysis of microbiota descriptors

The final OTU matrices obtained after data filtering were used to estimate microbiota α-diversity descriptors with summary.single function in Mothur. Microbiota composition (β-diversity) was inferred and analyzed with two complementary approaches. Firstly, a Hellinger transformation was performed by using the vegan R package and the relative Hellinger distance was inferred with the decostand command in the vegan R package prior β-diversity analyses. For both fungal and bacterial datasets, the Hellinger distance matrices were reduced by running a Principal Coordinates Analysis (PCoA) with the ape R package. Secondly, we applied constrained ordination (tb-RDA) on the OTU matrices. For this, we first estimated the relative abundance of each bacterial and fungal OTU, and selected OTUs with a frequency ≥ 1. The trimmed OTU count table was then normalized and tb-RDA inferred with the rda function in vegan R package. To control for plant compartment, tb-RDA was independently run on rhizosphere and root samples. Natural variation for microbiota descriptors for both fungal and bacterial communities was used to run liner mixed models with the lmer function in the lsmeans R package and lm4 by considering the plant compartment and genotype as fixed factors in the model. *P*-values obtained by linear-mixed analysis were corrected for the FDR. As both fungal and bacterial communities were dependent to the plant compartment, linear mixed models were inferred separately on rhizosphere and root samples.

Box-plots on microbiota diversity descriptors were performed in the R environment by using general coding (scripts available in the section below). To visualize the season effect on α-diversity descriptors we plot Shannon and richness data for both fungal and bacterial communities in a Finlay Wilkinson by using ggplot2. To test the impact of plant population on the variation of fungal and bacterial microbiota descriptors, we performed broad-sense heritability ($H^2$) by lmer model from the lme4 package, and by considering the plant population as a random effect. $H^2$ was then estimated by dividing the variance obtained for the plant genotype effect divided by the sum of the genotype variance on the residual variance that was divided by the number of replicates in the dataset. As $H^2$ showed a high effect of the plant population on microbiota descriptors, we estimated Best-Linear-Unbiased-Predictions (BLUPs) by running a linear model with lmer and by considering the plant population as a fixed effect. BLUPs were estimated for species richness, Shannon’s index, the first four PCoA axis, and the most relevant, but variable among the plant populations, fungal and bacterial OTUs, which are present in at least ten samples. As we found a seasonal and plant compartment effect, BLUPs were individually estimated for each combination within the fungal and bacterial descriptors: R-FA (roots in fall), R-SP (roots in spring), RH-FA (rhizosphere in fall) and RH-SP (rhizosphere in spring). Script for statistical analysis and figure’s construction are provided below in the script section.

Genome-Environmental-Analysis (GEA)

GEA analysis was run on fungal and bacterial BLUPs estimated for each microbiome descriptor and for the four-plant compartment/seasonal groups described above. For this the SNPs trimmed matrix and BLUPs for microbiome descriptors, were implemented in the BayPass software (21) that is estimating for each SNPs a Bayesian Factor ($BF_{is}$) and associated regression (Beta_is). We used the same parameters described in Roux *et al*., 2023 (22). To increase the speed of the analysis, for each plant compartment/seasonal groups we divided the datasets in several subsets in R environment using the pooldata2genobaypass from the poolfstat package (Gautier 2022) used to parallelized the GEA. To define the highly significant associations between plant SNPs and microbiome descriptors; the GEA outputs obtained with the core model implemented in BayPass were integrated in the Local Score (LS) approach (23) allowing the detection of significant genomic regions by accumulating statistical signals from contiguous genetic markers such as SNPs. The LS approach increases the chances to detect QTLs with small effects. For all the plant genomic regions associated with high LS factors we performed Manhattan plots in the R environment with the script provided below. We then retrieved all candidate genes located in the QTLs associated with significant LS factors on the reference *B. rapa* C1.3 genome. As few information are provided on the *B. rapa* gene functions, we searched for *A. thaliana* orthologues and related function on the TAIR10 database available at https://www.arabidopsis.org/. For all the genes falling in significant QTLs and for which we retrieved *A. thaliana* homologous, we intersected the plant compartment/seasonal groups within each fungal and bacterial dataset to identify common genes among plant compartments and seasons. This was made by using the jvenn software available at https://jvenn.toulouse.inrae.fr/app/index.html. To identify the Holobiont Generalist Genes (i.e. genes shared among fungal and bacterial descriptors) we intersected all genes found to be associated to fungal and bacterial descriptors without considering plant compartment/seasonal groups. Finally, to identify regions that are under selection, for each plant compartment/seasonal group, we estimated a measure of spatial genetic differentiation (XtX) among the populations. The XtX is implemented in the BayPAss software (21). We then retrieved plant regions enriched with significant XtX and intersected with the regions showing significant LS values and that were falling into *B. rapa* genes.

#### Supplementary Tables

**Table S1**. Analysis of Deviance Table (Type II tests). General-Linear-Model (GLM) on fungal communities was run on the complete dataset to test the compartment effect. Degree of freedom = 1. *P*-values were corrected for FDR. Bold format indicates significant values after FDR correction.

| | Fungi | |
|---|---|---|
| | $\chi^2$ | *P* |
| Richness | 468.73 | **2.20E-16** |
| Shannon's index | 678.19 | **2.20E-16** |
| PCoA1 | 0.03 | 8.73E-01 |
| PCoA2 | 0.01 | 9.39E-01 |

**Table S2**. Analysis of Deviance Table (Type II tests). General-Linear-Model (GLM) on fungal communities was run separately on roots and rhizosphere compartments to test the season effect. Degree of freedom = 1. *P*-values were corrected for FDR. Bold format indicates significant values after FDR correction.

| | Fungi (R) | | Fungi (RH) | |
|---|---|---|---|---|
| | $\chi^2$ | *P* | $\chi^2$ | *P* |
| Richness | 44.03 | **3.23E-11** | 0.22 | 6.37E-01 |
| Shannon's index | 54.66 | **1.43E-13** | 2.44 | 1.18E-01 |
| PCoA1 | 0.08 | 7.77E-01 | 0.00 | 1.00E+00 |
| PCoA2 | 0.02 | 9.00E-01 | 0.00 | 1.00E+00 |

**Table S3**. Transformation-based Canonical Redundancy Analysis (tb-RDA) on fungal communities showing the effect of plant population on fungal composition between seasons. Bold values reflect statistical significance.

| | Fungi (R-FA) | | | Fungi (R-SP) | | | Fungi (RH-AT) | | | Fungi (RH-SP) | | |
|---|---|---|---|---|---|---|---|---|---|---|---|---|
| | Variance | *F* | *P* | Variance | *F* | *P* | Variance | *F* | *P* | Variance | *F* | *P* |
| Genotype | 0,12 | 1,37 | **1,00E-03** | 0,15 | 1,21 | **1,00E-03** | 0,15 | 1,34 | **1,00E-03** | 0,15 | 1,29 | **1,00E-03** |
| Residual | 0,42 | | | 0,57 | | | 0,52 | | | 0,56 | | |

**Table S4.** Analysis of Deviance Table (Type II tests). General-Linear-Model (GLM) on fungal and bacterial community's descriptors was run on compartment (Roots = R, Rhizosphere = RH) and season datasets to test the effect of plant populations. Degree of freedom = 25. *P*-values were corrected for FDR. Bold format indicates significant values after FDR correction.

| | Fungi (R) Fall | | Fungi (R) Spring | | Fungi (RH) Fall | | Fungi (RH) Spring | | Bacteria (RH) Fall | | Bacteria (RH) Spring | |
|---|---|---|---|---|---|---|---|---|---|---|---|---|
| | $X^2$ | *P* | $X^2$ | *P* | $X^2$ | *P* | $X^2$ | *P* | $X^2$ | *P* | $X^2$ | *P* |
| Richness | 70.80 | **2.96E-06** | 57.10 | **2.61E-04** | 33.70 | 1.15E-01 | 66.40 | **1.29E-05** | **1.69E+02** | **2.20E-16** | 56.90 | **2.76E-04** |
| Shannon's index | 85.70 | **1.44E-08** | 69.50 | **4.57E-06** | 27.60 | 3.26E-01 | 67.60 | **8.81E-06** | 8.73E+01 | **7.99E-09** | 38.18 | **4.44E-02** |
| PCoA1 | 54.80 | **5.20E-04** | 61.60 | **6.30E-05** | 404.50 | **2.20E-16** | 192.60 | **2.20E-16** | 4.01E+01 | **2.84E-02** | 67.91 | **7.82E-06** |
| PCoA2 | 125.90 | **2.03E-15** | 149.00 | **2.20E-16** | 103.90 | **1.37E-11** | 148.30 | **2.20E-16** | 6.65E+01 | **1.26E-05** | 37.02 | 5.75E-02 |

**Table S5.** Analysis of Deviance Table (Type II tests). General-Linear-Model (GLM) on bacterial communities was run separately the rhizosphere compartment to test the season effect. Degree of freedom = 1. *P*-values were corrected for FDR.

| | Bacteria (RH) | |
|---|---|---|
| | $\chi 2$ | *P* |
| Richness | 1,15 | 0,28 |
| Shannon's index | 0,31 | 0,58 |
| PCoA1 | 0,00 | 0,97 |
| PCoA2 | 0,51 | 0,48 |

**Table S6**. Transformation-based Canonical Redundancy Analysis (tb-RDA) on bacterial communities showing the effect of plant population on bacterial composition along the seasons. Degree of freedom = 25. Bold values reflect statistical significance.

| | Bacteria (RH-SP) | | | Bacteria (RH-FA) | | |
|---|---|---|---|---|---|---|
| | Variance | *F* | *P* | Variance | *F* | *P* |
| Genotype | 0,25561 | 1,4463 | **1,00E-03** | 0,26314 | 1,4074 | **1,00E-03** |
| Residual | 0,65038 | | | 0,64318 | | |

**Table S7.** Broad-sense heritability ($H^2$) estimated on both fungal and bacterial descriptors over the two seasons. Bold values represent significant $H^2$.

| Heritability for fungal descriptors | | | | | |
|---|---|---|---|---|---|
| Compartment | Season | Shannon's index | Richness | PCoA1 | PCoA2 |
| R | FA | **0,72** | **0,66** | **0,56** | **0,81** |
| R | SP | **0,65** | **0,58** | **0,61** | **0,84** |
| RH | FA | 0,11 | 0,27 | **0,94** | **0,77** |
| RH | SP | **0,64** | **0,63** | **0,87** | **0,84** |

| Heritability for bacterial descriptors | | | | | |
|---|---|---|---|---|---|
| Compartment | Season | Shannon's index | Richness | PCoA1 | PCoA2 |
| RH | FA | **0,78** | **0,88** | **0,45** | **0,71** |
| RH | SP | **0,47** | **0,66** | **0,74** | **0,44** |

### Supplementary Figures

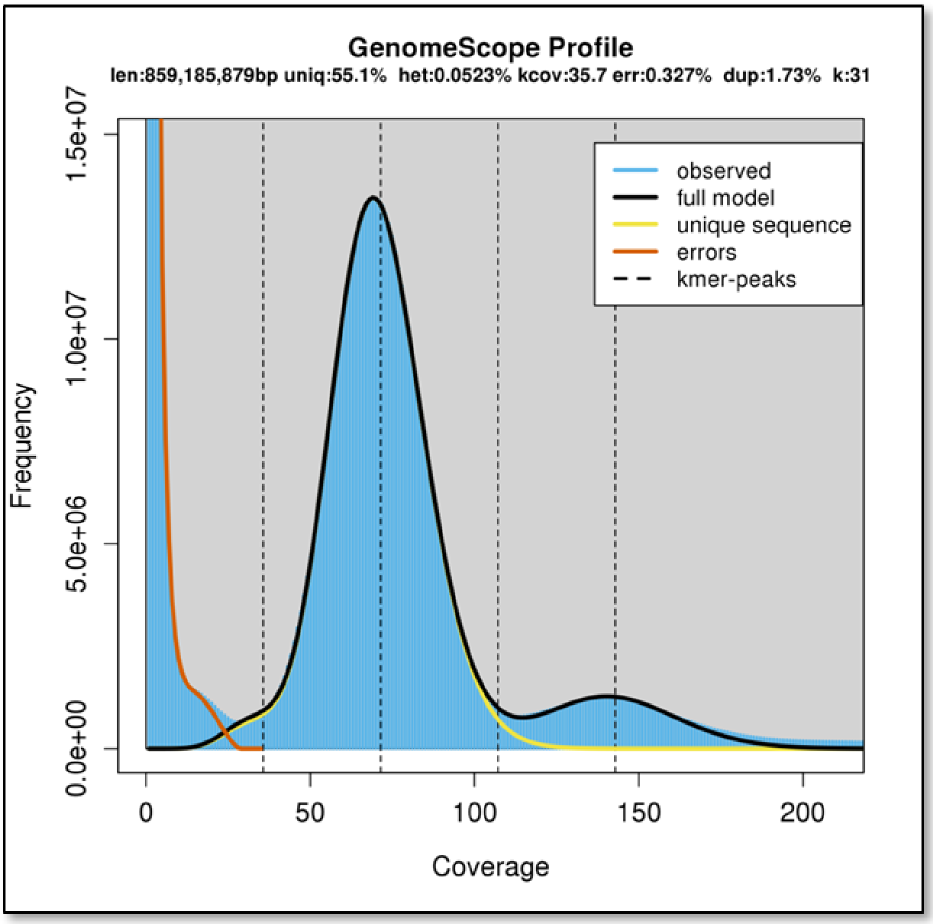


**Figure S1:** Genomescope profile from short-reads of *Brassica napus* RCC-S0.

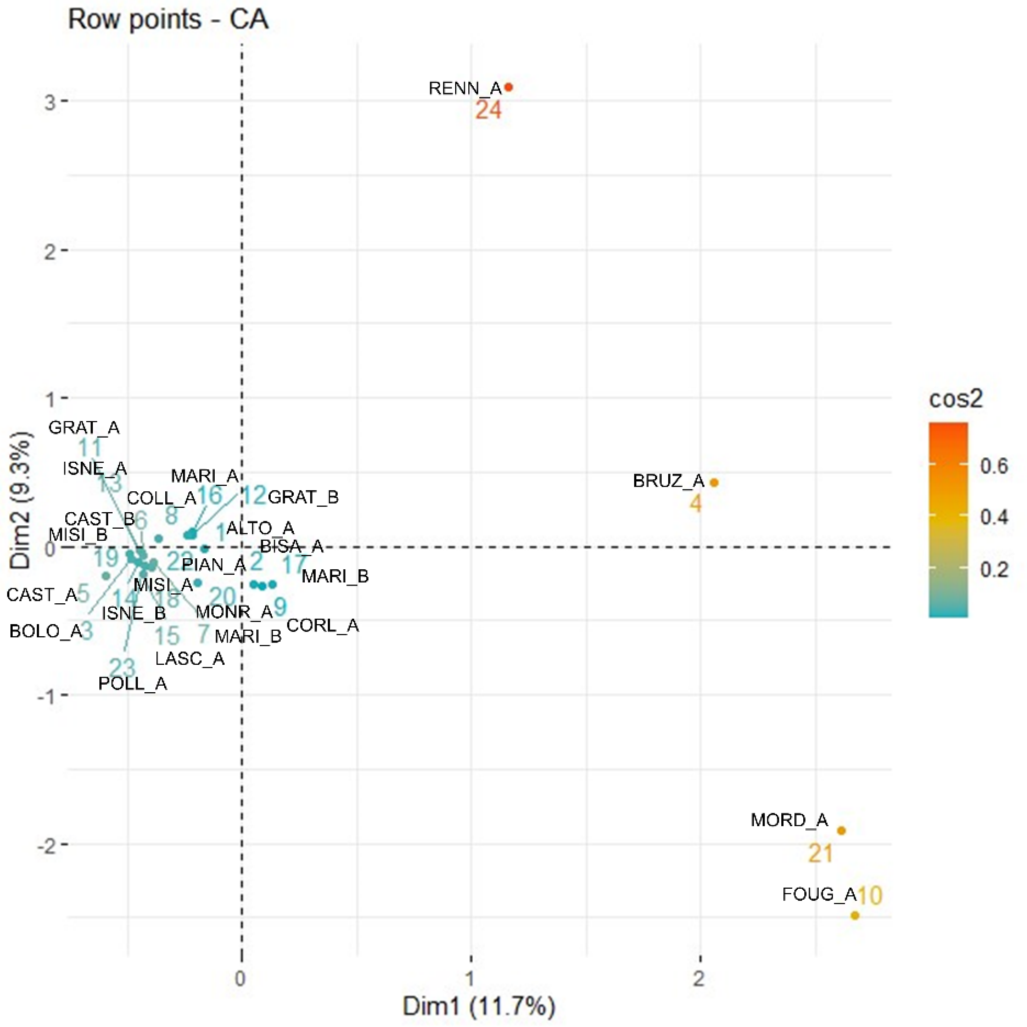


**Figure S2.** Correspondence Analysis (CA) based on plant communities associated with *Brassica rapa* wild populations. Two French populations were not characterized for plant communities' assembling (APIG_A and BOEL_A). CA was inferred on a presence/absence matrix. Red to blue gradient indicates the significance of the population differentiation in terms of plant communities 'assembling (dark red = highly significant, light blue = not significant differentiation among the populations. French populations indicated under the name of FOUG_A, MORD_A, BRUZ_A and RENN_A were highly differentiated in term of associated-plant communities when comparing with the Italian populations.

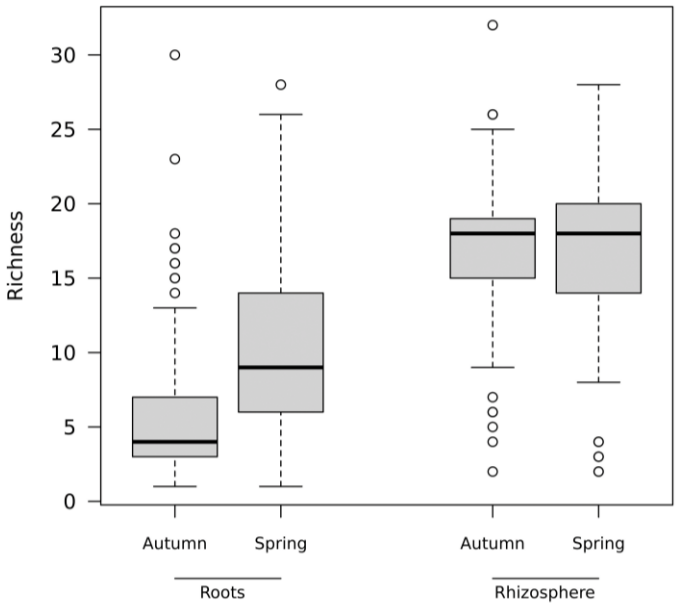


**Figure S3**. Box plot on the species richness estimated on the ITS dataset and showing the effect of plant compartment on the $\alpha$-diversity of fungal communities.

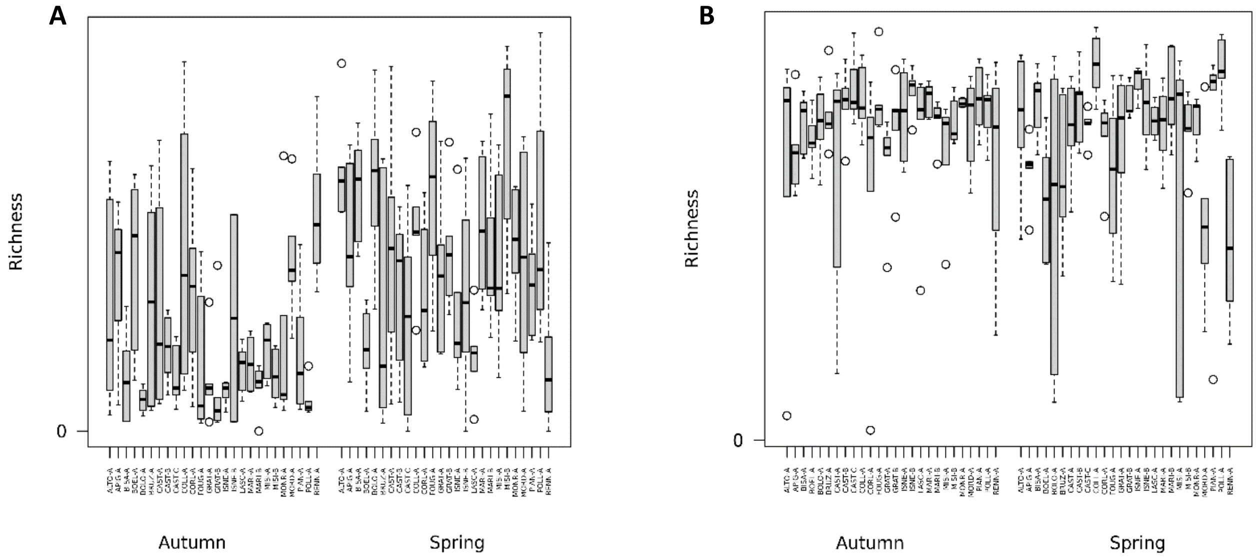


**Figure S4**. Box plot on the species richness estimated on the ITS dataset and showing the effect of plant population on the $\alpha$-diversity of fungal communities in root (A) and rhizosphere (B) compartment.

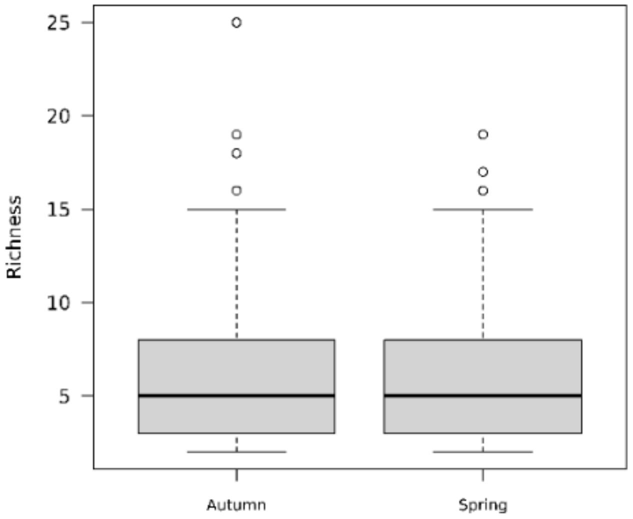


**Figure S5.** Box plot on the species richness estimated on the *rpoB* dataset and showing that season did not affect bacterial diversity in the rhizosphere compartment.

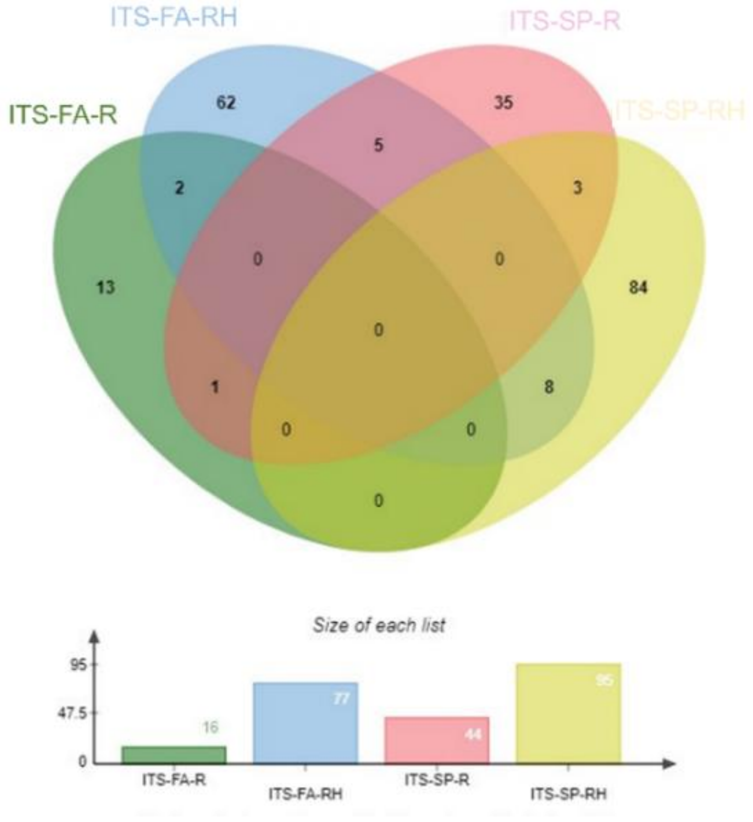

**Figure S6.** Venn Diagram on genes falling on significant Local Score SNPs for fungal communities. FA = fall, SP = spring, R = roots and RH = rhizosphere.

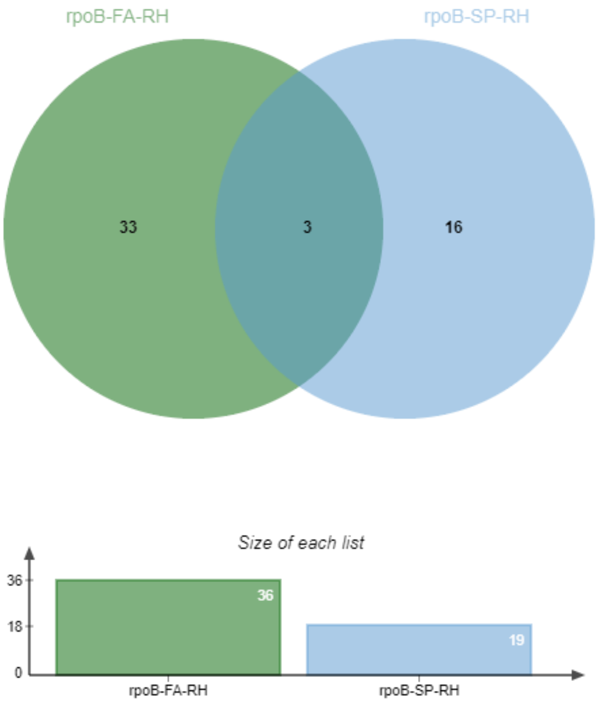


**Figure S7.** Venn Diagram on genes falling on significant local score SNPs for bacterial communities. FA = fall, SP = spring and RH = rhizosphere.

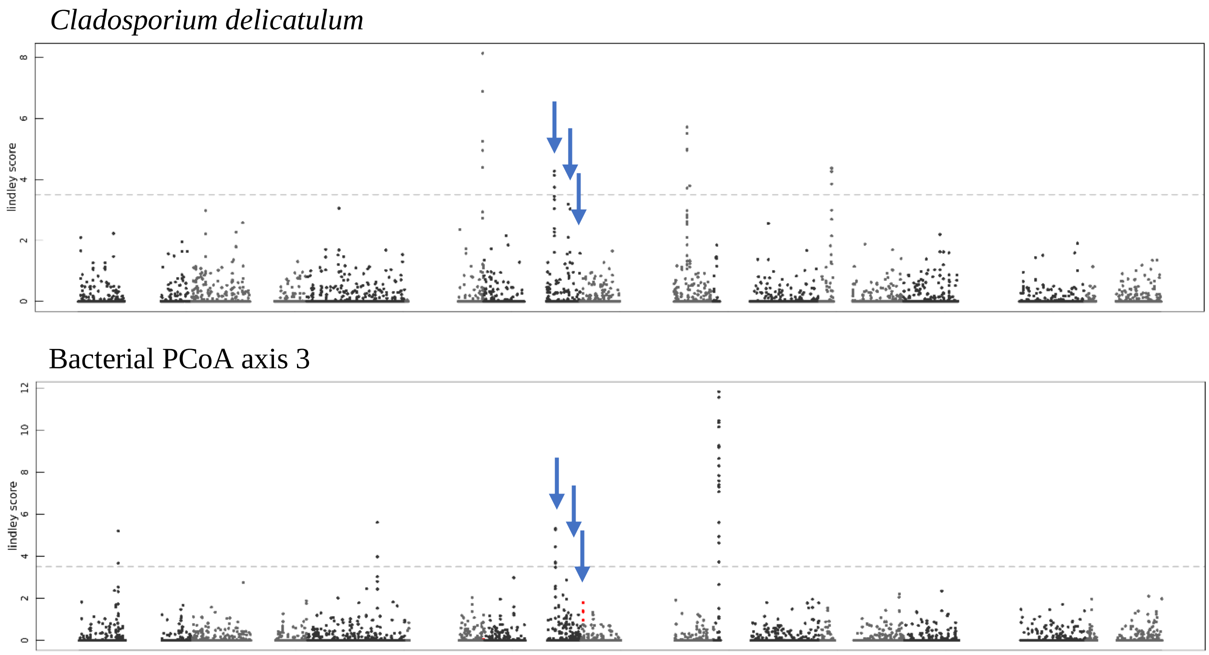


**Figure S8.** Manhattan plots illustrating the Genome-Environmental-Analysis (GEA) for holobiont generalist genes (HGG) found to be physically close**.** Top panel) Manhattan plot for the fungal OTU *Cladosporium delicatulum* relative abundance estimated in the root compartment during spring. Bottom panel) Manhattan plot for the bacterial PCoA axis 3 estimated in the rhizosphere compartment during fall. Gray gradient indicates 10 *Brassica rapa* chromosomes. Blue arrows indicate the three physically close genes A05p25360.1_BnaRCC, A05p25370.1_BnaRCC and A05p25380.1_BnaRCC. Blank ranges indicate centromeric regions.

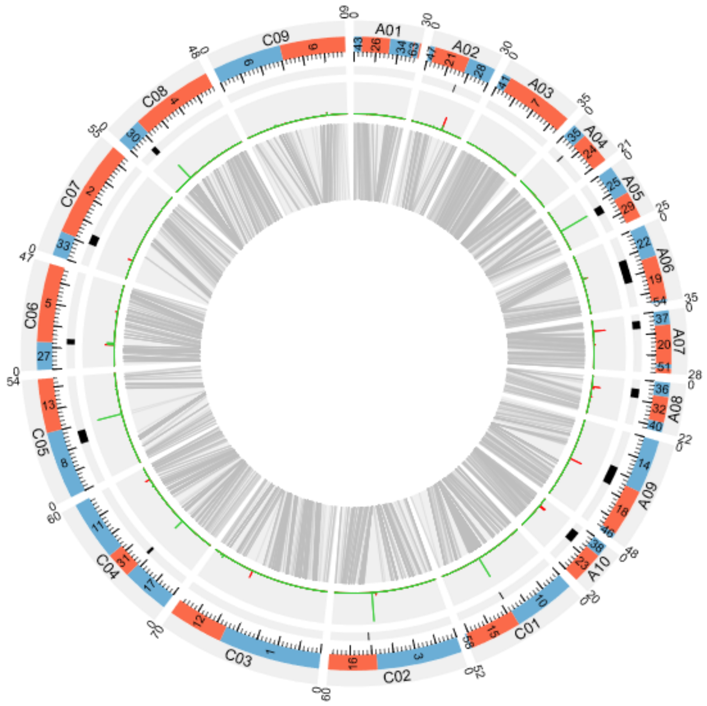


**Figure S9**. Genome overview of the 19 chromosomes of *B. napus* RCC-S0 after scaffolds anchoring on ‘DY’ genetic map. Tracks from outside to inside: chromosomes names (A01 to C09); ordered and oriented scaffolds names; localization of centromere (black squares) from flanking markers identified by Mason *et al*. (24); density of repeated pericentromere-specific sequences (green and red) of Brassica allowing more precise localization of centromeres; genetic and physical links between ‘DY’ genetic map and physical markers on corresponding scaffolds.

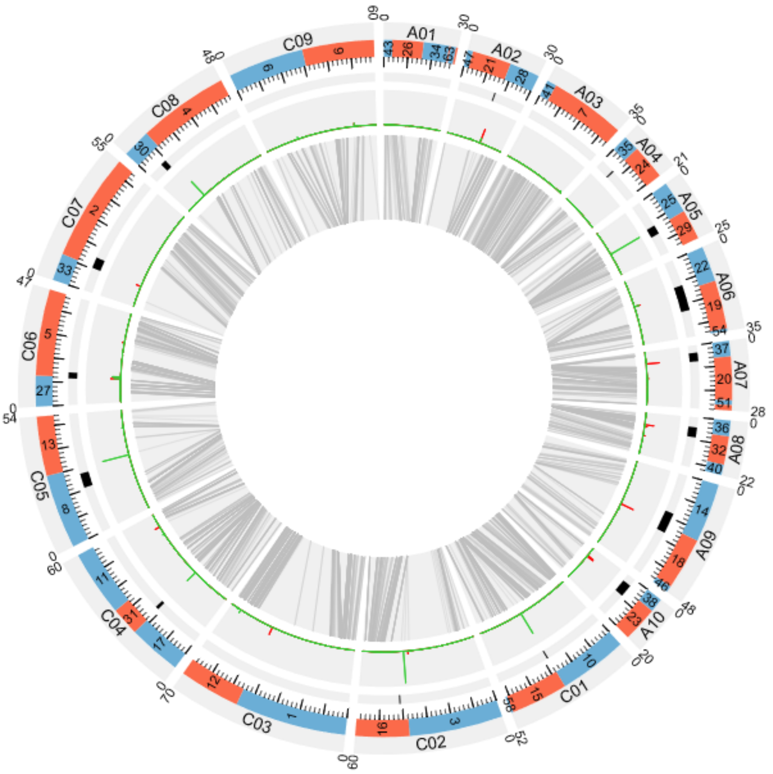


**Figure S10**. Genome overview of the 19 chromosomes of *B. napus* RCC-S0 after scaffolds anchoring on 'DS' genetic map. Tracks from outside to inside: chromosomes names (A01 to C09); ordered and oriented scaffolds names; localization of centromere (black squares) from flanking markers identified by Mason *et al*. (24); density of repeated pericentromere-specific sequences (green and red) of Brassica allowing more precise localization of centromeres; genetic and physical links between 'DS' genetic map and physical markers on corresponding scaffolds.

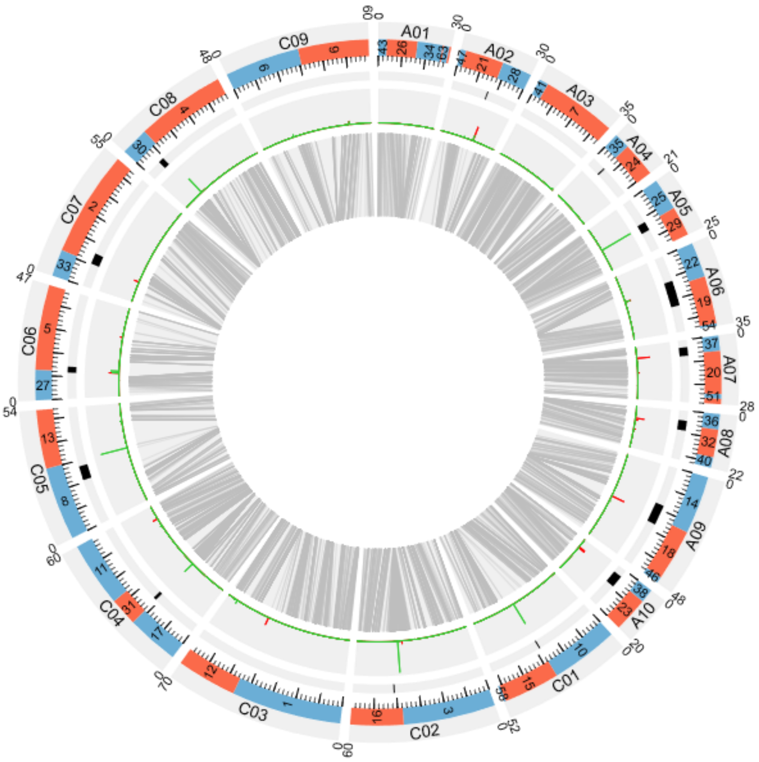


**Figure S11**. Genome overview of the 19 chromosomes of *B. napus* RCC-S0 after scaffolds anchoring on ‘WY’ genetic map. Tracks from outside to inside: chromosomes names (A01 to C09); ordered and oriented scaffolds names; localization of centromere (black squares) from flanking markers identified by Mason *et al*. (24); density of repeated pericentromere-specific sequences (green and red) of Brassica allowing more precise localization of centromeres; genetic and physical links between ‘WY’ genetic map and physical markers on corresponding scaffolds.

**SCRIPTS FOR DATA ANALYSIS**

**The following scripts have been used for the statistical analysis performed in R to calculate microbiota descriptors and microbiota representation. Only alpha-diversity descriptors were estimated in mothur**

**# codes required to install the needed packages**

```
if (!require("BiocManager", quietly = TRUE))
    install.packages("BiocManager")


BiocManager::install("mixOmics")


install.packages("RVAideMemoire")
install.packages("stringi")
install.packages("qpcR")
library(RVAideMemoire)
library(qpcR)
library(stringi)
library(stringr)
library(dplyr)
library(tidyr)
library(lme4)
library(dplyr)
library(lsmeans)
library(nlme)
library(ape)
library(vegan)
library(car)
library(ggplot2)
library(RColorBrewer)
library(data.table)
library(tidyverse)
```

**#Load the statistic table provided by Mothur to keep only samples with up to 1000 reads.**

```
mothur_stats <- read.table(text=paste0(head(readLines("combo.trim.contigs.good.unique.agc.unique_list.0.03.rep.count_table.min5.stats"), -3), collapse="\n"))
```

**#1000 reads filtering**

```
samples_to_keep <- subset(mothur_stats[-1,], mothur_stats[-1,]$V2 > 1000)

rownames(samples_to_keep) <- NULL


mothur_counts <- read.table("combo.trim.contigs.good.unique.agc.unique_list.shared.min5", header=T)

mothur_counts_samples <- subset(mothur_counts, mothur_counts$Group %in% samples_to_keep$V1)

write.table(mothur_counts_samples, file="shared_1000.txt", col.names=TRUE, row.names = FALSE, quote=FALSE, sep='\t')
```

**# run the perl script to select the most abundant OTUs 1% of cutoff (script from Matthieu Barret)**

```
perl elim_LowFreqOTU_from_shared_make_database.pl -f 1 -s shared_1000.txt -o shared_1000_abd1

awk '{print NF}' shared_1000_abd1 | sort -nu | tail -n 1

shared_1000_abd1=read.table("shared_1000_abd1", header=T)

shared_1000_abd1 <- shared_1000_abd1[,-c(1,3)]

shared_1000_abd1$Group[314] <- str_sub(shared_1000_abd1$Group[314], end = -2)

stri_sub(shared_1000_abd1$Group, -1, 1) <- "_"

write.table(shared_1000_abd1, file="rpoB_OTU_table.txt", col.names=TRUE, row.names = TRUE, quote=FALSE)
```

**#Alpha-diversity indices are calculated in Mothur with the following formula**

```
mothur "#summary.single(shared=shared_1000_abd1_fmt, calc=invsimpson-npshannon-chao-sobs)"
```

**#Beta-diversity estimation**

**##Creation of datasets for each season and plant compartment**

```
shared_1000_abd1$combo <- paste(str_split_fixed(shared_1000_abd1$Group, '_', 5)[,4], str_split_fixed(shared_1000_abd1$Group, '_', 5)[,3], sep= "_")

shared_1000_abd1_RH_AT <- as.data.frame(split(shared_1000_abd1, f=shared_1000_abd1$combo)["RH_AT"])

shared_1000_abd1_RH_SP <- as.data.frame(split(shared_1000_abd1, f=shared_1000_abd1$combo)["RH_SP"])
```

```
shared_1000_abd1_R_AT <- as.data.frame(split(shared_1000_abd1, f=shared_1000_abd1$combo)["R_AT"])

shared_1000_abd1_R_SP <- as.data.frame(split(shared_1000_abd1, f=shared_1000_abd1$combo)["R_SP"])


shared_1000_abd1_RH_AT <- shared_1000_abd1_RH_AT[-ncol(shared_1000_abd1_RH_AT)]

shared_1000_abd1_RH_SP <- shared_1000_abd1_RH_SP[-ncol(shared_1000_abd1_RH_SP)]

shared_1000_abd1_R_AT <- shared_1000_abd1_R_AT[-ncol(shared_1000_abd1_R_AT)]

shared_1000_abd1_R_SP <- shared_1000_abd1_R_SP[-ncol(shared_1000_abd1_R_SP)]


colnames(shared_1000_abd1_RH_AT) <- str_sub(colnames(shared_1000_abd1_RH_AT), start = 7)

colnames(shared_1000_abd1_RH_SP) <- str_sub(colnames(shared_1000_abd1_RH_SP), start = 7)

colnames(shared_1000_abd1_R_AT) <- str_sub(colnames(shared_1000_abd1_R_AT), start = 6)

colnames(shared_1000_abd1_R_SP) <- str_sub(colnames(shared_1000_abd1_R_SP), start = 6)
```

**## Hellinger transformation**

```
P1=shared_1000_abd1_R_AT[,c(4:ncol(shared_1000_abd1_R_AT))]

row.names(P1)=shared_1000_abd1_R_AT$Group

A.hel=decostand(P1, "hellinger", na.rm=TRUE)

write.table(A.hel, file="hellinger_rpoB_AT_R.tab", col.names=TRUE, row.names = TRUE, quote=FALSE)

A.hel.D=dist(A.hel)

A.hel.D_AT_R=as.matrix(A.hel.D)

write.table(A.hel.D_AT_R, file="hellinger_distance_matrix_rpoB_AT_R.txt", col.names=TRUE, row.names = TRUE, quote=FALSE)


P2=shared_1000_abd1_R_SP[,c(4:ncol(shared_1000_abd1_R_SP))]

row.names(P2)=shared_1000_abd1_R_SP$Group

A.hel=decostand(P2, "hellinger", na.rm=TRUE)

write.table(A.hel, file="hellinger_rpoB_SP_R.tab", col.names=TRUE, row.names = TRUE, quote=FALSE)

A.hel.D=dist(A.hel)

A.hel.D_SP_R=as.matrix(A.hel.D)
```

```
write.table(A.hel.D_SP_R, file="hellinger_distance_matrix_rpoB_SP_R.txt", col.names=TRUE, row.names = TRUE, quote=FALSE)


P3=shared_1000_abd1_RH_SP[,c(4:ncol(shared_1000_abd1_RH_SP))]

row.names(P3)=shared_1000_abd1_RH_SP$Group

A.hel=decostand(P3, "hellinger", na.rm=TRUE)

write.table(A.hel, file="hellinger_rpoB_SP_RH.tab", col.names=TRUE, row.names = TRUE, quote=FALSE)

A.hel.D=dist(A.hel)

A.hel.D_SP_RH=as.matrix(A.hel.D)

write.table(A.hel.D_SP_RH, file="hellinger_distance_matrix_rpoB_SP_RH.txt", col.names=TRUE, row.names = TRUE, quote=FALSE)


P4=shared_1000_abd1_RH_AT[,c(4:ncol(shared_1000_abd1_RH_AT))]

row.names(P4)=shared_1000_abd1_RH_AT$Group

A.hel=decostand(P4, "hellinger", na.rm=TRUE)

write.table(A.hel, file="hellinger_rpoB_AT_RH.tab", col.names=TRUE, row.names = TRUE, quote=FALSE)

A.hel.D=dist(A.hel)

A.hel.D_AT_RH=as.matrix(A.hel.D)

write.table(A.hel.D_AT_RH, file="hellinger_distance_matrix_rpoB_AT_RH.txt", col.names=TRUE, row.names = TRUE, quote=FALSE)
```

**## perform PCoA**

```
library("ape")
```

**## extract the eigenvalues and PCoA axis**

```
res_AT_R=pcoa(A.hel.D_AT_R)

res_AT_RH=pcoa(A.hel.D_AT_RH)

res_SP_R=pcoa(A.hel.D_SP_R)

res_SP_RH=pcoa(A.hel.D_SP_RH)


write.table(res_AT_R$values, file="eighvalues_pcoa_rpoB_hel_AT_R.txt", col.names=TRUE, row.names = TRUE, quote=FALSE)
```

```
write.table(res_AT_RH$values, file="eighvalues_pcoa_rpoB_hel_AT_RH.txt", col.names=TRUE, row.names = TRUE, quote=FALSE)

write.table(res_SP_R$values, file="eighvalues_pcoa_rpoB_hel_SP_R.txt", col.names=TRUE, row.names = TRUE, quote=FALSE)

write.table(res_SP_RH$values, file="eighvalues_pcoa_rpoB_hel_SP_RH.txt", col.names=TRUE, row.names = TRUE, quote=FALSE)


axes_AT_R=res_AT_R$vectors

axes_AT_RH=res_AT_RH$vectors

axes_SP_R=res_SP_R$vectors

axes_SP_RH=res_SP_RH$vectors


tiff("PCOA_AT_R.tiff", res=600, width=15, height=15, unit="cm",compression="lzw")

plot(axes_AT_R[,1:2])

dev.off()


tiff("PCOA_AT_RH.tiff", res=600, width=15, height=15, unit="cm",compression="lzw")

plot(axes_AT_RH[,1:2])

dev.off()


tiff("PCOA_SP_R.tiff", res=600, width=15, height=15, unit="cm",compression="lzw")

plot(axes_SP_R[,1:2])

dev.off()


tiff("PCOA_SP_RH.tiff", res=600, width=15, height=15, unit="cm",compression="lzw")

plot(axes_SP_RH[,1:2])

dev.off()


axes_AT_R_4=axes_AT_R[,1:4]

axes_AT_RH_4=axes_AT_RH[,1:4]

axes_SP_R_4=axes_SP_R[,1:4]

axes_SP_RH_4=axes_SP_RH[,1:4]
```

write.table(axes_AT_R_4, file="pcoa_axis_rpoB_hel_AT_R.txt", col.names=TRUE, row.names = TRUE, quote=FALSE)

write.table(axes_AT_RH_4, file="pcoa_axis_rpoB_hel_AT_RH.txt", col.names=TRUE, row.names = TRUE, quote=FALSE)

write.table(axes_SP_R_4, file="pcoa_axis_rpoB_hel_SP_R.txt", col.names=TRUE, row.names = TRUE, quote=FALSE)

write.table(axes_SP_RH_4, file="pcoa_axis_rpoB_hel_SP_RH.txt", col.names=TRUE, row.names = TRUE, quote=FALSE)

**##Separate data in 4 groups (AT/SP, R/RH)**

**##Insert Rep column**

groups.summary <- read.table("shared_1000_abd1groups.summary", header=T)

groups.summary$group[314] <- str_sub(groups.summary$group[314], end = -2)

stri_sub(groups.summary$group, -1, 1) <- "_"

groups.summary <- groups.summary[-1]

groups.summary$rep <- str_split_fixed(groups.summary$group, '_', 5)[,5]

groups.summary <- groups.summary[,c(1,ncol(groups.summary),3:ncol(groups.summary)-1)]

groups.summary$combo <- paste(str_split_fixed(groups.summary$group, '_', 5)[,4], str_split_fixed(groups.summary$group, '_', 5)[,3], sep= "_")

groups.summary_RH_AT <- as.data.frame(split(groups.summary, f=groups.summary$combo)["RH_AT"])

groups.summary_RH_SP <- as.data.frame(split(groups.summary, f=groups.summary$combo)["RH_SP"])

groups.summary_R_AT <- as.data.frame(split(groups.summary, f=groups.summary$combo)["R_AT"])

groups.summary_R_SP <- as.data.frame(split(groups.summary, f=groups.summary$combo)["R_SP"])

groups.summary_RH_AT <- groups.summary_RH_AT[-ncol(groups.summary_RH_AT)]

groups.summary_RH_SP <- groups.summary_RH_SP[-ncol(groups.summary_RH_SP)]

groups.summary_R_AT <- groups.summary_R_AT[-ncol(groups.summary_R_AT)]

groups.summary_R_SP <- groups.summary_R_SP[-ncol(groups.summary_R_SP)]

colnames(groups.summary_RH_AT) <- str_sub(colnames(groups.summary_RH_AT), start = 7)

colnames(groups.summary_RH_SP) <- str_sub(colnames(groups.summary_RH_SP), start = 7) colnames(groups.summary_R_AT) <- str_sub(colnames(groups.summary_R_AT), start = 6)

colnames(groups.summary_R_SP) <- str_sub(colnames(groups.summary_R_SP), start = 6)

**## Insert PCOA axes**

axes_AT_RH_4 <- read.table("pcoa_axis_rpoB_hel_AT_RH.txt", header=T)

axes_AT_R_4 <- read.table("pcoa_axis_rpoB_hel_AT_R.txt", header=T)

axes_SP_RH_4 <- read.table("pcoa_axis_rpoB_hel_SP_RH.txt", header=T)

axes_SP_R_4 <- read.table("pcoa_axis_rpoB_hel_SP_R.txt", header=T)

axes_AT_RH_4$group <- rownames(axes_AT_RH_4)

axes_AT_R_4$group <- rownames(axes_AT_R_4)

axes_SP_RH_4$group <- rownames(axes_SP_RH_4)

axes_SP_R_4$group <- rownames(axes_SP_R_4)

groups.summary_RH_AT_pcoa <- merge(groups.summary_RH_AT,axes_AT_RH_4, by.x="group")

groups.summary_RH_SP_pcoa <- merge(groups.summary_RH_SP,axes_SP_RH_4, by="group")

groups.summary_R_AT_pcoa <- merge(groups.summary_R_AT,axes_AT_R_4, by="group")

groups.summary_R_SP_pcoa <- merge(groups.summary_R_SP,axes_SP_R_4, by="group")

**#Delete samples at shannon -0.000**

groups.summary_RH_AT_pcoa <- groups.summary_RH_AT_pcoa[groups.summary_RH_AT_pcoa$npshannon != 0,]

groups.summary_RH_SP_pcoa <- groups.summary_RH_SP_pcoa[groups.summary_RH_SP_pcoa$npshannon != 0,]

groups.summary_R_AT_pcoa <- groups.summary_R_AT_pcoa[groups.summary_R_AT_pcoa$npshannon != 0,]

groups.summary_R_SP_pcoa <- groups.summary_R_SP_pcoa[groups.summary_R_SP_pcoa$npshannon != 0,]

**#Transformation in presence/absence matrix**

**#Selection of the most prevalent OTUs for each combination**

shared_1000_abd1_R_AT_1=shared_1000_abd1_R_AT[,c(4:ncol(shared_1000_abd1_R_AT))]

shared_1000_abd1_R_SP_1=shared_1000_abd1_R_SP[,c(4:ncol(shared_1000_abd1_R_SP))]

```
shared_1000_abd1_RH_AT_1=shared_1000_abd1_RH_AT[,c(4:ncol(shared_1000_abd1_RH_AT))]
shared_1000_abd1_RH_SP_1=shared_1000_abd1_RH_SP[,c(4:ncol(shared_1000_abd1_RH_SP))]

row.names(shared_1000_abd1_R_AT_1)=shared_1000_abd1_R_AT$Group
row.names(shared_1000_abd1_R_SP_1)=shared_1000_abd1_R_SP$Group
row.names(shared_1000_abd1_RH_AT_1)=shared_1000_abd1_RH_AT$Group
row.names(shared_1000_abd1_RH_SP_1)=shared_1000_abd1_RH_SP$Group

shared_1000_abd1_R_AT_PA <- t((shared_1000_abd1_R_AT_1>0)*1L)
shared_1000_abd1_R_SP_PA <- t((shared_1000_abd1_R_SP_1>0)*1L)
shared_1000_abd1_RH_AT_PA <- t((shared_1000_abd1_RH_AT_1>0)*1L)
shared_1000_abd1_RH_SP_PA <- t((shared_1000_abd1_RH_SP_1>0)*1L)

shared_1000_abd1_R_AT_PA <- as.data.frame(shared_1000_abd1_R_AT_PA)
shared_1000_abd1_R_SP_PA <- as.data.frame(shared_1000_abd1_R_SP_PA)
shared_1000_abd1_RH_AT_PA <- as.data.frame(shared_1000_abd1_RH_AT_PA)
shared_1000_abd1_RH_SP_PA <- as.data.frame(shared_1000_abd1_RH_SP_PA)

shared_1000_abd1_R_AT_PA$Sums <- rowSums(shared_1000_abd1_R_AT_PA)
shared_1000_abd1_R_SP_PA$Sums <- rowSums(shared_1000_abd1_R_SP_PA)
shared_1000_abd1_RH_AT_PA$Sums <- rowSums(shared_1000_abd1_RH_AT_PA)
shared_1000_abd1_RH_SP_PA$Sums <- rowSums(shared_1000_abd1_RH_SP_PA)
```

**# OTUs with at least sum of 10 samples**

```
shared_1000_abd1_R_AT_PA <- shared_1000_abd1_R_AT_PA[shared_1000_abd1_R_AT_PA$Sums >= 10,]
shared_1000_abd1_R_SP_PA <- shared_1000_abd1_R_SP_PA[shared_1000_abd1_R_SP_PA$Sums >= 10,]
shared_1000_abd1_RH_AT_PA <- shared_1000_abd1_RH_AT_PA[shared_1000_abd1_RH_AT_PA$Sums >= 10,]
shared_1000_abd1_RH_SP_PA <- shared_1000_abd1_RH_SP_PA[shared_1000_abd1_RH_SP_PA$Sums >= 10,]
```

shared_1000_abd1_R_AT_PA <- head(t(shared_1000_abd1_R_AT_PA), -1)

shared_1000_abd1_R_SP_PA <- head(t(shared_1000_abd1_R_SP_PA), -1)

shared_1000_abd1_RH_AT_PA <- head(t(shared_1000_abd1_RH_AT_PA), -1)

shared_1000_abd1_RH_SP_PA <- head(t(shared_1000_abd1_RH_SP_PA), -1)

shared_1000_abd1_R_AT_PA <- cbind(rownames(shared_1000_abd1_R_AT_PA), data.frame(shared_1000_abd1_R_AT_PA, row.names=NULL))

shared_1000_abd1_R_SP_PA <- cbind(rownames(shared_1000_abd1_R_SP_PA), data.frame(shared_1000_abd1_R_SP_PA, row.names=NULL))

shared_1000_abd1_RH_AT_PA <- cbind(rownames(shared_1000_abd1_RH_AT_PA), data.frame(shared_1000_abd1_RH_AT_PA, row.names=NULL))

shared_1000_abd1_RH_SP_PA <- cbind(rownames(shared_1000_abd1_RH_SP_PA), data.frame(shared_1000_abd1_RH_SP_PA, row.names=NULL))

names(shared_1000_abd1_R_AT_PA)[names(shared_1000_abd1_R_AT_PA) == "rownames(shared_1000_abd1_R_AT_PA)"] <- "sample"

names(shared_1000_abd1_R_SP_PA)[names(shared_1000_abd1_R_SP_PA) == "rownames(shared_1000_abd1_R_SP_PA)"] <- "sample"

names(shared_1000_abd1_RH_AT_PA)[names(shared_1000_abd1_RH_AT_PA) == "rownames(shared_1000_abd1_RH_AT_PA)"] <- "sample"

names(shared_1000_abd1_RH_SP_PA)[names(shared_1000_abd1_RH_SP_PA) == "rownames(shared_1000_abd1_RH_SP_PA)"] <- "sample"

**# Creation of stat files**

groups.summary_RH_AT_pcoa$Pop <- paste(str_split_fixed(groups.summary_RH_AT_pcoa$group, '_', 5)[,1], str_split_fixed(groups.summary_RH_AT_pcoa$group, '_', 5)[,2], sep= "_")

groups.summary_RH_AT_pcoa$Season <- str_split_fixed(groups.summary_RH_AT_pcoa$group, '_', 5)[,3]

groups.summary_RH_AT_pcoa$Compartment <- str_split_fixed(groups.summary_RH_AT_pcoa$group, '_', 5)[,4]

groups.summary_RH_SP_pcoa$Pop <- paste(str_split_fixed(groups.summary_RH_SP_pcoa$group, '_', 5)[,1], str_split_fixed(groups.summary_RH_SP_pcoa$group, '_', 5)[,2], sep= "_")

groups.summary_RH_SP_pcoa$Season <- str_split_fixed(groups.summary_RH_SP_pcoa$group, '_', 5)[,3]

groups.summary_RH_SP_pcoa$Compartment <- str_split_fixed(groups.summary_RH_SP_pcoa$group, '_', 5)[,4] groups.summary_R_AT_pcoa$Pop <- paste(str_split_fixed(groups.summary_R_AT_pcoa$group, '_', 5)[,1], str_split_fixed(groups.summary_R_AT_pcoa$group, '_', 5)[,2], sep= "_")

groups.summary_R_AT_pcoa$Season <- str_split_fixed(groups.summary_R_AT_pcoa$group, '_', 5)[,3]

groups.summary_R_AT_pcoa$Compartment <- str_split_fixed(groups.summary_R_AT_pcoa$group, '_', 5)[,4]

groups.summary_R_SP_pcoa$Pop <- paste(str_split_fixed(groups.summary_R_SP_pcoa$group, '_', 5)[,1], str_split_fixed(groups.summary_R_SP_pcoa$group, '_', 5)[,2], sep= "_")

groups.summary_R_SP_pcoa$Season <- str_split_fixed(groups.summary_R_SP_pcoa$group, '_', 5)[,3]

groups.summary_R_SP_pcoa$Compartment <- str_split_fixed(groups.summary_R_SP_pcoa$group, '_', 5)[,4]

**#Estimation of BLUPS for GEA analysis. BLUPs were setimated in the separated datasets**

blups_AT_RH <- subset(groups.summary_RH_AT_pcoa, select = -c(invsimpson_lci, invsimpson_hci, chao, chao_lci, chao_hci, Season, Compartment))

blups_SP_RH <- subset(groups.summary_RH_SP_pcoa, select = -c(invsimpson_lci, invsimpson_hci, chao, chao_lci, chao_hci, Season, Compartment))

blups_AT_R <- subset(groups.summary_R_AT_pcoa, select = -c(invsimpson_lci, invsimpson_hci, chao, chao_lci, chao_hci, Season, Compartment))

blups_SP_R <- subset(groups.summary_R_SP_pcoa, select = -c(invsimpson_lci, invsimpson_hci, chao, chao_lci, chao_hci, Season, Compartment))

blups_AT_RH <- blups_AT_RH[, c(1, 10, 2, 3, 4, 5, 6, 7, 8, 9)]

blups_SP_RH <- blups_SP_RH[, c(1, 10, 2, 3, 4, 5, 6, 7, 8, 9)]

blups_AT_R <- blups_AT_R[, c(1, 10, 2, 3, 4, 5, 6, 7, 8, 9)]

blups_SP_R <- blups_SP_R[, c(1, 10, 2, 3, 4, 5, 6, 7, 8, 9)]

blups_AT_R <- na.omit(qpcR:::cbind.na(blups_AT_R, shared_1000_abd1_R_AT_PA))

blups_SP_R <- na.omit(qpcR:::cbind.na(blups_SP_R, shared_1000_abd1_R_SP_PA))

blups_AT_RH <- na.omit(qpcR:::cbind.na(blups_AT_RH, shared_1000_abd1_RH_AT_PA))

blups_SP_RH <- na.omit(qpcR:::cbind.na(blups_SP_RH, shared_1000_abd1_RH_SP_PA))

blups_AT_R <- subset(blups_AT_R, select = -c(sample))

```
blups_SP_R <- subset(blups_SP_R, select = -c(sample))

blups_AT_RH <- subset(blups_AT_RH, select = -c(sample))

blups_SP_RH <- subset(blups_SP_RH, select = -c(sample))


dim(blups_AT_R)

#102 29

dim(blups_AT_RH)

#106 20

dim(blups_SP_R)

#116 30

dim(blups_SP_RH)

#101 15


BLUPS_R_AT=as.data.frame(matrix(NA, ncol=length(blups_AT_R)+1, nrow=length(unique(blups_AT_R$Pop))))

names(BLUPS_R_AT)[1]="Pop"

BLUPS_R_AT$Pop=unique(blups_AT_R$Pop)

names(BLUPS_R_AT)[2:length(colnames(BLUPS_R_AT))]=colnames(blups_AT_R)

BLUPS_R_AT <- BLUPS_R_AT[,-c(2:4)]


for (trait in colnames(blups_AT_R[,c(4:ncol(blups_AT_R))])){

# fit the model

 mixed.lmer<- lmer(blups_AT_R[,which(colnames(blups_AT_R)==trait)] ~ (1|rep)+(1|Pop), data = blups_AT_R)


# estimate BLUPS

  blup = ranef(mixed.lmer)


# extract blup for Population

  BLUPpop = blup$Pop


  BLUPpop=cbind(Pop= rownames(BLUPpop), BLUPpop)

  rownames(BLUPpop)=NULL
```

```
  BLUPpop$mean=NA

  for(i in unique(BLUPpop$Pop)){
    mean=mean(na.omit(blups_AT_R[blups_AT_R$Pop==i, which(colnames(blups_AT_R)==trait)]))
    BLUPpop[match(i, BLUPpop$Pop), which(colnames(BLUPpop)=="mean")]=mean
  }

  BLUPpop$BLUP=BLUPpop$`(Intercept)`+ BLUPpop$mean
  colnames(BLUPpop)[4] <- trait

  write.table(BLUPpop, paste("./BLUP_", trait, ".txt", sep=""), sep="\t", quote=F, row.names=F)
  BLUPS_R_AT[, which(colnames(BLUPS_R_AT)==trait)]=BLUPpop[4]
}
write.table(BLUPS_R_AT, file="BLUPS_R_AT.txt", col.names=TRUE, row.names = TRUE, quote=FALSE)
BLUP_AT_R_deheaded_t <- as.data.frame(t(BLUPS_R_AT)[-1,])
BLUP_AT_R_deheaded_t <- apply(BLUP_AT_R_deheaded_t, 2, str_remove_all, " ")
write.table(BLUP_AT_R_deheaded_t, file="BLUP_AT_R_deheaded_t.txt", col.names=FALSE, row.names = FALSE, quote=FALSE)
i <- 0
poolsize <- list()
while(i < nrow(BLUP_AT_R_deheaded_t)){
   poolsize <- append(poolsize, "30")
   i = i+1
}
write.table(poolsize, file="poolsize_R_AT.txt", col.names= FALSE, row.names = FALSE, quote=FALSE)

BLUPS_RH_AT=as.data.frame(matrix(NA, ncol=length(blups_AT_RH)+1, nrow=length(unique(blups_AT_RH$Pop))))
names(BLUPS_RH_AT)[1]="Pop"
BLUPS_RH_AT$Pop=unique(blups_AT_RH$Pop)
names(BLUPS_RH_AT)[2:length(colnames(BLUPS_RH_AT))]=colnames(blups_AT_RH)
```

```
BLUPS_RH_AT <- BLUPS_RH_AT[,-c(2:4)]


for (trait in colnames(blups_AT_RH[,c(4:ncol(blups_AT_RH))])){
  # fit the model
  mixed.lmer<- lmer(blups_AT_RH[,which(colnames(blups_AT_RH)==trait)] ~ (1|rep)+(1|Pop), data
= blups_AT_RH)


  # estimate BLUPS
  blup = ranef(mixed.lmer)


  # extract blup for Population
  BLUPpop = blup$Pop


  BLUPpop=cbind(Pop= rownames(BLUPpop), BLUPpop)
  rownames(BLUPpop)=NULL


  BLUPpop$mean=NA


  for(i in unique(BLUPpop$Pop)){
    mean=mean(na.omit(blups_AT_RH[blups_AT_RH$Pop==i,
which(colnames(blups_AT_RH)==trait)]))
    BLUPpop[match(i, BLUPpop$Pop), which(colnames(BLUPpop)=="mean")]=mean
  }


  BLUPpop$BLUP=BLUPpop$`(Intercept)`+ BLUPpop$mean
  colnames(BLUPpop)[4] <- trait


  write.table(BLUPpop, paste("./BLUP_", trait, ".txt", sep=""), sep="\t", quote=F, row.names=F)
  BLUPS_RH_AT[, which(colnames(BLUPS_RH_AT)==trait)]=BLUPpop[4]
}
write.table(BLUPS_RH_AT, file="BLUPS_RH_AT.txt", col.names=TRUE, row.names = TRUE,
quote=FALSE)
BLUP_AT_RH_deheaded_t <- as.data.frame(t(BLUPS_RH_AT)[-1,])
```

```
BLUP_AT_RH_deheaded_t <- apply(BLUP_AT_RH_deheaded_t, 2, str_remove_all, " ")

write.table(BLUP_AT_RH_deheaded_t, file="BLUP_AT_RH_deheaded_t.txt", col.names=FALSE, row.names = FALSE, quote=FALSE)

i <- 0

poolsize <- list()

while(i < nrow(BLUP_AT_RH_deheaded_t)){

  poolsize <- append(poolsize, "30")

  i = i+1

}

write.table(poolsize, file="poolsize_RH_AT.txt", col.names= FALSE, row.names = FALSE, quote=FALSE)


BLUPS_RH_SP=as.data.frame(matrix(NA, ncol=length(blups_SP_RH)+1, nrow=length(unique(blups_SP_RH$Pop))))

names(BLUPS_RH_SP)[1]="Pop"

BLUPS_RH_SP$Pop=unique(blups_SP_RH$Pop)

names(BLUPS_RH_SP)[2:length(colnames(BLUPS_RH_SP))]=colnames(blups_SP_RH)

BLUPS_RH_SP <- BLUPS_RH_SP[,-c(2:4)]


for (trait in colnames(blups_SP_RH[,c(4:ncol(blups_SP_RH))])){
 # fit the model
 mixed.lmer<- lmer(blups_SP_RH[,which(colnames(blups_SP_RH)==trait)] ~ (1|rep)+(1|Pop), data = blups_SP_RH)


 # estimate BLUPS
 blup = ranef(mixed.lmer)


 # extract blup for Population
 BLUPpop = blup$Pop


 BLUPpop=cbind(Pop= rownames(BLUPpop), BLUPpop)
 rownames(BLUPpop)=NULL
```

```
  BLUPpop$mean=NA

  for(i in unique(BLUPpop$Pop)){
    mean=mean(na.omit(blups_SP_RH[blups_SP_RH$Pop==i,
which(colnames(blups_SP_RH)==trait)]))
    BLUPpop[match(i, BLUPpop$Pop), which(colnames(BLUPpop)=="mean")]=mean
  }

  BLUPpop$BLUP=BLUPpop$`(Intercept)`+ BLUPpop$mean
  colnames(BLUPpop)[4] <- trait

  write.table(BLUPpop, paste("./BLUP_", trait, ".txt", sep=""), sep="\t", quote=F, row.names=F)
  BLUPS_RH_SP[, which(colnames(BLUPS_RH_SP)==trait)]=BLUPpop[4]
}
write.table(BLUPS_RH_SP, file="BLUPS_RH_SP.txt", col.names=TRUE, row.names = TRUE,
quote=FALSE)
BLUP_SP_RH_deheaded_t <- as.data.frame(t(BLUPS_RH_SP)[-1,])
BLUP_SP_RH_deheaded_t <- apply(BLUP_SP_RH_deheaded_t, 2, str_remove_all, " ")
write.table(BLUP_SP_RH_deheaded_t, file="BLUP_SP_RH_deheaded_t.txt", col.names=FALSE,
row.names = FALSE, quote=FALSE)
i <- 0
poolsize <- list()
while(i < nrow(BLUP_SP_RH_deheaded_t)){
   poolsize <- append(poolsize, "30")
   i = i+1
}
write.table(poolsize, file="poolsize_RH_SP.txt", col.names= FALSE, row.names = FALSE,
quote=FALSE)

BLUPS_R_SP=as.data.frame(matrix(NA, ncol=length(blups_SP_R)+1,
nrow=length(unique(blups_SP_R$Pop))))
names(BLUPS_R_SP)[1]="Pop"
BLUPS_R_SP$Pop=unique(blups_SP_R$Pop)
names(BLUPS_R_SP)[2:length(colnames(BLUPS_R_SP))]=colnames(blups_SP_R)
```

```
BLUPS_R_SP <- BLUPS_R_SP[,-c(2:4)]


for (trait in colnames(blups_SP_R[,c(4:ncol(blups_SP_R))])){
  # fit the model
  mixed.lmer<- lmer(blups_SP_R[,which(colnames(blups_SP_R)==trait)] ~ (1|rep)+(1|Pop), data = blups_SP_R)


  # estimate BLUPS
  blup = ranef(mixed.lmer)


  # extract blup for Population
  BLUPpop = blup$Pop


  BLUPpop=cbind(Pop= rownames(BLUPpop), BLUPpop)
  rownames(BLUPpop)=NULL


  BLUPpop$mean=NA


  for(i in unique(BLUPpop$Pop)){
    mean=mean(na.omit(blups_SP_R[blups_SP_R$Pop==i, which(colnames(blups_SP_R)==trait)]))
    BLUPpop[match(i, BLUPpop$Pop), which(colnames(BLUPpop)=="mean")]=mean
  }


  BLUPpop$BLUP=BLUPpop$`(Intercept)`+ BLUPpop$mean
  colnames(BLUPpop)[4] <- trait


  write.table(BLUPpop, paste("./BLUP_", trait, ".txt", sep=""), sep="\t", quote=F, row.names=F)
  BLUPS_R_SP[, which(colnames(BLUPS_R_SP)==trait)]=BLUPpop[4]
}
write.table(BLUPS_R_SP, file="BLUPS_R_SP.txt", col.names=TRUE, row.names = TRUE, quote=FALSE)
BLUP_SP_R_deheaded_t <- as.data.frame(t(BLUPS_R_SP)[-1,])
BLUP_SP_R_deheaded_t <- apply(BLUP_SP_R_deheaded_t, 2, str_remove_all, " ")
```

write.table(BLUP_SP_R_deheaded_t, file="BLUP_SP_R_deheaded_t.txt", col.names=FALSE, row.names = FALSE, quote=FALSE)

i <- 0

poolsize <- list()

while(i < nrow(BLUP_SP_R_deheaded_t)){

poolsize <- append(poolsize, "30")

i = i+1

}

write.table(poolsize, file="poolsize_R_SP.txt", col.names= FALSE, row.names = FALSE, quote=FALSE)

**# Figures on alpha diversity indices**

**# Creation of rpoB_stats_merged.txt**

rpoB_stats_merged <- rbind(groups.summary_R_AT_pcoa, groups.summary_R_SP_pcoa, groups.summary_RH_AT_pcoa, groups.summary_RH_SP_pcoa)

rpoB_stats_merged <- subset(rpoB_stats_merged, select = -c(invsimpson_lci, invsimpson_hci, chao, chao_lci, chao_hci, Axis.3, Axis.4))

rpoB_stats_merged <- rpoB_stats_merged[, c(1, 8, 2, 9, 10, 3, 4, 5, 6, 7)]

**# boxplots**

tiff("rpoB_Shannon_compartment.tiff", res=600, width=15, height=15, unit="cm",compression="lzw")

boxplot(npshannon~Compartment, data=rpoB_stats_merged, ylab = "Shannon's Index", xlab="",main = "",

cex.main=1, cex.axis=0.8, las=2, at = c(1,2), axes=F)

box()

axis(2, at=seq(0,3,0.5), seq(0,3,0.5), las=1)

axis(1, at=c(1,2),labels = c("Roots","Rhizosphere"), cex.axis=0.8,las=1,side=1)

dev.off()

tiff("rpoB_Richness_compartment.tiff", res=600, width=15, height=15, unit="cm",compression="lzw")

boxplot(sobs~Compartment, data=rpoB_stats_merged, ylab = "Richness", xlab="",main = "",

cex.main=1, cex.axis=0.8, las=2, at = c(1,2), axes=F)

```
box()
axis(2, at=seq(0,30,5), seq(0,30,5), las=1)
axis(1, at=c(1,2),labels = c("Roots","Rhizosphere"), cex.axis=0.8,las=1,side=1)
dev.off()

tiff("rpoB_Shannon.tiff", res=600, width=15, height=15, unit="cm",compression="lzw")
boxplot(npshannon~Season*Compartment, data=rpoB_stats_merged, ylab = "Shannon's Index", xlab="",main = "",
cex.main=1, cex.axis=0.8, las=2, at = c(1,2, 4,5), axes=F)
box()
axis(2, at=seq(0,3,0.5), seq(0,3,0.5), las=1)
axis(1, at=c(1,2, 4,5),labels = c("Autumn","Spring", "Autumn","Spring"), cex.axis=0.8,las=1,side=1)
mtext("Roots", side = 1, line = 3, adj = 0.188, cex=0.8)
mtext("Rhizosphere", side = 1, line = 3, adj = 0.835, cex=0.8)
axis(1, at=c(1,2), line=3, tick=T, labels=rep("",2), lwd=1, lwd.ticks=0)
axis(1, at=3+c(1,2),line=3,tick=T,labels=rep("",2), lwd=1, lwd.ticks=0)
dev.off()

df_filtered <- subset(rpoB_stats_merged, Compartment == "RH")
tiff("rpoB_Shannon_RH.tiff", res=600, width=15, height=15, unit="cm",compression="lzw")
boxplot(npshannon~Season, data=df_filtered, ylab = "Shannon's Index", xlab="",main = "",
cex.main=1, cex.axis=0.8, las=2, at = c(1,2), axes=F)
box()
axis(2, at=seq(0,3,0.5), seq(0,3,0.5), las=1)
axis(1, at=c(1,2),labels = c("Autumn","Spring"), cex.axis=0.8,las=1,side=1)
mtext("Rhizosphere", side = 1, line = 3, adj = 0.188, cex=0.8)
axis(1, at=c(1,2), line=3, tick=T, labels=rep("",2), lwd=1, lwd.ticks=0)
dev.off()

df_filtered <- subset(rpoB_stats_merged, Compartment == "RH")
tiff("rpoB_Richness_RH.tiff", res=600, width=15, height=15, unit="cm",compression="lzw")
boxplot(sobs~Season, data=df_filtered, ylab = "Richness", xlab="",main = "",
```

```
cex.main=1, cex.axis=0.8, las=2, at = c(1,2), axes=F)
box()
axis(2, at=seq(0,30,5), seq(0,30,5), las=1)
axis(1, at=c(1,2),labels = c("Autumn","Spring"), cex.axis=0.8,las=1,side=1)
dev.off()
```

**#Test of compartment effet on microbiota variables**

```
rpoB_stats_merged$rep <- as.numeric(rpoB_stats_merged$rep)

F_stat <- glm(rpoB_stats_merged$sobs~Compartment +(1|rep), data = rpoB_stats_merged)
summary(F_stat)
Anova(F_stat)

F_stat <- glm(rpoB_stats_merged$npshannon~Compartment +(1|rep), data = rpoB_stats_merged)
summary(F_stat)
Anova(F_stat)

F_stat <- glm(rpoB_stats_merged$Axis.1~Compartment +(1|rep), data = rpoB_stats_merged)
summary(F_stat)
Anova(F_stat)

F_stat <- glm(rpoB_stats_merged$Axis.2~Compartment +(1|rep), data = rpoB_stats_merged)
summary(F_stat)
Anova(F_stat)
```

**#Test of season effect on microbiota variables for each compartment**

```
data_R=rpoB_stats_merged[which(rpoB_stats_merged$Compartment=='R'),]

F_stat <- glm(data_R$sobs~Season +(1|rep), data = data_R)
summary(F_stat)
Anova(F_stat)
```

```
F_stat <- glm(data_R$npshannon~Season +(1|rep), data = data_R)
summary(F_stat)
Anova(F_stat)

F_stat <- glm(data_R$Axis.1~Season +(1|rep), data = data_R)
summary(F_stat)
Anova(F_stat)

F_stat <- glm(data_R$Axis.2~Season +(1|rep), data = data_R)
summary(F_stat)
Anova(F_stat)

data_RH=rpoB_stats_merged[which(rpoB_stats_merged$Compartment=='RH'),]

F_stat <- glm(data_RH$sobs~Season +(1|rep), data = data_RH)
summary(F_stat)
Anova(F_stat)

F_stat <- glm(data_RH$npshannon~Season +(1|rep), data = data_RH)
summary(F_stat)
Anova(F_stat)

F_stat <- glm(data_RH$Axis.1~Season +(1|rep), data = data_RH)
summary(F_stat)
Anova(F_stat)

F_stat <- glm(data_RH$Axis.2~Season +(1|rep), data = data_RH)
summary(F_stat)
Anova(F_stat)

# boxplots genotypes
data_R_AT=data_R[which(data_R$Season=='AT'),]
```

```
data_R_SP=data_R[which(data_R$Season=='SP'),]

data_RH_AT=data_RH[which(data_RH$Season=='AT'),]

data_RH_SP=data_RH[which(data_RH$Season=='SP'),]

data_RH_SP = data_RH_SP[!(row.names(data_RH_SP) %in% c("NA","NA.1")),]



tiff("rpoB_Shannon_R_season_genotype.tiff", res=600, width=15, height=15, unit="cm",compression="lzw")

boxplot(npshannon~Pop*Season, data=data_R, ylab = "Shannon's Index", xlab="",main = "",

cex.main=1, cex.axis=0.8, las=2, at = c(1,2,3,4,5,6,7,8,9,10,11,12,13,14,15,16,17,18,19,20,21,22,23,24,25
,28,29,30,31,32,33,34,35,36,37,38,39,40,41,42,43,44,45,46,47,48,49,50,51,52), axes=F)

box()

axis(2, at=seq(0,3,0.5), seq(0,3,0.5), las=1)

axis(1, at=c(1,2,3,4,5,6,7,8,9,10,11,12,13,14,15,16,17,18,19,20,21,22,23,24,25
,28,29,30,31,32,33,34,35,36,37,38,39,40,41,42,43,44,45,46,47,48,49,50,51,52),

labels = c("ALTO-A","APIG-A","BISA-A","BOEL-A","BOLO-A","BRUZ-A","CAST-A","CAST-B","CAST-C","COLL-A","CORL-A","FOUG-A","GRAT-A","GRAT-B","ISNE-A","LASC-A","MARI-A","MARI-B", "MISI-A","MISI-B","MONR-A","MORD-A","PIAN-A","POLL-A","RENN-A","ALTO-A","APIG-A","BISA-A","BOEL-A","BOLO-A","BRUZ-A","CAST-A", "CAST-B","CAST-C","COLL-A","CORL-A","FOUG-A","GRAT-A","GRAT-B","ISNE-A","LASC-A","MARI-A","MARI-B", "MISI-A","MISI-B","MONR-A","MORD-A","PIAN-A","POLL-A","RENN-A" ), cex.axis=0.4,las=2)

mtext("Autumn", side = 1, line = 4, adj = 0.170, cex=1, mgp=c(3,3,0))

mtext("Spring", side = 1, line = 4, adj = 0.800, cex=1, mgp=c(3,3,0))

dev.off()



tiff("rpoB_Shannon_RH_season_genotype.tiff", res=600, width=15, height=15, unit="cm",compression="lzw")

boxplot(npshannon~Pop*Season, data=data_RH, ylab = "Shannon's Index", xlab="",main = "",

cex.main=1, cex.axis=0.8, las=2, at = c(1,2,3,4,5,6,7,8,9,10,11,12,13,14,15,16,17,18,19,20,21,22,23,24,25,26
,29,30,31,32,33,34,35,36,37,38,39,40,41,42,43,44,45,46,47,48,49,50,51,52,53,54), axes=F)

box()

axis(2, at=seq(0,3,0.5), seq(0,3,0.5), las=1)

axis(1, at=c(1,2,3,4,5,6,7,8,9,10,11,12,13,14,15,16,17,18,19,20,21,22,23,24,25,26
,29,30,31,32,33,34,35,36,37,38,39,40,41,42,43,44,45,46,47,48,49,50,51,52,53,54),
```

labels = c("ALTO-A","APIG-A","BISA-A","BOEL-A","BOLO-A","BRUZ-A","CAST-A", "CAST-B","CAST-C","COLL-A","CORL-A","FOUG-A","GRAT-A","GRAT-B","ISNE-A","ISNE-B","LASC-A","MARI-A","MARI-B", "MISI-A","MISI-B","MONR-A","MORD-A","PIAN-A","POLL-A","RENN-A","ALTO-A","APIG-A","BISA-A","BOEL-A","BOLO-A","BRUZ-A","CAST-A", "CAST-B","CAST-C","COLL-A","CORL-A","FOUG-A","GRAT-A","GRAT-B","ISNE-A","ISNE-B","LASC-A","MARI-A","MARI-B", "MISI-A","MISI-B","MONR-A","MORD-A","PIAN-A","POLL-A","RENN-A" ), cex.axis=0.4,las=2)

mtext("Autumn", side = 1, line = 4, adj = 0.170, cex=1, mgp=c(3,3,0))

mtext("Spring", side = 1, line = 4, adj = 0.800, cex=1, mgp=c(3,3,0))

dev.off()

tiff("rpoB_Richness_R_season_genotype.tiff", res=600, width=15, height=15, unit="cm",compression="lzw")

boxplot(npshannon~Pop*Season, data=data_R, ylab = "Richness", xlab="",main = "",

cex.main=1, cex.axis=0.8, las=2, at = c(1,2,3,4,5,6,7,8,9,10,11,12,13,14,15,16,17,18,19,20,21,22,23,24,25 ,28,29,30,31,32,33,34,35,36,37,38,39,40,41,42,43,44,45,46,47,48,49,50,51,52), axes=F)

box()

axis(2, at=seq(0,30,5), seq(0,30,5), las=1)

axis(1, at=c(1,2,3,4,5,6,7,8,9,10,11,12,13,14,15,16,17,18,19,20,21,22,23,24,25, 28,29,30,31,32,33,34,35,36,37,38,39,40,41,42,43,44,45,46,47,48,49,50,51,52),

labels = c("ALTO-A","APIG-A","BISA-A","BOEL-A","BOLO-A","BRUZ-A","CAST-A", "CAST-B","CAST-C","COLL-A","CORL-A","FOUG-A","GRAT-A","GRAT-B","ISNE-A","LASC-A","MARI-A","MARI-B", "MISI-A","MISI-B","MONR-A","MORD-A","PIAN-A","POLL-A","RENN-A","ALTO-A","APIG-A","BISA-A","BOEL-A","BOLO-A","BRUZ-A","CAST-A", "CAST-B","CAST-C","COLL-A","CORL-A","FOUG-A","GRAT-A","GRAT-B","ISNE-A","LASC-A","MARI-A","MARI-B", "MISI-A","MISI-B","MONR-A","MORD-A","PIAN-A","POLL-A","RENN-A" ), cex.axis=0.4,las=2)

mtext("Autumn", side = 1, line = 4, adj = 0.170, cex=1, mgp=c(3,3,0))

mtext("Spring", side = 1, line = 4, adj = 0.800, cex=1, mgp=c(3,3,0))

dev.off()

tiff("rpoB_Richness_RH_season_genotype.tiff", res=600, width=15, height=15, unit="cm",compression="lzw")

boxplot(npshannon~Pop*Season, data=data_RH, ylab = "Richness", xlab="",main = "",

cex.main=1, cex.axis=0.8, las=2, at = c(1,2,3,4,5,6,7,8,9,10,11,12,13,14,15,16,17,18,19,20,21,22,23,24,25,26 ,29,30,31,32,33,34,35,36,37,38,39,40,41,42,43,44,45,46,47,48,49,50,51,52,53,54), axes=F)

box()

```
axis(2, at=seq(0,30,5), seq(0,30,5), las=1)

axis(1, at=c(1,2,3,4,5,6,7,8,9,10,11,12,13,14,15,16,17,18,19,20,21,22,23,24,25,26
,29,30,31,32,33,34,35,36,37,38,39,40,41,42,43,44,45,46,47,48,49,50,51,52,53,54),

labels = c("ALTO-A","APIG-A","BISA-A","BOEL-A","BOLO-A","BRUZ-A","CAST-A", "CAST-B","CAST-C","COLL-A","CORL-A","FOUG-A","GRAT-A","GRAT-B","ISNE-A","ISNE-B","LASC-A","MARI-A","MARI-B", "MISI-A","MISI-B","MONR-A","MORD-A","PIAN-A","POLL-A","RENN-A","ALTO-A","APIG-A","BISA-A","BOEL-A","BOLO-A","BRUZ-A","CAST-A", "CAST-B","CAST-C","COLL-A","CORL-A","FOUG-A","GRAT-A","GRAT-B","ISNE-A","ISNE-B","LASC-A","MARI-A","MARI-B", "MISI-A","MISI-B","MONR-A","MORD-A","PIAN-A","POLL-A","RENN-A" ), cex.axis=0.4,las=2)

mtext("Autumn", side = 1, line = 4, adj = 0.170, cex=1, mgp=c(3,3,0))

mtext("Spring", side = 1, line = 4, adj = 0.800, cex=1, mgp=c(3,3,0))

dev.off()


# Finley Wilkinson Regression

#Formating

df1 = data.frame(matrix(ncol=1,nrow=1, dimnames=list("AT","Env")))

for (pop in unique(BLUPS_R_AT$Pop)){

  value <- BLUPS_R_AT$npshannon[which(BLUPS_R_AT$Pop==pop)]

  df1[,pop] <- value

}

df1["AT","Env"] <- "AT"


df2 = data.frame(matrix(ncol=1,nrow=1, dimnames=list("SP","Env")))

for (pop in unique(BLUPS_R_SP$Pop)){

  value <- BLUPS_R_SP$npshannon[which(BLUPS_R_SP$Pop==pop)]

  df2[,pop] <- value

}

df2["SP","Env"] <- "SP"


common_cols <- intersect(colnames(df1), colnames(df2)) # Pourquoi difference de colonnes ?!

df3 <- rbind(df1[,common_cols],df2[,common_cols])

# Means

df3$Mean <- rowMeans (df3 %>% select(-Env))
```

```
df3 <- rbind(df3, c("Mean", colMeans(df3 %>% select(-Env))))
```

**#Environmental index**

```
df3$Mean <- as.numeric(df3$Mean)
df3$Env_index <- df3$Mean - df3$Mean[nrow(df3)]
```

**#suppression of means**

```
df3 = subset(df3, select=-c(Mean))
df3<- df3 [-3,]
```

**#reshaping of the data set**

```
df4<- reshape2::melt(df3, id.vars = c("Env", "Env_index"))
colnames(df4)[3]<- c("Pop") #renaming the colomns
colnames(df4)[4]<- c("Shannon")
df4 <- df4 %>% group_by(Pop)

df4$Shannon <- as.numeric(df4$Shannon)

tiff("rpoB_Shannon_R_Finley_wilkinson_BLUPS.tiff", res=600, width=15, height=15, unit="cm",compression="lzw")
p<-ggplot(df4)+
  geom_line(aes(x=Env_index, y=Shannon, group=Pop, color=Pop))+
  geom_point(aes(x=Env_index, y=Shannon, group=Pop, color=Pop))+
  #theme(legend.position = "none")+
  scale_y_continuous(name="Shannon's Index", limits=c(-0.2,3.3,0.5 ))
  p
dev.off()
```

**#formating**

```
df1 = data.frame(matrix(ncol=1,nrow=1, dimnames=list("AT","Env")))
for (pop in unique(BLUPS_RH_AT$Pop)){
  value <- BLUPS_RH_AT$npshannon[which(BLUPS_RH_AT$Pop==pop)]
```

```
  df1[,pop] <- value
}
df1["AT","Env"] <- "AT"

df2 = data.frame(matrix(ncol=1,nrow=1, dimnames=list("SP","Env")))
for (pop in unique(BLUPS_RH_SP$Pop)){
  value <- BLUPS_RH_SP$npshannon[which(BLUPS_RH_SP$Pop==pop)]
  df2[,pop] <- value
}
df2["SP","Env"] <- "SP"

common_cols <- intersect(colnames(df1), colnames(df2)) # Pourquoi difference de colonnes ?!
df3 <- rbind(df1[,common_cols],df2[,common_cols])

# means
df3$Mean <- rowMeans (df3 %>% select(-Env))
df3 <- rbind(df3, c("Mean", colMeans(df3 %>% select(-Env))))

#Environmental index
df3$Mean <- as.numeric(df3$Mean)
df3$Env_index <- df3$Mean - df3$Mean[nrow(df3)]

#suppression of means
df3 = subset(df3, select=-c(Mean))
df3<- df3 [-3,]

#reshaping of the data set
df4<- reshape2::melt(df3, id.vars = c("Env", "Env_index"))
colnames(df4)[3]<- c("Pop") #renaming the colomns
colnames(df4)[4]<- c("Shannon")
df4 <- df4 %>% group_by(Pop)
df4$Shannon <- as.numeric(df4$Shannon)
```

```
tiff("rpoB_Shannon_RH_Finley_wilkinson_BLUPS.tiff", res=600, width=15, height=15, unit="cm",compression="lzw")
p<-ggplot(df4)+
  geom_line(aes(x=Env_index, y=Shannon, group=Pop, color=Pop))+
  geom_point(aes(x=Env_index, y=Shannon, group=Pop, color=Pop))+
  #theme(legend.position = "none")+
  scale_y_continuous(name="Shannon's Index", limits=c(-0.2,3.3,0.5 ))
  p
dev.off()
```

**#R AT**

```
F_stat <- lmer(data_R_AT$npshannon~Pop +(1|rep), data = data_R_AT)
summary(F_stat)
lsmeans(F_stat, pairwise~Pop)
Anova(F_stat)

F_stat <- lmer(data_R_AT$sobs~Pop +(1|rep), data = data_R_AT)
summary(F_stat)
lsmeans(F_stat, pairwise~Pop)
Anova(F_stat)

F_stat <- lmer(data_R_AT$Axis.1~Pop +(1|rep), data = data_R_AT)
summary(F_stat)
lsmeans(F_stat, pairwise~Pop)
Anova(F_stat)

F_stat <- lmer(data_R_AT$Axis.2~Pop +(1|rep), data = data_R_AT)
summary(F_stat)
lsmeans(F_stat, pairwise~Pop)
Anova(F_stat)
```

**#R SP**

```
F_stat <- lmer(data_R_SP$npshannon~Pop +(1|rep), data = data_R_SP)
summary(F_stat)
lsmeans(F_stat, pairwise~Pop)
Anova(F_stat)

F_stat <- lmer(data_R_SP$sobs~Pop +(1|rep), data = data_R_SP)
summary(F_stat)
lsmeans(F_stat, pairwise~Pop)
Anova(F_stat)

F_stat <- lmer(data_R_SP$Axis.1~Pop +(1|rep), data = data_R_SP)
summary(F_stat)
lsmeans(F_stat, pairwise~Pop)
Anova(F_stat)

F_stat <- lmer(data_R_SP$Axis.2~Pop +(1|rep), data = data_R_SP)
summary(F_stat)
lsmeans(F_stat, pairwise~Pop)
Anova(F_stat)
```

**#RH AT**

```
F_stat <- lmer(data_RH_AT$npshannon~Pop +(1|rep), data = data_RH_AT)
summary(F_stat)
lsmeans(F_stat, pairwise~Pop)
Anova(F_stat)

F_stat <- lmer(data_RH_AT$sobs~Pop +(1|rep), data = data_RH_AT)
summary(F_stat)
lsmeans(F_stat, pairwise~Pop)
Anova(F_stat)
```

```
F_stat <- lmer(data_RH_AT$Axis.1~Pop +(1|rep), data = data_RH_AT)
summary(F_stat)
lsmeans(F_stat, pairwise~Pop)
Anova(F_stat)

F_stat <- lmer(data_RH_AT$Axis.2~Pop +(1|rep), data = data_RH_AT)
summary(F_stat)
lsmeans(F_stat, pairwise~Pop)
Anova(F_stat)
```

**#RH SP**

```
F_stat <- lmer(data_RH_SP$npshannon~Pop +(1|rep), data = data_RH_SP)
summary(F_stat)
lsmeans(F_stat, pairwise~Pop)
Anova(F_stat)

F_stat <- lmer(data_RH_SP$sobs~Pop +(1|rep), data = data_RH_SP)
summary(F_stat)
lsmeans(F_stat, pairwise~Pop)
Anova(F_stat)

F_stat <- lmer(data_RH_SP$Axis.1~Pop +(1|rep), data = data_RH_SP)
summary(F_stat)
lsmeans(F_stat, pairwise~Pop)
Anova(F_stat)

F_stat <- lmer(data_RH_SP$Axis.2~Pop +(1|rep), data = data_RH_SP)
summary(F_stat)
lsmeans(F_stat, pairwise~Pop)
Anova(F_stat)
```

**# Scripts used for calculating Hereditability**

### use the diversity measures to calculate broad-sense heritability

library(lme4)

**#### Shannon's Index ####**

**# run the model**

**# R AT**

H1 = lmer(npshannon~(1|Pop) + (1|rep), data=data_R_AT, REML = TRUE)

summary(H1)

plop <- as.data.frame(VarCorr(H1))

v <- plop[which(plop$grp=='Pop'),"vcov"]

r <- plop[which(plop$grp=='Residual'),"vcov"]

h_R_AT <- v/((v)+(r)/6)

h_R_AT

**# R SP**

H2 = lmer(npshannon~(1|Pop) + (1|rep), data=data_R_SP, REML = TRUE)

summary(H2)

plop <- as.data.frame(VarCorr(H2))

v <- plop[which(plop$grp=='Pop'),"vcov"]

r <- plop[which(plop$grp=='Residual'),"vcov"]

h_R_SP <- v/((v)+(r)/6)

h_R_SP

**# RH AT**

H3 = lmer(npshannon~(1|Pop) + (1|rep), data=data_RH_AT, REML = TRUE)

```
summary(H3)

plop <- as.data.frame(VarCorr(H3))
v <- plop[which(plop$grp=='Pop'),"vcov"]
r <- plop[which(plop$grp=='Residual'),"vcov"]

h_RH_AT <- v/((v)+(r)/6)
h_RH_AT

# RH SP
H4 = lmer(npshannon~(1|Pop) + (1|rep), data=data_RH_SP, REML = TRUE)

summary(H4)

plop <- as.data.frame(VarCorr(H4))
v <- plop[which(plop$grp=='Pop'),"vcov"]
r <- plop[which(plop$grp=='Residual'),"vcov"]

h_RH_SP <- v/((v)+(r)/6)
h_RH_SP

#### Richness ####
# run the model
# R AT
H1 = lmer(sobs~(1|Pop) + (1|rep), data=data_R_AT, REML = TRUE)

summary(H1)

plop <- as.data.frame(VarCorr(H1))
v <- plop[which(plop$grp=='Pop'),"vcov"]
r <- plop[which(plop$grp=='Residual'),"vcov"]
```

```
h_R_AT <- v/((v)+(r)/6)
h_R_AT
```

**# R SP**

```
H2 = lmer(sobs~(1|Pop) + (1|rep), data=data_R_SP, REML = TRUE)

summary(H2)

plop <- as.data.frame(VarCorr(H2))
v <- plop[which(plop$grp=='Pop'),"vcov"]
r <- plop[which(plop$grp=='Residual'),"vcov"]

h_R_SP <- v/((v)+(r)/6)
h_R_SP
```

**# RH AT**

```
H3 = lmer(sobs~(1|Pop) + (1|rep), data=data_RH_AT, REML = TRUE)

summary(H3)

plop <- as.data.frame(VarCorr(H3))
v <- plop[which(plop$grp=='Pop'),"vcov"]
r <- plop[which(plop$grp=='Residual'),"vcov"]

h_RH_AT <- v/((v)+(r)/6)
h_RH_AT
```

**# RH SP**

```
H4 = lmer(sobs~(1|Pop) + (1|rep), data=data_RH_SP, REML = TRUE)

summary(H4)
```

```
plop <- as.data.frame(VarCorr(H4))
v <- plop[which(plop$grp=='Pop'),"vcov"]
r <- plop[which(plop$grp=='Residual'),"vcov"]

h_RH_SP <- v/((v)+(r)/6)
h_RH_SP
```

**#### PCoA1 ####**

**# run the model**

**# R AT**

```
H1 = lmer(Axis.1~(1|Pop) + (1|rep), data=data_R_AT, REML = TRUE)

summary(H1)

plop <- as.data.frame(VarCorr(H1))
v <- plop[which(plop$grp=='Pop'),"vcov"]
r <- plop[which(plop$grp=='Residual'),"vcov"]

h_R_AT <- v/((v)+(r)/6)
h_R_AT

# R SP
H2 = lmer(Axis.1~(1|Pop) + (1|rep), data=data_R_SP, REML = TRUE)

summary(H2)

plop <- as.data.frame(VarCorr(H2))
v <- plop[which(plop$grp=='Pop'),"vcov"]
r <- plop[which(plop$grp=='Residual'),"vcov"]

h_R_SP <- v/((v)+(r)/6)
```

```
h_R_SP

# RH AT
H3 = lmer(Axis.1~(1|Pop) + (1|rep), data=data_RH_AT, REML = TRUE)

summary(H3)

plop <- as.data.frame(VarCorr(H3))
v <- plop[which(plop$grp=='Pop'),"vcov"]
r <- plop[which(plop$grp=='Residual'),"vcov"]

h_RH_AT <- v/((v)+(r)/6)
h_RH_AT

# RH SP
H4 = lmer(Axis.1~(1|Pop) + (1|rep), data=data_RH_SP, REML = TRUE)

summary(H4)

plop <- as.data.frame(VarCorr(H4))
v <- plop[which(plop$grp=='Pop'),"vcov"]
r <- plop[which(plop$grp=='Residual'),"vcov"]

h_RH_SP <- v/((v)+(r)/6)
h_RH_SP

#### PCoA2 ####
# run the model
# R AT
H1 = lmer(Axis.2~(1|Pop) + (1|rep), data=data_R_AT, REML = TRUE)

summary(H1)
```

```
plop <- as.data.frame(VarCorr(H1))
v <- plop[which(plop$grp=='Pop'),"vcov"]
r <- plop[which(plop$grp=='Residual'),"vcov"]

h_R_AT <- v/((v)+(r)/6)
h_R_AT
```

**# R SP**

```
H2 = lmer(Axis.2~(1|Pop) + (1|rep), data=data_R_SP, REML = TRUE)

summary(H2)
plop <- as.data.frame(VarCorr(H2))
v <- plop[which(plop$grp=='Pop'),"vcov"]
r <- plop[which(plop$grp=='Residual'),"vcov"]

h_R_SP <- v/((v)+(r)/6)
h_R_SP
```

**# RH AT**

```
H3 = lmer(Axis.2~(1|Pop) + (1|rep), data=data_RH_AT, REML = TRUE)

summary(H3)

plop <- as.data.frame(VarCorr(H3))
v <- plop[which(plop$grp=='Pop'),"vcov"]
r <- plop[which(plop$grp=='Residual'),"vcov"]

h_RH_AT <- v/((v)+(r)/6)
h_RH_AT
```

**# RH SP**

```
H4 = lmer(Axis.2~(1|Pop) + (1|rep), data=data_RH_SP, REML = TRUE)

summary(H4)

plop <- as.data.frame(VarCorr(H4))
v <- plop[which(plop$grp=='Pop'),"vcov"]
r <- plop[which(plop$grp=='Residual'),"vcov"]

h_RH_SP <- v/((v)+(r)/6)
h_RH_SP
```

**############################## tbRDA ####################################**

**### Count ###**

```
tbRDA_count_R_AT=shared_1000_abd1_R_AT
tbRDA_count_RH_AT=shared_1000_abd1_RH_AT
tbRDA_count_RH_SP=shared_1000_abd1_RH_SP
tbRDA_count_R_SP=shared_1000_abd1_R_SP
```

**### Design ###**

**# R AT**

```
tbRDA_design_R_AT = data.frame(matrix(ncol=5,nrow=nrow(tbRDA_count_R_AT)))
colnames(tbRDA_design_R_AT) <- c("Sample",'Rep',"Genotype","Compartment","Season")
tbRDA_design_R_AT$Sample <- tbRDA_count_R_AT$Group
tbRDA_design_R_AT$Genotype <- substr(tbRDA_count_R_AT$Group,1,6)
tbRDA_design_R_AT$Season <- substr(tbRDA_count_R_AT$Group,8,9)
tbRDA_design_R_AT$Compartment <- substr(tbRDA_count_R_AT$Group,11,11)
tbRDA_design_R_AT$Rep <- substr(tbRDA_count_R_AT$Group,13,13)
```

**# R SP**

```
tbRDA_design_R_SP = data.frame(matrix(ncol=4,nrow=nrow(tbRDA_count_R_SP)))
```

colnames(tbRDA_design_R_SP) <- c("Sample","Genotype","Compartment","Season")

tbRDA_design_R_SP$Sample <- tbRDA_count_R_SP$Group

tbRDA_design_R_SP$Genotype <- substr(tbRDA_count_R_SP$Group,1,6)

tbRDA_design_R_SP$Season <- substr(tbRDA_count_R_SP$Group,8,9)

tbRDA_design_R_SP$Compartment <- substr(tbRDA_count_R_SP$Group,11,11)

tbRDA_design_R_SP$Rep <- substr(tbRDA_count_R_SP$Group,13,13)

**# RH AT**

tbRDA_design_RH_AT = data.frame(matrix(ncol=4,nrow=nrow(tbRDA_count_RH_AT)))

colnames(tbRDA_design_RH_AT) <- c("Sample","Genotype","Compartment","Season")

tbRDA_design_RH_AT$Sample <- tbRDA_count_RH_AT$Group

tbRDA_design_RH_AT$Genotype <- substr(tbRDA_count_RH_AT$Group,1,6)

tbRDA_design_RH_AT$Season <- substr(tbRDA_count_RH_AT$Group,8,9)

tbRDA_design_RH_AT$Compartment <- substr(tbRDA_count_RH_AT$Group,11,11)

tbRDA_design_RH_AT$Rep <- substr(tbRDA_count_RH_AT$Group,14,14)

**# RH SP**

tbRDA_design_RH_SP = data.frame(matrix(ncol=4,nrow=nrow(tbRDA_count_RH_SP)))

colnames(tbRDA_design_RH_SP) <- c("Sample","Genotype","Compartment","Season")

tbRDA_design_RH_SP$Sample <- tbRDA_count_RH_SP$Group

tbRDA_design_RH_SP$Genotype <- substr(tbRDA_count_RH_SP$Group,1,6)

tbRDA_design_RH_SP$Season <- substr(tbRDA_count_RH_SP$Group,8,9)

tbRDA_design_RH_SP$Compartment <- substr(tbRDA_count_RH_SP$Group,11,11)

tbRDA_design_RH_SP$Rep <- substr(tbRDA_count_RH_SP$Group,14,14)

tbRDA_count_R_AT <- data.frame(tbRDA_count_R_AT[,-1], row.names=tbRDA_count_R_AT[,1])

tbRDA_count_R_SP <- data.frame(tbRDA_count_R_SP[,-1], row.names=tbRDA_count_R_SP[,1])

tbRDA_count_RH_AT <- data.frame(tbRDA_count_RH_AT[,-1], row.names=tbRDA_count_RH_AT[,1])

tbRDA_count_RH_SP <- data.frame(tbRDA_count_RH_SP[,-1], row.names=tbRDA_count_RH_SP[,1])

**### tbRDA ###**

**# R_AT**

tbRDA_count_R_AT = t(tbRDA_count_R_AT)

tbRDA_count_R_AT_norm <- t(t(tbRDA_count_R_AT)/colSums(tbRDA_count_R_AT,na=T))*1000

nrow(tbRDA_count_R_AT_norm)

tbRDA_count_R_AT_norm[is.na(tbRDA_count_R_AT_norm)] <- 0

threshold <- 1

tbRDA_count_R_AT_norm <- tbRDA_count_R_AT_norm[ apply(tbRDA_count_R_AT_norm,1,max) > threshold, ]

nrow(tbRDA_count_R_AT_norm)

tbRDA_count_R_AT_norm <- t(tbRDA_count_R_AT_norm)

tbRDA_count_R_AT_norm = tbRDA_count_R_AT_norm[order(tbRDA_count_R_AT_norm[,1]),]

dim(tbRDA_count_R_AT_norm)

tbRDA_count_R_AT_log<- log1p(tbRDA_count_R_AT_norm) # species data are in percentage scale which is strongly rightskewed, better to transform them

tbRDA_count_R_AT_HELL <- decostand(tbRDA_count_R_AT_log, 'hell') # we are planning to do tb-RDA, this is Hellinger pre-transformation

**# Make the RDA on the Hellinger matrix**

**# check that Hellinger matrix and design have same N° of row**

nrow(tbRDA_design_R_AT)

nrow(tbRDA_count_R_AT_HELL)

**# perform rda**

tbRDA <- rda (tbRDA_count_R_AT_HELL~ (Genotype) + Condition(Rep),data = tbRDA_design_R_AT) # calculate tb-RDA with one explanatory variable (genotype in this case)

tbRDA

**# Estimate the explicative capacity of the RDA thanks to the % of the constrained variance of the analysis**

MVA.synt(tbRDA)

**# Perform a permutation test to assign a P-value to the explained variance**

```
MVA.anova(tbRDA)
```

**# Effect of one factor or a combination of factors**

```
pairwise.factorfit(tbRDA,tbRDA_design_R_AT$genotype,data = tbRDA_design_R_AT)
```

**# graphe of all genotypes**

```
graph<-MVA.plot(tbRDA)
```

**# graph bases on genotypes**

```
tiff("./tb-RDA_R_AT.tiff", res=600, width=15, height=15, unit="cm",compression="lzw")

MVA.plot(tbRDA, main ="",main.pos="topright", ylim=c(-1,1), xlim=c(-1,1),xlab=" Axis 1 (18.86 %)", ylab="Axis 2 (11.01

 %)",points=TRUE, main.cex=0.8, cex=0.5,labels=rownames(tbRDA), pch=16, col="black", fac=tbRDA_design_R_AT$Genotype)#tbRDA,

dev.off()
```

**# R_SP**

```
tbRDA_count_R_SP = t(tbRDA_count_R_SP)

tbRDA_count_R_SP_norm <- t(t(tbRDA_count_R_SP)/colSums(tbRDA_count_R_SP,na=T))*1000

nrow(tbRDA_count_R_SP_norm)

tbRDA_count_R_SP_norm[is.na(tbRDA_count_R_SP_norm)] <- 0


threshold <- 1

tbRDA_count_R_SP_norm <- tbRDA_count_R_SP_norm[ apply(tbRDA_count_R_SP_norm,1,max, na.rm=T) > threshold, ]

nrow(tbRDA_count_R_SP_norm)


tbRDA_count_R_SP_norm <- t(tbRDA_count_R_SP_norm)

tbRDA_count_R_SP_norm = tbRDA_count_R_SP_norm[order(tbRDA_count_R_SP_norm[,1]),]

dim(tbRDA_count_R_SP_norm)
```

tbRDA_count_R_SP_log<- log1p(tbRDA_count_R_SP_norm)  # species data are in percentage scale which is strongly rightskewed, better to transform them

tbRDA_count_R_SP_HELL <- decostand(tbRDA_count_R_SP_log, 'hell')  # we are planning to do tb-RDA, this is Hellinger pre-transformation

**# Make the RDA on the Hellinger matrix**

**# check that Hellinger matrix and design have same N° of row**

nrow(tbRDA_design_R_SP)

nrow(tbRDA_count_R_SP_HELL)

### perform rda

tbRDA <- rda (tbRDA_count_R_SP_HELL~ (Genotype) + Condition(Rep),data = tbRDA_design_R_SP)  # calculate tb-RDA with one explanatory variable (genotype in this case)

tbRDA

**# Estimate the explicative capacity of the RDA thanks to the % of the constrained variance of the analysis**

MVA.synt(tbRDA)

**# Perform a permutation test to assign a P-value to the explained variance**

MVA.anova(tbRDA)

**# Effect of one factor or a combination of factors**

pairwise.factorfit(tbRDA,tbRDA_design_R_SP$genotype,data = tbRDA_design_R_SP)

**# graphe of all genotypes**

graph<-MVA.plot(tbRDA)

**# graph bases on genotypes**

tiff("./tb-RDA_R_SP.tiff", res=600, width=15, height=15, unit="cm",compression="lzw")

MVA.plot(tbRDA, main ="",main.pos="topright", ylim=c(-1,1), xlim=c(-1,1),xlab=" Axis 1 (34.80 %)", ylab="Axis 2 (9.78

%)",points=TRUE, main.cex=0.8, cex=0.5,labels=rownames(tbRDA), pch=16, col="black", fac=tbRDA_design_R_SP$Genotype)#tbRDA,

dev.off()

**# RH_SP**

tbRDA_count_RH_SP = t(tbRDA_count_RH_SP)

tbRDA_count_RH_SP_norm <- t(t(tbRDA_count_RH_SP)/colSums(tbRDA_count_RH_SP,na=T))*1000

nrow(tbRDA_count_RH_SP_norm)

tbRDA_count_RH_SP_norm[is.na(tbRDA_count_RH_SP_norm)] <- 0

threshold <- 1

tbRDA_count_RH_SP_norm <- tbRDA_count_RH_SP_norm[ apply(tbRDA_count_RH_SP_norm,1,max, na.rm=T) > threshold, ]

nrow(tbRDA_count_RH_SP_norm)

tbRDA_count_RH_SP_norm <- t(tbRDA_count_RH_SP_norm)

tbRDA_count_RH_SP_norm = tbRDA_count_RH_SP_norm[order(tbRDA_count_RH_SP_norm[,1]),]

dim(tbRDA_count_RH_SP_norm)

tbRDA_count_RH_SP_log<- log1p(tbRDA_count_RH_SP_norm)  # species data are in percentage scale which is strongly rightskewed, better to transform them

tbRDA_count_RH_SP_HELL <- decostand(tbRDA_count_RH_SP_log, 'hell')  # we are planning to do tb-RDA, this is Hellinger pre-transformation

**# Make the RDA on the Hellinger matrix**

**# check that Hellinger matrix and design have same N° of row**

nrow(tbRDA_design_RH_SP)

nrow(tbRDA_count_RH_SP_HELL)

**# perform rda**

tbRDA <- rda (tbRDA_count_RH_SP_HELL~ (Genotype) + Condition(Rep),data = tbRDA_design_RH_SP)  # calculate tb-RDA with one explanatory variable (genotype in this case) tbRDA

**# Estimate the explicative capacity of the RDA thanks to the % of the constrained variance of the analysis**

MVA.synt(tbRDA)

**# Perform a permutation test to assign a P-value to the explained variance**

MVA.anova(tbRDA)

**# Effect of one factor or a combination of factors**

pairwise.factorfit(tbRDA,tbRDA_design_RH_SP$genotype,data = tbRDA_design_RH_SP)

**# graphe of all genotypes**

graph<-MVA.plot(tbRDA)

**# graph bases on genotypes**

tiff("./tb-RDA_RH_SP.tiff", res=600, width=15, height=15, unit="cm",compression="lzw")

MVA.plot(tbRDA, main ="",main.pos="topright", ylim=c(-1,1), xlim=c(-1.3,0.5),xlab=" Axis 1 (12.63 %)", ylab="Axis 2 (9.74

%)",points=TRUE, main.cex=0.8, cex=0.5,labels=rownames(tbRDA), pch=16, col="black", fac=tbRDA_design_RH_SP$Genotype)#tbRDA,

dev.off()

**# RH_AT**

tbRDA_count_RH_AT = t(tbRDA_count_RH_AT)

tbRDA_count_RH_AT_norm <- t(t(tbRDA_count_RH_AT)/colSums(tbRDA_count_RH_AT,na=T))*1000

nrow(tbRDA_count_RH_AT_norm)

tbRDA_count_RH_AT_norm[is.na(tbRDA_count_RH_AT_norm)] <- 0

threshold <- 1

tbRDA_count_RH_AT_norm <- tbRDA_count_RH_AT_norm[ apply(tbRDA_count_RH_AT_norm,1,max, na.rm=T) > threshold, ] nrow(tbRDA_count_RH_AT_norm)

tbRDA_count_RH_AT_norm <- t(tbRDA_count_RH_AT_norm)

tbRDA_count_RH_AT_norm = tbRDA_count_RH_AT_norm[order(tbRDA_count_RH_AT_norm[,1]),]

dim(tbRDA_count_RH_AT_norm)

tbRDA_count_RH_AT_log<- log1p(tbRDA_count_RH_AT_norm)  # species data are in percentage scale which is strongly rightskewed, better to transform them

tbRDA_count_RH_AT_HELL <- decostand(tbRDA_count_RH_AT_log, 'hell')  # we are planning to do tb-RDA, this is Hellinger pre-transformation

**# Make the RDA on the Hellinger matrix**

**# check that Hellinger matrix and design have same N° of row**

nrow(tbRDA_design_RH_AT)

nrow(tbRDA_count_RH_AT_HELL)

**# perform rda**

tbRDA <- rda (tbRDA_count_RH_AT_HELL~ (Genotype) + Condition(Rep),data = tbRDA_design_RH_AT)  # calculate tb-RDA with one explanatory variable (genotype in this case)

tbRDA

**# Estimate the explicative capacity of the RDA thanks to the % of the constrained variance of the analysis**

MVA.synt(tbRDA)

**# Perform a permutation test to assign a P-value to the explained variance**

MVA.anova(tbRDA)

**# Effect of one factor or a combination of factors**

pairwise.factorfit(tbRDA,tbRDA_design_RH_AT$genotype,data = tbRDA_design_RH_AT)

**# graphe of all genotypes**

graph<-MVA.plot(tbRDA)

**# graph bases on genotypes**

```
tiff("./tb-RDA_RH_AT.tiff", res=600, width=15, height=15, unit="cm",compression="lzw")

MVA.plot(tbRDA, main ="",main.pos="topright", ylim=c(-1,1), xlim=c(-0.7,1.2),xlab=" Axis 1 (13.92 %)", ylab="Axis 2 (9.84

 %)",points=TRUE, main.cex=0.8, cex=0.5,labels=rownames(tbRDA), pch=16, col="black", fac=tbRDA_design_RH_AT$Genotype)#tbRDA,

dev.off()
```

#### The following scripts have been used for GEA analysis

**############# AT RH ###########**

**# Singular value decomposition (SVD) on matric of allele frequency**

```
om.mat=as.matrix(read.table("./Dataset13_AT_RH_sub1_1_mat_omega.out"))

listpop=as.data.frame(matrix(NA, ncol=2, nrow=26))

colnames(listpop)=c("pop", "color")

listpop$pop=c("ALTO-A","APIG-A","BISA-A","BOEL-A","BOLO-A","BRUZ-A","CAST-A","CAST-B","CAST-C","COLL-A","CORL-A","FOUG-A","GRAT-A","GRAT-B","ISNE-A","ISNE-B","LASC-A","MARI-A","MARI-B","MISI-A","MISI-B","MONR-A","MORD-A","PIAN-A","POLL-A","RENN-A")

listpop$color=c("orange", "deepskyblue4", "deepskyblue4", "deepskyblue4", "deepskyblue4",

    "deepskyblue4", "deepskyblue4", "deepskyblue4", "deepskyblue4", "deepskyblue4",

    "orange", "orange", "deepskyblue4", "deepskyblue4", "orange", "orange", "orange",

    "deepskyblue4", "deepskyblue4", "deepskyblue4", "deepskyblue4","orange",

    "deepskyblue4", "deepskyblue4", "deepskyblue4", "deepskyblue4")
```

**# Singular value decomposition**

```
om.svd=svd(om.mat)
```

**# eigenvalues of the diagonal of the variance**

```
om.eig=om.svd$d**2 # <=> om.eig=om.svd$d^2

perc.var=100*om.eig/sum(om.eig)
```

```
PC=cbind(listpop, om.svd$u[,1:2])
colnames(PC)=c("population", "color","PC1", "PC2")
write.table(PC, "./Dataset15.txt", sep="\t")
```

**### Concatenate Bayesian Factor (BFis in dB) and XtX index from different parallised anaylses on 20 sub-samples###**

**# Open files with identity of markers within each sub-samples (sub1 to sub19)**

```
for(i in 1:20) {
  assign( paste("mrk", i, sep=""),
          read.table(paste("/groups/holoe2plant/Loeiz/R_bis/Dataset10.snpdet.sub", i, sep=""), h=F) )
}
```

**### concatenation bayes factors files**

```
Dataset5 <- read.table("/groups/holoe2plant/Loeiz/R_rpob/BLUPS_RH_AT.txt", h=T)
traits=colnames(Dataset5[,-1])
```

**# first repeat**

```
for(i in 1:20) {
     assign( paste("sub", i, "_1", sep=""),
             read.table(paste("./Dataset13_AT_RH_sub",i,"_1_summary_betai_reg.out", sep=""), h=T) )
}
```

**# second repeat**

```
for(i in 1:20) {
     assign( paste("sub", i, "_2", sep=""),
             read.table(paste("./Dataset13_AT_RH_sub",i,"_2_summary_betai_reg.out", sep=""), h=T) )
}
```

**# Third repeat**

```
for(i in 1:20) {
     assign( paste("sub", i, "_3", sep=""),
```

```
read.table(paste("./Dataset13_AT_RH_sub",i,"_3_summary_betai_reg.out", sep=""), h=T) )
}

for (i in 1:length(traits)) {

sub1=cbind(sub1_1[sub1_1$COVARIABLE==i,],sub1_2[sub1_2$COVARIABLE==i,],sub1_3[sub1_3$COVARIABLE==i,])

sub2=cbind(sub2_1[sub2_1$COVARIABLE==i,],sub2_2[sub2_2$COVARIABLE==i,],sub2_3[sub2_3$COVARIABLE==i,])

sub3=cbind(sub3_1[sub3_1$COVARIABLE==i,],sub3_2[sub3_2$COVARIABLE==i,],sub3_3[sub3_3$COVARIABLE==i,])

sub4=cbind(sub4_1[sub4_1$COVARIABLE==i,],sub4_2[sub4_2$COVARIABLE==i,],sub4_3[sub4_3$COVARIABLE==i,])

sub5=cbind(sub5_1[sub5_1$COVARIABLE==i,],sub5_2[sub5_2$COVARIABLE==i,],sub5_3[sub5_3$COVARIABLE==i,])

sub6=cbind(sub6_1[sub6_1$COVARIABLE==i,],sub6_2[sub6_2$COVARIABLE==i,],sub6_3[sub6_3$COVARIABLE==i,])

sub7=cbind(sub7_1[sub7_1$COVARIABLE==i,],sub7_2[sub7_2$COVARIABLE==i,],sub7_3[sub7_3$COVARIABLE==i,])

sub8=cbind(sub8_1[sub8_1$COVARIABLE==i,],sub8_2[sub8_2$COVARIABLE==i,],sub8_3[sub8_3$COVARIABLE==i,])

sub9=cbind(sub9_1[sub9_1$COVARIABLE==i,],sub9_2[sub9_2$COVARIABLE==i,],sub9_3[sub9_3$COVARIABLE==i,])

sub10=cbind(sub10_1[sub10_1$COVARIABLE==i,],sub10_2[sub10_2$COVARIABLE==i,],sub10_3[sub10_3$COVARIABLE==i,])

sub11=cbind(sub11_1[sub11_1$COVARIABLE==i,],sub11_2[sub11_2$COVARIABLE==i,],sub11_3[sub11_3$COVARIABLE==i,])

sub12=cbind(sub12_1[sub12_1$COVARIABLE==i,],sub12_2[sub12_2$COVARIABLE==i,],sub12_3[sub12_3$COVARIABLE==i,])
```

```
sub13=cbind(sub13_1[sub13_1$COVARIABLE==i,],sub13_2[sub13_2$COVARIABLE==i,],sub13_3[sub13_3$COVARIABLE==i,])

sub14=cbind(sub14_1[sub14_1$COVARIABLE==i,],sub14_2[sub14_2$COVARIABLE==i,],sub14_3[sub14_3$COVARIABLE==i,])

sub15=cbind(sub15_1[sub15_1$COVARIABLE==i,],sub15_2[sub15_2$COVARIABLE==i,],sub15_3[sub15_3$COVARIABLE==i,])

sub16=cbind(sub16_1[sub16_1$COVARIABLE==i,],sub16_2[sub16_2$COVARIABLE==i,],sub16_3[sub16_3$COVARIABLE==i,])

sub17=cbind(sub17_1[sub17_1$COVARIABLE==i,],sub17_2[sub17_2$COVARIABLE==i,],sub17_3[sub17_3$COVARIABLE==i,])

sub18=cbind(sub18_1[sub18_1$COVARIABLE==i,],sub18_2[sub18_2$COVARIABLE==i,],sub18_3[sub18_3$COVARIABLE==i,])

sub19=cbind(sub19_1[sub19_1$COVARIABLE==i,],sub19_2[sub19_2$COVARIABLE==i,],sub19_3[sub19_3$COVARIABLE==i,])

sub20=cbind(sub20_1[sub20_1$COVARIABLE==i,],sub20_2[sub20_2$COVARIABLE==i,],sub20_3[sub20_3$COVARIABLE==i,])


  sub1$mean_M_Pearson=rowMeans(sub1[,c(3,11,19)])
  sub1$mean_BFdB=rowMeans(sub1[,c(5,13,21)])
  sub1$mean_Beta_is=rowMeans(sub1[,c(6,14,22)])
  sub2$mean_M_Pearson=rowMeans(sub2[,c(3,11,19)])
  sub2$mean_BFdB=rowMeans(sub2[,c(5,13,21)])
  sub2$mean_Beta_is=rowMeans(sub2[,c(6,14,22)])
  sub3$mean_M_Pearson=rowMeans(sub3[,c(3,11,19)])
  sub3$mean_BFdB=rowMeans(sub3[,c(5,13,21)])
  sub3$mean_Beta_is=rowMeans(sub3[,c(6,14,22)])
  sub4$mean_M_Pearson=rowMeans(sub4[,c(3,11,19)])
  sub4$mean_BFdB=rowMeans(sub4[,c(5,13,21)])
  sub4$mean_Beta_is=rowMeans(sub4[,c(6,14,22)])
  sub5$mean_M_Pearson=rowMeans(sub5[,c(3,11,19)])
```

```
sub5$mean_BFdB=rowMeans(sub5[,c(5,13,21)])
sub5$mean_Beta_is=rowMeans(sub5[,c(6,14,22)])
sub6$mean_M_Pearson=rowMeans(sub6[,c(3,11,19)])
sub6$mean_BFdB=rowMeans(sub6[,c(5,13,21)])
sub6$mean_Beta_is=rowMeans(sub6[,c(6,14,22)])
sub7$mean_M_Pearson=rowMeans(sub7[,c(3,11,19)])
sub7$mean_BFdB=rowMeans(sub7[,c(5,13,21)])
sub7$mean_Beta_is=rowMeans(sub7[,c(6,14,22)])
sub8$mean_M_Pearson=rowMeans(sub8[,c(3,11,19)])
sub8$mean_BFdB=rowMeans(sub8[,c(5,13,21)])
sub8$mean_Beta_is=rowMeans(sub8[,c(6,14,22)])
sub9$mean_M_Pearson=rowMeans(sub9[,c(3,11,19)])
sub9$mean_BFdB=rowMeans(sub9[,c(5,13,21)])
sub9$mean_Beta_is=rowMeans(sub9[,c(6,14,22)])
sub10$mean_M_Pearson=rowMeans(sub10[,c(3,11,19)])
sub10$mean_BFdB=rowMeans(sub10[,c(5,13,21)])
sub10$mean_Beta_is=rowMeans(sub10[,c(6,14,22)])
sub11$mean_M_Pearson=rowMeans(sub11[,c(3,11,19)])
sub11$mean_BFdB=rowMeans(sub11[,c(5,13,21)])
sub11$mean_Beta_is=rowMeans(sub11[,c(6,14,22)])
sub12$mean_M_Pearson=rowMeans(sub12[,c(3,11,19)])
sub12$mean_BFdB=rowMeans(sub12[,c(5,13,21)])
sub12$mean_Beta_is=rowMeans(sub12[,c(6,14,22)])
sub13$mean_M_Pearson=rowMeans(sub13[,c(3,11,19)])
sub13$mean_BFdB=rowMeans(sub13[,c(5,13,21)])
sub13$mean_Beta_is=rowMeans(sub13[,c(6,14,22)])
sub14$mean_M_Pearson=rowMeans(sub14[,c(3,11,19)])
sub14$mean_BFdB=rowMeans(sub14[,c(5,13,21)])
sub14$mean_Beta_is=rowMeans(sub14[,c(6,14,22)])
sub15$mean_M_Pearson=rowMeans(sub15[,c(3,11,19)])
sub15$mean_BFdB=rowMeans(sub15[,c(5,13,21)])
sub15$mean_Beta_is=rowMeans(sub15[,c(6,14,22)])
```

```
sub16$mean_M_Pearson=rowMeans(sub16[,c(3,11,19)])
sub16$mean_BFdB=rowMeans(sub16[,c(5,13,21)])
sub16$mean_Beta_is=rowMeans(sub16[,c(6,14,22)])
sub17$mean_M_Pearson=rowMeans(sub17[,c(3,11,19)])
sub17$mean_BFdB=rowMeans(sub17[,c(5,13,21)])
sub17$mean_Beta_is=rowMeans(sub17[,c(6,14,22)])
sub18$mean_M_Pearson=rowMeans(sub18[,c(3,11,19)])
sub18$mean_BFdB=rowMeans(sub18[,c(5,13,21)])
sub18$mean_Beta_is=rowMeans(sub18[,c(6,14,22)])
sub19$mean_M_Pearson=rowMeans(sub19[,c(3,11,19)])
sub19$mean_BFdB=rowMeans(sub19[,c(5,13,21)])
sub19$mean_Beta_is=rowMeans(sub19[,c(6,14,22)])
sub20$mean_M_Pearson=rowMeans(sub20[,c(3,11,19)])
sub20$mean_BFdB=rowMeans(sub20[,c(5,13,21)])
sub20$mean_Beta_is=rowMeans(sub20[,c(6,14,22)])

tsub1=cbind(mrk1[,c(1:2)],sub1[,c(25,26,27)])
tsub2=cbind(mrk2[,c(1:2)],sub2[,c(25,26,27)])
tsub3=cbind(mrk3[,c(1:2)],sub3[,c(25,26,27)])
tsub4=cbind(mrk4[,c(1:2)],sub4[,c(25,26,27)])
tsub5=cbind(mrk5[,c(1:2)],sub5[,c(25,26,27)])
tsub6=cbind(mrk6[,c(1:2)],sub6[,c(25,26,27)])
tsub7=cbind(mrk7[,c(1:2)],sub7[,c(25,26,27)])
tsub8=cbind(mrk8[,c(1:2)],sub8[,c(25,26,27)])
tsub9=cbind(mrk9[,c(1:2)],sub9[,c(25,26,27)])
tsub10=cbind(mrk10[,c(1:2)],sub10[,c(25,26,27)])
tsub11=cbind(mrk11[,c(1:2)],sub11[,c(25,26,27)])
tsub12=cbind(mrk12[,c(1:2)],sub12[,c(25,26,27)])
tsub13=cbind(mrk13[,c(1:2)],sub13[,c(25,26,27)])
tsub14=cbind(mrk14[,c(1:2)],sub14[,c(25,26,27)])
tsub15=cbind(mrk15[,c(1:2)],sub15[,c(25,26,27)])
tsub16=cbind(mrk16[,c(1:2)],sub16[,c(25,26,27)])
```

```
  tsub17=cbind(mrk17[,c(1:2)],sub17[,c(25,26,27)])
  tsub18=cbind(mrk18[,c(1:2)],sub18[,c(25,26,27)])
  tsub19=cbind(mrk19[,c(1:2)],sub19[,c(25,26,27)])
  tsub20=cbind(mrk20[,c(1:2)],sub20[,c(25,26,27)])


datafinal=rbind(tsub1,tsub2,tsub3,tsub4,tsub5,tsub6,tsub7,tsub8,tsub9,tsub10,tsub11,tsub12,tsub13,tsub14,tsub15,tsub16,tsub17,tsub18,tsub19,tsub20)
  colnames(datafinal)=c("scaffold","pos","M_Pearson","BFdB","Beta_is")

  name=traits[i]


write.table(datafinal,paste("./dataset16_AT_RH_",name,".txt",sep=""),row.names=F,col.names=T,quote=F)
}
```

**### XtX values concatenation**

```
for(i in 1:20) {
  assign( paste("mrk", i, sep=""),
         read.table(paste("/groups/holoe2plant/Loeiz/R_bis/Dataset10.snpdet.sub", i, sep=""), h=F) )
}
```

**# first repeat**

```
for(i in 1:20) {
    assign( paste("XtX", i, "_1", sep=""),
            read.table(paste("./Dataset13_AT_RH_sub",i,"_1_summary_pi_xtx.out", sep=""), h=T) )
}
```

**# second repeat**

```
for(i in 1:20) {
    assign( paste("XtX", i, "_2", sep=""),
            read.table(paste("./Dataset13_AT_RH_sub",i,"_2_summary_pi_xtx.out", sep=""), h=T) )
```

```
}
```

**# Third repeat**

```
for(i in 1:20) {
    assign( paste("XtX", i, "_3", sep=""),
            read.table(paste("./Dataset13_AT_RH_sub",i,"_3_summary_pi_xtx.out", sep=""), h=T) )
}

XtX1=cbind(XtX1_1,XtX1_2,XtX1_3)
XtX2=cbind(XtX2_1,XtX2_2,XtX2_3)
XtX3=cbind(XtX3_1,XtX3_2,XtX3_3)
XtX4=cbind(XtX4_1,XtX4_2,XtX4_3)
XtX5=cbind(XtX5_1,XtX5_2,XtX5_3)
XtX6=cbind(XtX6_1,XtX6_2,XtX6_3)
XtX7=cbind(XtX7_1,XtX7_2,XtX7_3)
XtX8=cbind(XtX8_1,XtX8_2,XtX8_3)
XtX9=cbind(XtX9_1,XtX9_2,XtX9_3)
XtX10=cbind(XtX10_1,XtX10_2,XtX10_3)
XtX11=cbind(XtX11_1,XtX11_2,XtX11_3)
XtX12=cbind(XtX12_1,XtX12_2,XtX12_3)
XtX13=cbind(XtX13_1,XtX13_2,XtX13_3)
XtX14=cbind(XtX14_1,XtX14_2,XtX14_3)
XtX15=cbind(XtX15_1,XtX15_2,XtX15_3)
XtX16=cbind(XtX16_1,XtX16_2,XtX16_3)
XtX17=cbind(XtX17_1,XtX17_2,XtX17_3)
XtX18=cbind(XtX18_1,XtX18_2,XtX18_3)
XtX19=cbind(XtX19_1,XtX19_2,XtX19_3)
XtX20=cbind(XtX20_1,XtX20_2,XtX20_3)

XtX1$mean_M_P=rowMeans(XtX1[,c(2,9,16)])
XtX1$mean_SD_P=rowMeans(XtX1[,c(3,10,17)])
XtX1$mean_M_XtX=rowMeans(XtX1[,c(4,11,18)])
```

```
XtX1$mean_SD_XtX=rowMeans(XtX1[,c(5,12,19)])
XtX1$mean_XtXst=rowMeans(XtX1[,c(6,13,20)])
XtX1$mean_log10.pval=rowMeans(XtX1[,c(7,14,21)])
XtX2$mean_M_P=rowMeans(XtX2[,c(2,9,16)])
XtX2$mean_SD_P=rowMeans(XtX2[,c(3,10,17)])
XtX2$mean_M_XtX=rowMeans(XtX2[,c(4,11,18)])
XtX2$mean_SD_XtX=rowMeans(XtX2[,c(5,12,19)])
XtX2$mean_XtXst=rowMeans(XtX2[,c(6,13,20)])
XtX2$mean_log10.pval=rowMeans(XtX2[,c(7,14,21)])
XtX3$mean_M_P=rowMeans(XtX3[,c(2,9,16)])
XtX3$mean_SD_P=rowMeans(XtX3[,c(3,10,17)])
XtX3$mean_M_XtX=rowMeans(XtX3[,c(4,11,18)])
XtX3$mean_SD_XtX=rowMeans(XtX3[,c(5,12,19)])
XtX3$mean_XtXst=rowMeans(XtX3[,c(6,13,20)])
XtX3$mean_log10.pval=rowMeans(XtX3[,c(7,14,21)])
XtX4$mean_M_P=rowMeans(XtX4[,c(2,9,16)])
XtX4$mean_SD_P=rowMeans(XtX4[,c(3,10,17)])
XtX4$mean_M_XtX=rowMeans(XtX4[,c(4,11,18)])
XtX4$mean_SD_XtX=rowMeans(XtX4[,c(5,12,19)])
XtX4$mean_XtXst=rowMeans(XtX4[,c(6,13,20)])
XtX4$mean_log10.pval=rowMeans(XtX4[,c(7,14,21)])
XtX5$mean_M_P=rowMeans(XtX5[,c(2,9,16)])
XtX5$mean_SD_P=rowMeans(XtX5[,c(3,10,17)])
XtX5$mean_M_XtX=rowMeans(XtX5[,c(4,11,18)])
XtX5$mean_SD_XtX=rowMeans(XtX5[,c(5,12,19)])
XtX5$mean_XtXst=rowMeans(XtX5[,c(6,13,20)])
XtX5$mean_log10.pval=rowMeans(XtX5[,c(7,14,21)])
XtX6$mean_M_P=rowMeans(XtX6[,c(2,9,16)])
XtX6$mean_SD_P=rowMeans(XtX6[,c(3,10,17)])
XtX6$mean_M_XtX=rowMeans(XtX6[,c(4,11,18)])
XtX6$mean_SD_XtX=rowMeans(XtX6[,c(5,12,19)])
XtX6$mean_XtXst=rowMeans(XtX6[,c(6,13,20)])
```

```
XtX6$mean_log10.pval=rowMeans(XtX6[,c(7,14,21)])
XtX7$mean_M_P=rowMeans(XtX7[,c(2,9,16)])
XtX7$mean_SD_P=rowMeans(XtX7[,c(3,10,17)])
XtX7$mean_M_XtX=rowMeans(XtX7[,c(4,11,18)])
XtX7$mean_SD_XtX=rowMeans(XtX7[,c(5,12,19)])
XtX7$mean_XtXst=rowMeans(XtX7[,c(6,13,20)])
XtX7$mean_log10.pval=rowMeans(XtX7[,c(7,14,21)])
XtX8$mean_M_P=rowMeans(XtX8[,c(2,9,16)])
XtX8$mean_SD_P=rowMeans(XtX8[,c(3,10,17)])
XtX8$mean_M_XtX=rowMeans(XtX8[,c(4,11,18)])
XtX8$mean_SD_XtX=rowMeans(XtX8[,c(5,12,19)])
XtX8$mean_XtXst=rowMeans(XtX8[,c(6,13,20)])
XtX8$mean_log10.pval=rowMeans(XtX8[,c(7,14,21)])
XtX9$mean_M_P=rowMeans(XtX9[,c(2,9,16)])
XtX9$mean_SD_P=rowMeans(XtX9[,c(3,10,17)])
XtX9$mean_M_XtX=rowMeans(XtX9[,c(4,11,18)])
XtX9$mean_SD_XtX=rowMeans(XtX9[,c(5,12,19)])
XtX9$mean_XtXst=rowMeans(XtX9[,c(6,13,20)])
XtX9$mean_log10.pval=rowMeans(XtX9[,c(7,14,21)])
XtX10$mean_M_P=rowMeans(XtX10[,c(2,9,16)])
XtX10$mean_SD_P=rowMeans(XtX10[,c(3,10,17)])
XtX10$mean_M_XtX=rowMeans(XtX10[,c(4,11,18)])
XtX10$mean_SD_XtX=rowMeans(XtX10[,c(5,12,19)])
XtX10$mean_XtXst=rowMeans(XtX10[,c(6,13,20)])
XtX10$mean_log10.pval=rowMeans(XtX10[,c(7,14,21)])
XtX11$mean_M_P=rowMeans(XtX11[,c(2,9,16)])
XtX11$mean_SD_P=rowMeans(XtX11[,c(3,10,17)])
XtX11$mean_M_XtX=rowMeans(XtX11[,c(4,11,18)])
XtX11$mean_SD_XtX=rowMeans(XtX11[,c(5,12,19)])
XtX11$mean_XtXst=rowMeans(XtX11[,c(6,13,20)])
XtX11$mean_log10.pval=rowMeans(XtX11[,c(7,14,21)])
XtX12$mean_M_P=rowMeans(XtX12[,c(2,9,16)])
```

```
XtX12$mean_SD_P=rowMeans(XtX12[,c(3,10,17)])
XtX12$mean_M_XtX=rowMeans(XtX12[,c(4,11,18)])
XtX12$mean_SD_XtX=rowMeans(XtX12[,c(5,12,19)])
XtX12$mean_XtXst=rowMeans(XtX12[,c(6,13,20)])
XtX12$mean_log10.pval=rowMeans(XtX12[,c(7,14,21)])
XtX13$mean_M_P=rowMeans(XtX13[,c(2,9,16)])
XtX13$mean_SD_P=rowMeans(XtX13[,c(3,10,17)])
XtX13$mean_M_XtX=rowMeans(XtX13[,c(4,11,18)])
XtX13$mean_SD_XtX=rowMeans(XtX13[,c(5,12,19)])
XtX13$mean_XtXst=rowMeans(XtX13[,c(6,13,20)])
XtX13$mean_log10.pval=rowMeans(XtX13[,c(7,14,21)])
XtX14$mean_M_P=rowMeans(XtX14[,c(2,9,16)])
XtX14$mean_SD_P=rowMeans(XtX14[,c(3,10,17)])
XtX14$mean_M_XtX=rowMeans(XtX14[,c(4,11,18)])
XtX14$mean_SD_XtX=rowMeans(XtX14[,c(5,12,19)])
XtX14$mean_XtXst=rowMeans(XtX14[,c(6,13,20)])
XtX14$mean_log10.pval=rowMeans(XtX14[,c(7,14,21)])
XtX15$mean_M_P=rowMeans(XtX15[,c(2,9,16)])
XtX15$mean_SD_P=rowMeans(XtX15[,c(3,10,17)])
XtX15$mean_M_XtX=rowMeans(XtX15[,c(4,11,18)])
XtX15$mean_SD_XtX=rowMeans(XtX15[,c(5,12,19)])
XtX15$mean_XtXst=rowMeans(XtX15[,c(6,13,20)])
XtX15$mean_log10.pval=rowMeans(XtX15[,c(7,14,21)])
XtX16$mean_M_P=rowMeans(XtX16[,c(2,9,16)])
XtX16$mean_SD_P=rowMeans(XtX16[,c(3,10,17)])
XtX16$mean_M_XtX=rowMeans(XtX16[,c(4,11,18)])
XtX16$mean_SD_XtX=rowMeans(XtX16[,c(5,12,19)])
XtX16$mean_XtXst=rowMeans(XtX16[,c(6,13,20)])
XtX16$mean_log10.pval=rowMeans(XtX16[,c(7,14,21)])
XtX17$mean_M_P=rowMeans(XtX17[,c(2,9,16)])
XtX17$mean_SD_P=rowMeans(XtX17[,c(3,10,17)])
XtX17$mean_M_XtX=rowMeans(XtX17[,c(4,11,18)])
```

```
XtX17$mean_SD_XtX=rowMeans(XtX17[,c(5,12,19)])
XtX17$mean_XtXst=rowMeans(XtX17[,c(6,13,20)])
XtX17$mean_log10.pval=rowMeans(XtX17[,c(7,14,21)])
XtX18$mean_M_P=rowMeans(XtX18[,c(2,9,16)])
XtX18$mean_SD_P=rowMeans(XtX18[,c(3,10,17)])
XtX18$mean_M_XtX=rowMeans(XtX18[,c(4,11,18)])
XtX18$mean_SD_XtX=rowMeans(XtX18[,c(5,12,19)])
XtX18$mean_XtXst=rowMeans(XtX18[,c(6,13,20)])
XtX18$mean_log10.pval=rowMeans(XtX18[,c(7,14,21)])
XtX19$mean_M_P=rowMeans(XtX19[,c(2,9,16)])
XtX19$mean_SD_P=rowMeans(XtX19[,c(3,10,17)])
XtX19$mean_M_XtX=rowMeans(XtX19[,c(4,11,18)])
XtX19$mean_SD_XtX=rowMeans(XtX19[,c(5,12,19)])
XtX19$mean_XtXst=rowMeans(XtX19[,c(6,13,20)])
XtX19$mean_log10.pval=rowMeans(XtX19[,c(7,14,21)])
XtX20$mean_M_P=rowMeans(XtX20[,c(2,9,16)])
XtX20$mean_SD_P=rowMeans(XtX20[,c(3,10,17)])
XtX20$mean_M_XtX=rowMeans(XtX20[,c(4,11,18)])
XtX20$mean_SD_XtX=rowMeans(XtX20[,c(5,12,19)])
XtX20$mean_XtXst=rowMeans(XtX20[,c(6,13,20)])
XtX20$mean_log10.pval=rowMeans(XtX20[,c(7,14,21)])

tXtX1=cbind(mrk1[,c(1:2)],XtX1[,c(22:27)])
tXtX2=cbind(mrk2[,c(1:2)],XtX2[,c(22:27)])
tXtX3=cbind(mrk3[,c(1:2)],XtX3[,c(22:27)])
tXtX4=cbind(mrk4[,c(1:2)],XtX4[,c(22:27)])
tXtX5=cbind(mrk5[,c(1:2)],XtX5[,c(22:27)])
tXtX6=cbind(mrk6[,c(1:2)],XtX6[,c(22:27)])
tXtX7=cbind(mrk7[,c(1:2)],XtX7[,c(22:27)])
tXtX8=cbind(mrk8[,c(1:2)],XtX8[,c(22:27)])
tXtX9=cbind(mrk9[,c(1:2)],XtX9[,c(22:27)])
tXtX10=cbind(mrk10[,c(1:2)],XtX10[,c(22:27)])
```

```
tXtX11=cbind(mrk11[,c(1:2)],XtX11[,c(22:27)])

tXtX12=cbind(mrk12[,c(1:2)],XtX12[,c(22:27)])

tXtX13=cbind(mrk13[,c(1:2)],XtX13[,c(22:27)])

tXtX14=cbind(mrk14[,c(1:2)],XtX14[,c(22:27)])

tXtX15=cbind(mrk15[,c(1:2)],XtX15[,c(22:27)])

tXtX16=cbind(mrk16[,c(1:2)],XtX16[,c(22:27)])

tXtX17=cbind(mrk17[,c(1:2)],XtX17[,c(22:27)])

tXtX18=cbind(mrk18[,c(1:2)],XtX18[,c(22:27)])

tXtX19=cbind(mrk19[,c(1:2)],XtX19[,c(22:27)])

tXtX20=cbind(mrk20[,c(1:2)],XtX20[,c(22:27)])


XtXtot<-
rbind(tXtX1,tXtX2,tXtX3,tXtX4,tXtX5,tXtX6,tXtX7,tXtX8,tXtX9,tXtX10,tXtX11,tXtX12,tXtX13,tX
tX14,tXtX15,tXtX16,tXtX17,tXtX18,tXtX19,tXtX20)

colnames(XtXtot)=c("chr", "pos", "mean_M_P", "mean_SD_P", "mean_M_XtX", "mean_SD_XtX",
"mean_XtXst", "mean_log10.pval")

#colnames(XtXtot)=c("chr", "pos", "M_P", "SD_P", "DELTA_P", "ACC_P", "M_XtX", "SD_XtX")

write.table(XtXtot, "./dataset16_AT_RH_XtX.txt", sep="\t")
```

**###Script to perform the local score appraoch on BFis, including the ranking for pvalue estimation (from Fabriello et al. 2017, Bonhomme et al. 2019)**

**# Rank the BFis data to provide a p-value**

```
Dataset5 <- read.table("/groups/holoe2plant/Loeiz/R_rpob/BLUPS_RH_AT.txt", h=T)

traits=colnames(Dataset5[,-1])


for(trait in traits) {

  data=read.table(paste("./dataset16_AT_RH_",trait,".txt",sep=""), h=T)


  rank=data[order(data$BFdB, decreasing=T),]

  rank$rank=c(1:nrow(rank))

  rank$pval=rank$rank/nrow(rank)

  write.table(rank, paste("./Res_pvalue",trait,".txt",sep=""), sep="\t", quote=F, row.names=FALSE)
```

}

#### Local score approach from Fariello et al 2017 & Bonhomme et al. 2019.

```
library(ggplot2)
library(data.table)
library(RColorBrewer)
library(data.table)
```

**#Import functions**

**# Function available on https://forge-dga.jouy.inra.fr/projects/local-score/**

```
source('/groups/holoe2plant/Loeiz/GEA/scorelocalfunctions.R')

sig_zone_local_score <- function(lindley,pos,th){
  zones=c(0,0,0,0)
  LIND=lindley
  auxpos=pos
  while(max(LIND,na.rm=TRUE)>=th){
    M_loc <- which.max(LIND)
    if(min(LIND[1:M_loc],na.rm=TRUE)==0){
      beg_peak <- max(which(LIND[1:M_loc]==min(LIND[1:M_loc],na.rm=TRUE)))+1 #found first 0 before max peak to identify peak start
    }else{
      beg_peak <- max(which(LIND[1:M_loc]==min(LIND[1:M_loc],na.rm=TRUE)))
    }
    if(length(which(LIND[M_loc+1:length(LIND)]==0))>1){
      end_peak <- min(which(LIND[M_loc+1:length(LIND)]==min(LIND[M_loc+1:length(LIND)],na.rm=TRUE)))+M_loc #found first 0 after max peak to identify peak end
    }else{
      end_peak <- M_loc
    }
    zones=rbind(zones, c(auxpos[beg_peak],auxpos[end_peak],auxpos[M_loc],max(LIND)))
```

```
    LIND <- LIND[-(beg_peak:end_peak)]
    auxpos <- auxpos[-(beg_peak:end_peak)]
  }
  zones=matrix(zones, ncol=4)
  zones=data.table(beg=zones[,1],end=zones[,2],max_zone=zones[,3],peak=zones[,4])
  if (nrow(zones)>1){zones=zones[-1,]}
  return(zones)
}

files = list.files("./output/", pattern=".assoc.txt", all.files=FALSE, full.names=FALSE)
files = tools::file_path_sans_ext(files)
files = tools::file_path_sans_ext(files)



MARF = 0.1
MISS = 17

for (trait in traits[1:length(traits)]){

  ### Choice of $\xi$ (1,2,3 or 4).
  xi=3

  data=read.table(paste("./Res_pvalue",trait,".txt",sep=""), h=T)
  colnames(data)[colnames(data)=="scaffold"]="chr"
  mydata=as.data.table(data[,c(which(colnames(data)=="chr"), which(colnames(data)=="pos"),
which(colnames(data)=="pval"), which(colnames(data)=="BFdB"))])
  mydata=mydata[order(mydata$pos, decreasing=F),]

    setkey(mydata, chr) # for sorting mydata according to chr, the column will be marked as sorted with
a key
    Nchr=length(mydata[,unique(chr)])
```

**### Computation of absolute position in the genome.**

```
#This is useful for doing genomewide plots.
chrInfo=mydata[,.(L=.N,cor=autocor(pval)),chr]
setkey(chrInfo,chr)
data.table(chr=mydata[,unique(chr),], S=cumsum(c(0,chrInfo$L[-Nchr])))
mydata$log_pval = -log10(mydata$pval)

mydata[,score:= mydata$log_pval-xi]
mean(mydata$score)
mydata[,lindley:=lindley(score),chr]

# The score mean must be negative, ksi must be chosen between mean and max of -log10(p-value)
mean(mydata$log_pval);max(mydata$log_pval)
mean(mydata$score)
#hist(mydata$score)
max(mydata$lindley)

# Compute significance threshold for each chromosome
## Uniform distribution of p-values
chrInfo[,th:=thresUnif(L, cor, xi),chr]
mydata=mydata[chrInfo]
sigZones <- mydata[chrInfo,sig_zone_local_score(lindley, pos, unique(th)),chr]
sigZones$QTL_length <- sigZones$end - sigZones$beg

#sigZones <- filter(sigZones,peak>0)
sigZones <- sigZones[which(sigZones$peak > 0),]

sigZonesSNPDens = data.frame()
all_SNP_signif_log4 = vector()

mydata$signif = round(mydata$lindley-mydata$th,3)
mydata$th = round(mydata$th,3)
```

```
  mydata$cor = round(mydata$cor,3)
  mydata$log_pval = round(mydata$log_pval,6)

  for (zone in c(1:nrow(sigZones))) {
    CHR = sigZones$chr[zone]
    BEG = sigZones$beg[zone]
    END = sigZones$end[zone]
    SNP_number_log3 = nrow(mydata[mydata$chr==CHR & mydata$signif>0 & mydata$log_pval>3])
    SNP_number_log4 = nrow(mydata[mydata$chr==CHR & mydata$signif>0 & mydata$log_pval>4])
    SNP_number_log5 = nrow(mydata[mydata$chr==CHR & mydata$signif>0 & mydata$log_pval>5])
    SNP_number_log6 = nrow(mydata[mydata$chr==CHR & mydata$signif>0 & mydata$log_pval>6])
    subsetzone = cbind(sigZones[zone,],SNP_number_log3,SNP_number_log4,SNP_number_log5,SNP_number_log6)
    sigZonesSNPDens = rbind(sigZonesSNPDens,subsetzone)
  results = as.data.frame(cbind(mydata$chr,mydata$pos,mydata$BFdB,mydata$log_pval,mydata$cor,mydata$lindley,mydata$th,mydata$signif))
  colnames(results) = c("chr","pos","BFdB", "log_pval","cor","lindley","th","signif")

  write.table(results,file=paste0("./Dataset17_",trait,".txt"),quote=F, sep="\t",row.names=F,)
  write.table(as.data.frame(sigZonesSNPDens),file=paste0("./Dataset18_",trait,"_ksi3.txt"),quote=F, sep="\t",row.names=F)

 }
```

##### Manhattan plots

```
for (trait in traits) {

 #emmax=read.table(paste("./Dataset18_",trait,"_ksi3.txt",sep=""), h=T)
 emmax=read.table(paste("./Dataset17_",trait,".txt",sep=""), h=T)
```

```
emmax$manhattan=0
emmax$col = 0

emmax[emmax$chr == 'A01',which(colnames(emmax)=="col")] = "grey20"
emmax[emmax$chr == 'A02',which(colnames(emmax)=="col")] = "grey40"
emmax[emmax$chr == 'A03',which(colnames(emmax)=="col")] = "grey20"
emmax[emmax$chr == 'A04',which(colnames(emmax)=="col")] = "grey40"
emmax[emmax$chr == 'A05',which(colnames(emmax)=="col")] = "grey20"
emmax[emmax$chr == 'A06',which(colnames(emmax)=="col")] = "grey40"
emmax[emmax$chr == 'A07',which(colnames(emmax)=="col")] = "grey20"
emmax[emmax$chr == 'A08',which(colnames(emmax)=="col")] = "grey40"
emmax[emmax$chr == 'A09',which(colnames(emmax)=="col")] = "grey20"
emmax[emmax$chr == 'A10',which(colnames(emmax)=="col")] = "grey40"

emmax[emmax$chr==unique(emmax$chr)[1],which(colnames(emmax)=="manhattan")]=
  emmax[emmax$chr==unique(emmax$chr)[1],which(colnames(emmax)=="pos")]
for (i in 2:length(unique(emmax$chr))){
  emmax[emmax$chr==unique(emmax$chr)[i],which(colnames(emmax)=="manhattan")]=
emmax[emmax$chr==unique(emmax$chr)[i],which(colnames(emmax)=="pos")]+max(emmax[emmax$chr==unique(emmax$chr)[i-1],which(colnames(emmax)=="manhattan")])
}

deb_emmax=min(emmax$manhattan)
fin_emmax=max(emmax$manhattan)
```

**### plot it**

```
tiff(paste("./Manhattan_local_score_", trait,"_xsi3.tiff",sep=""), res=600, width=25, height=15, unit="cm",compression="lzw")

m=matrix(1:5,5,1)
```

```
  layout(m,c(5),c(1.5,10,1.5,10,3))
  layout.show(5)

  par(mar=c(0,0,0,0), ps=8)
    plot(0,0,xlim=c(0,0.1),ylim=c(0,1),type="n",axes=FALSE,frame.plot=FALSE)
    text(0.05,0.2,paste(trait, sep=""),cex=1.5)
  par(mar=c(0,2,0.7,0.5),ps=8,mgp=c(0.6,0.3,0))

plot(emmax$manhattan,emmax$BFdB,xlab="",ylab="",frame.plot=FALSE,xlim=c(deb_emmax,fin_emmax),axes=FALSE, cex =0.3 ,  col=emmax$col ,pch=16)
    axis(1,cex.axis=1,lwd=0.5,xaxp=c(deb_emmax,fin_emmax,10),labels=F, tck=0)
    axis(2,cex.axis=1,lwd=0.5,tck=0.03)
    box(lwd=0.5)
    mtext("BFdB", side=2, line=1, cex=0.8)
  par(mar=c(0,0.7,0.7,0.5),ps=8,mgp=c(0.6,0.3,0))
    plot(0,0,xlim=c(0,1),ylim=c(0,1),type="n",axes=FALSE,frame.plot=FALSE)
  par(mar=c(0,2,0.7,0.5),ps=8,mgp=c(0.6,0.3,0))
    plot(emmax$manhattan, emmax$lindley, xlab="", ylab="", frame.plot=FALSE, xlim=c(deb_emmax, fin_emmax), axes=FALSE, cex=0.5, col=ifelse(
        (emmax$chr == "A06" & (emmax$pos >= 6655190 & emmax$pos <= 6659802))
        , "red", emmax$col), pch=16)
    axis(1,cex.axis=1,lwd=0.5,xaxp=c(deb_emmax,fin_emmax,10),labels=F, tck=0)
    axis(2,cex.axis=1,lwd=0.5,tck=0.03)
    abline(h=mean(emmax$th), col="grey", lty=2, las=1)
    box(lwd=0.5)
    mtext("lindley score", side=2, line=1, cex=0.8)
  par(mar=c(0,0,0,0), ps=8)
    plot(0,0,xlim=c(0,0.1),ylim=c(0,1),type="n",axes=FALSE,frame.plot=FALSE)

  dev.off()
}
```

**############ SP RH ###########**

**# Singular value decomposition (SVD) on matric of allele frequency**

```
om.mat=as.matrix(read.table("./Dataset13_SP_RH_sub1_1_mat_omega.out"))

listpop=as.data.frame(matrix(NA, ncol=2, nrow=26))

colnames(listpop)=c("pop", "color")

listpop$pop=c("ALTO-A","APIG-A","BISA-A","BOEL-A","BOLO-A","BRUZ-A","CAST-A","CAST-B","CAST-C","COLL-A","CORL-A","FOUG-A","GRAT-A","GRAT-B","ISNE-A","ISNE-B","LASC-A","MARI-A","MARI-B","MISI-A","MISI-B","MONR-A","MORD-A","PIAN-A","POLL-A","RENN-A")

listpop$color=c("orange", "deepskyblue4", "deepskyblue4", "deepskyblue4", "deepskyblue4",
    "deepskyblue4", "deepskyblue4", "deepskyblue4", "deepskyblue4", "deepskyblue4",
    "orange", "orange", "deepskyblue4", "deepskyblue4", "orange", "orange", "orange",
    "deepskyblue4", "deepskyblue4", "deepskyblue4", "deepskyblue4","orange",
    "deepskyblue4", "deepskyblue4", "deepskyblue4", "deepskyblue4")


# Singular value decomposition

om.svd=svd(om.mat)


# eigenvalues of the diagonal of the variance

om.eig=om.svd$d**2 # <=> om.eig=om.svd$d^2


perc.var=100*om.eig/sum(om.eig)


PC=cbind(listpop, om.svd$u[,1:2])

colnames(PC)=c("population", "color","PC1", "PC2")

write.table(PC, "./Dataset15.txt", sep="\t")
```

**### Concatenate Bayesian Factor (BFis in dB) and XtX index from different parallised anaylses on 20 sub-samples###**

**# Open files with identity of markers within each sub-samples (sub1 to sub19)**

```
for(i in 1:20) {

  assign( paste("mrk", i, sep=""),
```

```
        read.table(paste("/groups/holoe2plant/Loeiz/R_bis/Dataset10.snpdet.sub", i, sep=""), h=F) )
}
```

**### concatenation bayes factors files**

```
Dataset5 <- read.table("/groups/holoe2plant/Loeiz/R_rpob/BLUPS_RH_SP.txt", h=T)
traits=colnames(Dataset5[,-1])
```

**# first repeat**

```
for(i in 1:20) {
   assign( paste("sub", i, "_1", sep=""),
          read.table(paste("./Dataset13_SP_RH_sub",i,"_1_summary_betai_reg.out", sep=""), h=T) )
}
```

**# second repeat**

```
for(i in 1:20) {
   assign( paste("sub", i, "_2", sep=""),
          read.table(paste("./Dataset13_SP_RH_sub",i,"_2_summary_betai_reg.out", sep=""), h=T) )
}
```

**# Third repeat**

```
for(i in 1:20) {
   assign( paste("sub", i, "_3", sep=""),
          read.table(paste("./Dataset13_SP_RH_sub",i,"_3_summary_betai_reg.out", sep=""), h=T) )
}

for (i in 1:length(traits)) {

sub1=cbind(sub1_1[sub1_1$COVARIABLE==i,],sub1_2[sub1_2$COVARIABLE==i,],sub1_3[sub1_
3$COVARIABLE==i,])

sub2=cbind(sub2_1[sub2_1$COVARIABLE==i,],sub2_2[sub2_2$COVARIABLE==i,],sub2_3[sub2_
3$COVARIABLE==i,])

sub3=cbind(sub3_1[sub3_1$COVARIABLE==i,],sub3_2[sub3_2$COVARIABLE==i,],sub3_3[sub3_
3$COVARIABLE==i,])
```

```
sub4=cbind(sub4_1[sub4_1$COVARIABLE==i,],sub4_2[sub4_2$COVARIABLE==i,],sub4_3[sub4_3$COVARIABLE==i,])

sub5=cbind(sub5_1[sub5_1$COVARIABLE==i,],sub5_2[sub5_2$COVARIABLE==i,],sub5_3[sub5_3$COVARIABLE==i,])

sub6=cbind(sub6_1[sub6_1$COVARIABLE==i,],sub6_2[sub6_2$COVARIABLE==i,],sub6_3[sub6_3$COVARIABLE==i,])

sub7=cbind(sub7_1[sub7_1$COVARIABLE==i,],sub7_2[sub7_2$COVARIABLE==i,],sub7_3[sub7_3$COVARIABLE==i,])

sub8=cbind(sub8_1[sub8_1$COVARIABLE==i,],sub8_2[sub8_2$COVARIABLE==i,],sub8_3[sub8_3$COVARIABLE==i,])

sub9=cbind(sub9_1[sub9_1$COVARIABLE==i,],sub9_2[sub9_2$COVARIABLE==i,],sub9_3[sub9_3$COVARIABLE==i,])

sub10=cbind(sub10_1[sub10_1$COVARIABLE==i,],sub10_2[sub10_2$COVARIABLE==i,],sub10_3[sub10_3$COVARIABLE==i,])

sub11=cbind(sub11_1[sub11_1$COVARIABLE==i,],sub11_2[sub11_2$COVARIABLE==i,],sub11_3[sub11_3$COVARIABLE==i,])

sub12=cbind(sub12_1[sub12_1$COVARIABLE==i,],sub12_2[sub12_2$COVARIABLE==i,],sub12_3[sub12_3$COVARIABLE==i,])

sub13=cbind(sub13_1[sub13_1$COVARIABLE==i,],sub13_2[sub13_2$COVARIABLE==i,],sub13_3[sub13_3$COVARIABLE==i,])

sub14=cbind(sub14_1[sub14_1$COVARIABLE==i,],sub14_2[sub14_2$COVARIABLE==i,],sub14_3[sub14_3$COVARIABLE==i,])

sub15=cbind(sub15_1[sub15_1$COVARIABLE==i,],sub15_2[sub15_2$COVARIABLE==i,],sub15_3[sub15_3$COVARIABLE==i,])

sub16=cbind(sub16_1[sub16_1$COVARIABLE==i,],sub16_2[sub16_2$COVARIABLE==i,],sub16_3[sub16_3$COVARIABLE==i,])

sub17=cbind(sub17_1[sub17_1$COVARIABLE==i,],sub17_2[sub17_2$COVARIABLE==i,],sub17_3[sub17_3$COVARIABLE==i,])
```

```
sub18=cbind(sub18_1[sub18_1$COVARIABLE==i,],sub18_2[sub18_2$COVARIABLE==i,],sub18_3[sub18_3$COVARIABLE==i,])

sub19=cbind(sub19_1[sub19_1$COVARIABLE==i,],sub19_2[sub19_2$COVARIABLE==i,],sub19_3[sub19_3$COVARIABLE==i,])

sub20=cbind(sub20_1[sub20_1$COVARIABLE==i,],sub20_2[sub20_2$COVARIABLE==i,],sub20_3[sub20_3$COVARIABLE==i,])

sub1$mean_M_Pearson=rowMeans(sub1[,c(3,11,19)])
sub1$mean_BFdB=rowMeans(sub1[,c(5,13,21)])
sub1$mean_Beta_is=rowMeans(sub1[,c(6,14,22)])
sub2$mean_M_Pearson=rowMeans(sub2[,c(3,11,19)])
sub2$mean_BFdB=rowMeans(sub2[,c(5,13,21)])
sub2$mean_Beta_is=rowMeans(sub2[,c(6,14,22)])
sub3$mean_M_Pearson=rowMeans(sub3[,c(3,11,19)])
sub3$mean_BFdB=rowMeans(sub3[,c(5,13,21)])
sub3$mean_Beta_is=rowMeans(sub3[,c(6,14,22)])
sub4$mean_M_Pearson=rowMeans(sub4[,c(3,11,19)])
sub4$mean_BFdB=rowMeans(sub4[,c(5,13,21)])
sub4$mean_Beta_is=rowMeans(sub4[,c(6,14,22)])
sub5$mean_M_Pearson=rowMeans(sub5[,c(3,11,19)])
sub5$mean_BFdB=rowMeans(sub5[,c(5,13,21)])
sub5$mean_Beta_is=rowMeans(sub5[,c(6,14,22)])
sub6$mean_M_Pearson=rowMeans(sub6[,c(3,11,19)])
sub6$mean_BFdB=rowMeans(sub6[,c(5,13,21)])
sub6$mean_Beta_is=rowMeans(sub6[,c(6,14,22)])
sub7$mean_M_Pearson=rowMeans(sub7[,c(3,11,19)])
sub7$mean_BFdB=rowMeans(sub7[,c(5,13,21)])
sub7$mean_Beta_is=rowMeans(sub7[,c(6,14,22)])
sub8$mean_M_Pearson=rowMeans(sub8[,c(3,11,19)])
sub8$mean_BFdB=rowMeans(sub8[,c(5,13,21)])
sub8$mean_Beta_is=rowMeans(sub8[,c(6,14,22)])
```

```
sub9$mean_M_Pearson=rowMeans(sub9[,c(3,11,19)])
sub9$mean_BFdB=rowMeans(sub9[,c(5,13,21)])
sub9$mean_Beta_is=rowMeans(sub9[,c(6,14,22)])
sub10$mean_M_Pearson=rowMeans(sub10[,c(3,11,19)])
sub10$mean_BFdB=rowMeans(sub10[,c(5,13,21)])
sub10$mean_Beta_is=rowMeans(sub10[,c(6,14,22)])
sub11$mean_M_Pearson=rowMeans(sub11[,c(3,11,19)])
sub11$mean_BFdB=rowMeans(sub11[,c(5,13,21)])
sub11$mean_Beta_is=rowMeans(sub11[,c(6,14,22)])
sub12$mean_M_Pearson=rowMeans(sub12[,c(3,11,19)])
sub12$mean_BFdB=rowMeans(sub12[,c(5,13,21)])
sub12$mean_Beta_is=rowMeans(sub12[,c(6,14,22)])
sub13$mean_M_Pearson=rowMeans(sub13[,c(3,11,19)])
sub13$mean_BFdB=rowMeans(sub13[,c(5,13,21)])
sub13$mean_Beta_is=rowMeans(sub13[,c(6,14,22)])
sub14$mean_M_Pearson=rowMeans(sub14[,c(3,11,19)])
sub14$mean_BFdB=rowMeans(sub14[,c(5,13,21)])
sub14$mean_Beta_is=rowMeans(sub14[,c(6,14,22)])
sub15$mean_M_Pearson=rowMeans(sub15[,c(3,11,19)])
sub15$mean_BFdB=rowMeans(sub15[,c(5,13,21)])
sub15$mean_Beta_is=rowMeans(sub15[,c(6,14,22)])
sub16$mean_M_Pearson=rowMeans(sub16[,c(3,11,19)])
sub16$mean_BFdB=rowMeans(sub16[,c(5,13,21)])
sub16$mean_Beta_is=rowMeans(sub16[,c(6,14,22)])
sub17$mean_M_Pearson=rowMeans(sub17[,c(3,11,19)])
sub17$mean_BFdB=rowMeans(sub17[,c(5,13,21)])
sub17$mean_Beta_is=rowMeans(sub17[,c(6,14,22)])
sub18$mean_M_Pearson=rowMeans(sub18[,c(3,11,19)])
sub18$mean_BFdB=rowMeans(sub18[,c(5,13,21)])
sub18$mean_Beta_is=rowMeans(sub18[,c(6,14,22)])
sub19$mean_M_Pearson=rowMeans(sub19[,c(3,11,19)])
sub19$mean_BFdB=rowMeans(sub19[,c(5,13,21)])
```

```
sub19$mean_Beta_is=rowMeans(sub19[,c(6,14,22)])
sub20$mean_M_Pearson=rowMeans(sub20[,c(3,11,19)])
sub20$mean_BFdB=rowMeans(sub20[,c(5,13,21)])
sub20$mean_Beta_is=rowMeans(sub20[,c(6,14,22)])

tsub1=cbind(mrk1[,c(1:2)],sub1[,c(25,26,27)])
tsub2=cbind(mrk2[,c(1:2)],sub2[,c(25,26,27)])
tsub3=cbind(mrk3[,c(1:2)],sub3[,c(25,26,27)])
tsub4=cbind(mrk4[,c(1:2)],sub4[,c(25,26,27)])
tsub5=cbind(mrk5[,c(1:2)],sub5[,c(25,26,27)])
tsub6=cbind(mrk6[,c(1:2)],sub6[,c(25,26,27)])
tsub7=cbind(mrk7[,c(1:2)],sub7[,c(25,26,27)])
tsub8=cbind(mrk8[,c(1:2)],sub8[,c(25,26,27)])
tsub9=cbind(mrk9[,c(1:2)],sub9[,c(25,26,27)])
tsub10=cbind(mrk10[,c(1:2)],sub10[,c(25,26,27)])
tsub11=cbind(mrk11[,c(1:2)],sub11[,c(25,26,27)])
tsub12=cbind(mrk12[,c(1:2)],sub12[,c(25,26,27)])
tsub13=cbind(mrk13[,c(1:2)],sub13[,c(25,26,27)])
tsub14=cbind(mrk14[,c(1:2)],sub14[,c(25,26,27)])
tsub15=cbind(mrk15[,c(1:2)],sub15[,c(25,26,27)])
tsub16=cbind(mrk16[,c(1:2)],sub16[,c(25,26,27)])
tsub17=cbind(mrk17[,c(1:2)],sub17[,c(25,26,27)])
tsub18=cbind(mrk18[,c(1:2)],sub18[,c(25,26,27)])
tsub19=cbind(mrk19[,c(1:2)],sub19[,c(25,26,27)])
tsub20=cbind(mrk20[,c(1:2)],sub20[,c(25,26,27)])


datafinal=rbind(tsub1,tsub2,tsub3,tsub4,tsub5,tsub6,tsub7,tsub8,tsub9,tsub10,tsub11,tsub12,tsub13,tsub14,tsub15,tsub16,tsub17,tsub18,tsub19,tsub20)
colnames(datafinal)=c("scaffold","pos","M_Pearson","BFdB","Beta_is")

name=traits[i]
```

```
write.table(datafinal,paste("./dataset16_SP_RH_",name,".txt",sep=""),row.names=F,col.names=T,quote=F)
}
```

**### XtX values concatenation**

```
for(i in 1:20) {
  assign( paste("mrk", i, sep=""),
          read.table(paste("/groups/holoe2plant/Loeiz/R_bis/Dataset10.snpdet.sub", i, sep=""), h=F) )
}
```

**# first repeat**

```
for(i in 1:20) {
     assign( paste("XtX", i, "_1", sep=""),
             read.table(paste("./Dataset13_SP_RH_sub",i,"_1_summary_pi_xtx.out", sep=""), h=T) )
}
```

**# second repeat**

```
for(i in 1:20) {
     assign( paste("XtX", i, "_2", sep=""),
             read.table(paste("./Dataset13_SP_RH_sub",i,"_2_summary_pi_xtx.out", sep=""), h=T) )
}
```

**# Third repeat**

```
for(i in 1:20) {
     assign( paste("XtX", i, "_3", sep=""),
             read.table(paste("./Dataset13_SP_RH_sub",i,"_3_summary_pi_xtx.out", sep=""), h=T) )
}

XtX1=cbind(XtX1_1,XtX1_2,XtX1_3)
XtX2=cbind(XtX2_1,XtX2_2,XtX2_3)
XtX3=cbind(XtX3_1,XtX3_2,XtX3_3)
XtX4=cbind(XtX4_1,XtX4_2,XtX4_3)
```

```
XtX5=cbind(XtX5_1,XtX5_2,XtX5_3)
XtX6=cbind(XtX6_1,XtX6_2,XtX6_3)
XtX7=cbind(XtX7_1,XtX7_2,XtX7_3)
XtX8=cbind(XtX8_1,XtX8_2,XtX8_3)
XtX9=cbind(XtX9_1,XtX9_2,XtX9_3)
XtX10=cbind(XtX10_1,XtX10_2,XtX10_3)
XtX11=cbind(XtX11_1,XtX11_2,XtX11_3)
XtX12=cbind(XtX12_1,XtX12_2,XtX12_3)
XtX13=cbind(XtX13_1,XtX13_2,XtX13_3)
XtX14=cbind(XtX14_1,XtX14_2,XtX14_3)
XtX15=cbind(XtX15_1,XtX15_2,XtX15_3)
XtX16=cbind(XtX16_1,XtX16_2,XtX16_3)
XtX17=cbind(XtX17_1,XtX17_2,XtX17_3)
XtX18=cbind(XtX18_1,XtX18_2,XtX18_3)
XtX19=cbind(XtX19_1,XtX19_2,XtX19_3)
XtX20=cbind(XtX20_1,XtX20_2,XtX20_3)

XtX1$mean_M_P=rowMeans(XtX1[,c(2,9,16)])
XtX1$mean_SD_P=rowMeans(XtX1[,c(3,10,17)])
XtX1$mean_M_XtX=rowMeans(XtX1[,c(4,11,18)])
XtX1$mean_SD_XtX=rowMeans(XtX1[,c(5,12,19)])
XtX1$mean_XtXst=rowMeans(XtX1[,c(6,13,20)])
XtX1$mean_log10.pval=rowMeans(XtX1[,c(7,14,21)])
XtX2$mean_M_P=rowMeans(XtX2[,c(2,9,16)])
XtX2$mean_SD_P=rowMeans(XtX2[,c(3,10,17)])
XtX2$mean_M_XtX=rowMeans(XtX2[,c(4,11,18)])
XtX2$mean_SD_XtX=rowMeans(XtX2[,c(5,12,19)])
XtX2$mean_XtXst=rowMeans(XtX2[,c(6,13,20)])
XtX2$mean_log10.pval=rowMeans(XtX2[,c(7,14,21)])
XtX3$mean_M_P=rowMeans(XtX3[,c(2,9,16)])
XtX3$mean_SD_P=rowMeans(XtX3[,c(3,10,17)])
XtX3$mean_M_XtX=rowMeans(XtX3[,c(4,11,18)])
```

```
XtX3$mean_SD_XtX=rowMeans(XtX3[,c(5,12,19)])
XtX3$mean_XtXst=rowMeans(XtX3[,c(6,13,20)])
XtX3$mean_log10.pval=rowMeans(XtX3[,c(7,14,21)])
XtX4$mean_M_P=rowMeans(XtX4[,c(2,9,16)])
XtX4$mean_SD_P=rowMeans(XtX4[,c(3,10,17)])
XtX4$mean_M_XtX=rowMeans(XtX4[,c(4,11,18)])
XtX4$mean_SD_XtX=rowMeans(XtX4[,c(5,12,19)])
XtX4$mean_XtXst=rowMeans(XtX4[,c(6,13,20)])
XtX4$mean_log10.pval=rowMeans(XtX4[,c(7,14,21)])
XtX5$mean_M_P=rowMeans(XtX5[,c(2,9,16)])
XtX5$mean_SD_P=rowMeans(XtX5[,c(3,10,17)])
XtX5$mean_M_XtX=rowMeans(XtX5[,c(4,11,18)])
XtX5$mean_SD_XtX=rowMeans(XtX5[,c(5,12,19)])
XtX5$mean_XtXst=rowMeans(XtX5[,c(6,13,20)])
XtX5$mean_log10.pval=rowMeans(XtX5[,c(7,14,21)])
XtX6$mean_M_P=rowMeans(XtX6[,c(2,9,16)])
XtX6$mean_SD_P=rowMeans(XtX6[,c(3,10,17)])
XtX6$mean_M_XtX=rowMeans(XtX6[,c(4,11,18)])
XtX6$mean_SD_XtX=rowMeans(XtX6[,c(5,12,19)])
XtX6$mean_XtXst=rowMeans(XtX6[,c(6,13,20)])
XtX6$mean_log10.pval=rowMeans(XtX6[,c(7,14,21)])
XtX7$mean_M_P=rowMeans(XtX7[,c(2,9,16)])
XtX7$mean_SD_P=rowMeans(XtX7[,c(3,10,17)])
XtX7$mean_M_XtX=rowMeans(XtX7[,c(4,11,18)])
XtX7$mean_SD_XtX=rowMeans(XtX7[,c(5,12,19)])
XtX7$mean_XtXst=rowMeans(XtX7[,c(6,13,20)])
XtX7$mean_log10.pval=rowMeans(XtX7[,c(7,14,21)])
XtX8$mean_M_P=rowMeans(XtX8[,c(2,9,16)])
XtX8$mean_SD_P=rowMeans(XtX8[,c(3,10,17)])
XtX8$mean_M_XtX=rowMeans(XtX8[,c(4,11,18)])
XtX8$mean_SD_XtX=rowMeans(XtX8[,c(5,12,19)])
XtX8$mean_XtXst=rowMeans(XtX8[,c(6,13,20)])
```

```
XtX8$mean_log10.pval=rowMeans(XtX8[,c(7,14,21)])
XtX9$mean_M_P=rowMeans(XtX9[,c(2,9,16)])
XtX9$mean_SD_P=rowMeans(XtX9[,c(3,10,17)])
XtX9$mean_M_XtX=rowMeans(XtX9[,c(4,11,18)])
XtX9$mean_SD_XtX=rowMeans(XtX9[,c(5,12,19)])
XtX9$mean_XtXst=rowMeans(XtX9[,c(6,13,20)])
XtX9$mean_log10.pval=rowMeans(XtX9[,c(7,14,21)])
XtX10$mean_M_P=rowMeans(XtX10[,c(2,9,16)])
XtX10$mean_SD_P=rowMeans(XtX10[,c(3,10,17)])
XtX10$mean_M_XtX=rowMeans(XtX10[,c(4,11,18)])
XtX10$mean_SD_XtX=rowMeans(XtX10[,c(5,12,19)])
XtX10$mean_XtXst=rowMeans(XtX10[,c(6,13,20)])
XtX10$mean_log10.pval=rowMeans(XtX10[,c(7,14,21)])
XtX11$mean_M_P=rowMeans(XtX11[,c(2,9,16)])
XtX11$mean_SD_P=rowMeans(XtX11[,c(3,10,17)])
XtX11$mean_M_XtX=rowMeans(XtX11[,c(4,11,18)])
XtX11$mean_SD_XtX=rowMeans(XtX11[,c(5,12,19)])
XtX11$mean_XtXst=rowMeans(XtX11[,c(6,13,20)])
XtX11$mean_log10.pval=rowMeans(XtX11[,c(7,14,21)])
XtX12$mean_M_P=rowMeans(XtX12[,c(2,9,16)])
XtX12$mean_SD_P=rowMeans(XtX12[,c(3,10,17)])
XtX12$mean_M_XtX=rowMeans(XtX12[,c(4,11,18)])
XtX12$mean_SD_XtX=rowMeans(XtX12[,c(5,12,19)])
XtX12$mean_XtXst=rowMeans(XtX12[,c(6,13,20)])
XtX12$mean_log10.pval=rowMeans(XtX12[,c(7,14,21)])
XtX13$mean_M_P=rowMeans(XtX13[,c(2,9,16)])
XtX13$mean_SD_P=rowMeans(XtX13[,c(3,10,17)])
XtX13$mean_M_XtX=rowMeans(XtX13[,c(4,11,18)])
XtX13$mean_SD_XtX=rowMeans(XtX13[,c(5,12,19)])
XtX13$mean_XtXst=rowMeans(XtX13[,c(6,13,20)])
XtX13$mean_log10.pval=rowMeans(XtX13[,c(7,14,21)])
XtX14$mean_M_P=rowMeans(XtX14[,c(2,9,16)])
```

```
XtX14$mean_SD_P=rowMeans(XtX14[,c(3,10,17)])
XtX14$mean_M_XtX=rowMeans(XtX14[,c(4,11,18)])
XtX14$mean_SD_XtX=rowMeans(XtX14[,c(5,12,19)])
XtX14$mean_XtXst=rowMeans(XtX14[,c(6,13,20)])
XtX14$mean_log10.pval=rowMeans(XtX14[,c(7,14,21)])
XtX15$mean_M_P=rowMeans(XtX15[,c(2,9,16)])
XtX15$mean_SD_P=rowMeans(XtX15[,c(3,10,17)])
XtX15$mean_M_XtX=rowMeans(XtX15[,c(4,11,18)])
XtX15$mean_SD_XtX=rowMeans(XtX15[,c(5,12,19)])
XtX15$mean_XtXst=rowMeans(XtX15[,c(6,13,20)])
XtX15$mean_log10.pval=rowMeans(XtX15[,c(7,14,21)])
XtX16$mean_M_P=rowMeans(XtX16[,c(2,9,16)])
XtX16$mean_SD_P=rowMeans(XtX16[,c(3,10,17)])
XtX16$mean_M_XtX=rowMeans(XtX16[,c(4,11,18)])
XtX16$mean_SD_XtX=rowMeans(XtX16[,c(5,12,19)])
XtX16$mean_XtXst=rowMeans(XtX16[,c(6,13,20)])
XtX16$mean_log10.pval=rowMeans(XtX16[,c(7,14,21)])
XtX17$mean_M_P=rowMeans(XtX17[,c(2,9,16)])
XtX17$mean_SD_P=rowMeans(XtX17[,c(3,10,17)])
XtX17$mean_M_XtX=rowMeans(XtX17[,c(4,11,18)])
XtX17$mean_SD_XtX=rowMeans(XtX17[,c(5,12,19)])
XtX17$mean_XtXst=rowMeans(XtX17[,c(6,13,20)])
XtX17$mean_log10.pval=rowMeans(XtX17[,c(7,14,21)])
XtX18$mean_M_P=rowMeans(XtX18[,c(2,9,16)])
XtX18$mean_SD_P=rowMeans(XtX18[,c(3,10,17)])
XtX18$mean_M_XtX=rowMeans(XtX18[,c(4,11,18)])
XtX18$mean_SD_XtX=rowMeans(XtX18[,c(5,12,19)])
XtX18$mean_XtXst=rowMeans(XtX18[,c(6,13,20)])
XtX18$mean_log10.pval=rowMeans(XtX18[,c(7,14,21)])
XtX19$mean_M_P=rowMeans(XtX19[,c(2,9,16)])
XtX19$mean_SD_P=rowMeans(XtX19[,c(3,10,17)])
XtX19$mean_M_XtX=rowMeans(XtX19[,c(4,11,18)])
```

```
XtX19$mean_SD_XtX=rowMeans(XtX19[,c(5,12,19)])
XtX19$mean_XtXst=rowMeans(XtX19[,c(6,13,20)])
XtX19$mean_log10.pval=rowMeans(XtX19[,c(7,14,21)])
XtX20$mean_M_P=rowMeans(XtX20[,c(2,9,16)])
XtX20$mean_SD_P=rowMeans(XtX20[,c(3,10,17)])
XtX20$mean_M_XtX=rowMeans(XtX20[,c(4,11,18)])
XtX20$mean_SD_XtX=rowMeans(XtX20[,c(5,12,19)])
XtX20$mean_XtXst=rowMeans(XtX20[,c(6,13,20)])
XtX20$mean_log10.pval=rowMeans(XtX20[,c(7,14,21)])

tXtX1=cbind(mrk1[,c(1:2)],XtX1[,c(22:27)])
tXtX2=cbind(mrk2[,c(1:2)],XtX2[,c(22:27)])
tXtX3=cbind(mrk3[,c(1:2)],XtX3[,c(22:27)])
tXtX4=cbind(mrk4[,c(1:2)],XtX4[,c(22:27)])
tXtX5=cbind(mrk5[,c(1:2)],XtX5[,c(22:27)])
tXtX6=cbind(mrk6[,c(1:2)],XtX6[,c(22:27)])
tXtX7=cbind(mrk7[,c(1:2)],XtX7[,c(22:27)])
tXtX8=cbind(mrk8[,c(1:2)],XtX8[,c(22:27)])
tXtX9=cbind(mrk9[,c(1:2)],XtX9[,c(22:27)])
tXtX10=cbind(mrk10[,c(1:2)],XtX10[,c(22:27)])
tXtX11=cbind(mrk11[,c(1:2)],XtX11[,c(22:27)])
tXtX12=cbind(mrk12[,c(1:2)],XtX12[,c(22:27)])
tXtX13=cbind(mrk13[,c(1:2)],XtX13[,c(22:27)])
tXtX14=cbind(mrk14[,c(1:2)],XtX14[,c(22:27)])
tXtX15=cbind(mrk15[,c(1:2)],XtX15[,c(22:27)])
tXtX16=cbind(mrk16[,c(1:2)],XtX16[,c(22:27)])
tXtX17=cbind(mrk17[,c(1:2)],XtX17[,c(22:27)])
tXtX18=cbind(mrk18[,c(1:2)],XtX18[,c(22:27)])
tXtX19=cbind(mrk19[,c(1:2)],XtX19[,c(22:27)])
tXtX20=cbind(mrk20[,c(1:2)],XtX20[,c(22:27)])
```

```
XtXtot<-
rbind(tXtX1,tXtX2,tXtX3,tXtX4,tXtX5,tXtX6,tXtX7,tXtX8,tXtX9,tXtX10,tXtX11,tXtX12,tXtX13,tXtX14,tXtX15,tXtX16,tXtX17,tXtX18,tXtX19,tXtX20)

colnames(XtXtot)=c("chr", "pos", "mean_M_P", "mean_SD_P", "mean_M_XtX", "mean_SD_XtX", "mean_XtXst", "mean_log10.pval")

#colnames(XtXtot)=c("chr", "pos", "M_P", "SD_P", "DELTA_P", "ACC_P", "M_XtX", "SD_XtX")

write.table(XtXtot, "./dataset16_SP_RH_XtX.txt", sep="\t")
```

**###Script to perform the local score appraoch on BFis, including the ranking for pvalue estimation (from Fabriello et al. 2017, Bonhomme et al. 2019)**

**# Rank the BFis data to provide a p-value**

```
Dataset5 <- read.table("/groups/holoe2plant/Loeiz/R_rpob/BLUPS_RH_SP.txt", h=T)

traits=colnames(Dataset5[,-1])


for(trait in traits) {
  data=read.table(paste("./dataset16_SP_RH_",trait,".txt",sep=""), h=T)


  rank=data[order(data$BFdB, decreasing=T),]
  rank$rank=c(1:nrow(rank))
  rank$pval=rank$rank/nrow(rank)
  write.table(rank, paste("./Res_pvalue",trait,".txt",sep=""), sep="\t", quote=F, row.names=FALSE)
}
```

**## Local score approach from Fariello et al 2017 & Bonhomme et al. 2019.**

```
library(ggplot2)
library(data.table)
library(RColorBrewer)
library(data.table)
```

**#Import functions**

**# Function available on https://forge-dga.jouy.inra.fr/projects/local-score/**

```
source('/groups/holoe2plant/Loeiz/GEA/scorelocalfunctions.R')


sig_zone_local_score <- function(lindley,pos,th){
  zones=c(0,0,0,0)
  LIND=lindley
  auxpos=pos
  while(max(LIND,na.rm=TRUE)>=th){
    M_loc <- which.max(LIND)
    if(min(LIND[1:M_loc],na.rm=TRUE)==0){
      beg_peak <- max(which(LIND[1:M_loc]==min(LIND[1:M_loc],na.rm=TRUE)))+1 #found first 0 before max peak to identify peak start
    }else{
      beg_peak <- max(which(LIND[1:M_loc]==min(LIND[1:M_loc],na.rm=TRUE)))
    }
    if(length(which(LIND[M_loc+1:length(LIND)]==0))>1){
      end_peak <- min(which(LIND[M_loc+1:length(LIND)]==min(LIND[M_loc+1:length(LIND)],na.rm=TRUE)))+M_loc #found first 0 after max peak to identify peak end
    }else{
      end_peak <- M_loc
    }
    zones=rbind(zones, c(auxpos[beg_peak],auxpos[end_peak],auxpos[M_loc],max(LIND)))
    LIND <- LIND[-(beg_peak:end_peak)]
    auxpos <- auxpos[-(beg_peak:end_peak)]
  }
  zones=matrix(zones, ncol=4)
  zones=data.table(beg=zones[,1],end=zones[,2],max_zone=zones[,3],peak=zones[,4])
  if (nrow(zones)>1){zones=zones[-1,]}
  return(zones)
}


files = list.files("./output/", pattern=".assoc.txt", all.files=FALSE, full.names=FALSE)
files = tools::file_path_sans_ext(files)
```

```
files = tools::file_path_sans_ext(files)



MARF = 0.1
MISS = 17

for (trait in traits[1:length(traits)]){

  ### Choice of $\xi$ (1,2,3 or 4).
  xi=3

  data=read.table(paste("./Res_pvalue",trait,".txt",sep=""), h=T)
  colnames(data)[colnames(data)=="scaffold"]="chr"
  mydata=as.data.table(data[,c(which(colnames(data)=="chr"), which(colnames(data)=="pos"), which(colnames(data)=="pval"), which(colnames(data)=="BFdB"))])
  mydata=mydata[order(mydata$pos, decreasing=F),]

    setkey(mydata, chr) # for sorting mydata according to chr, the column will be marked as sorted with a key
    Nchr=length(mydata[,unique(chr)])

  ### Computation of absolute position in the genome.
    #This is useful for doing genomewide plots.
    chrInfo=mydata[,.(L=.N,cor=autocor(pval)),chr]
    setkey(chrInfo,chr)
    data.table(chr=mydata[,unique(chr),], S=cumsum(c(0,chrInfo$L[-Nchr])))
    mydata$log_pval = -log10(mydata$pval)

    mydata[,score:= mydata$log_pval-xi]
    mean(mydata$score)
    mydata[,lindley:=lindley(score),chr]
```

**# The score mean must be negative, ksi must be chosen between mean and max of -log10(p-value)**

```
mean(mydata$log_pval);max(mydata$log_pval)
mean(mydata$score)
#hist(mydata$score)
max(mydata$lindley)

# Compute significance threshold for each chromosome
## Uniform distribution of p-values
chrInfo[,th:=thresUnif(L, cor, xi),chr]
mydata=mydata[chrInfo]
sigZones <- mydata[chrInfo,sig_zone_local_score(lindley, pos, unique(th)),chr]
sigZones$QTL_length <- sigZones$end - sigZones$beg

#sigZones <- filter(sigZones,peak>0)
sigZones <- sigZones[which(sigZones$peak > 0),]

sigZonesSNPDens = data.frame()
all_SNP_signif_log4 = vector()

mydata$signif = round(mydata$lindley-mydata$th,3)
mydata$th = round(mydata$th,3)
mydata$cor = round(mydata$cor,3)
mydata$log_pval = round(mydata$log_pval,6)

for (zone in c(1:nrow(sigZones))) {
  CHR = sigZones$chr[zone]
  BEG = sigZones$beg[zone]
  END = sigZones$end[zone]
  SNP_number_log3 = nrow(mydata[mydata$chr==CHR & mydata$signif>0 & mydata$log_pval>3])
  SNP_number_log4 = nrow(mydata[mydata$chr==CHR & mydata$signif>0 & mydata$log_pval>4])
```

SNP_number_log5 = nrow(mydata[mydata$chr==CHR & mydata$signif>0 & mydata$log_pval>5])

SNP_number_log6 = nrow(mydata[mydata$chr==CHR & mydata$signif>0 & mydata$log_pval>6])

subsetzone = cbind(sigZones[zone,],SNP_number_log3 ,SNP_number_log4,SNP_number_log5,SNP_number_log6)

sigZonesSNPDens = rbind(sigZonesSNPDens,subsetzone)

}

results = as.data.frame(cbind(mydata$chr,mydata$pos,mydata$BFdB,mydata$log_pval,mydata$cor,mydata$lindley,mydata$th,mydata$signif))

colnames(results) = c("chr","pos","BFdB", "log_pval","cor","lindley","th","signif")

write.table(results,file=paste0("./Dataset17_",trait,".txt"),quote=F, sep="\t",row.names=F,)

write.table(as.data.frame(sigZonesSNPDens),file=paste0("./Dataset18_",trait,"_ksi3.txt"),quote=F, sep="\t",row.names=F)

}

**### Manhattan plots**

for (trait in traits) {

#emmax=read.table(paste("./Dataset18_",trait,"_ksi3.txt",sep=""), h=T)

emmax=read.table(paste("./Dataset17_",trait,".txt",sep=""), h=T)

emmax$manhattan=0

emmax$col = 0

emmax[emmax$chr == 'A01',which(colnames(emmax)=="col")] = "grey20"

emmax[emmax$chr == 'A02',which(colnames(emmax)=="col")] = "grey40"

emmax[emmax$chr == 'A03',which(colnames(emmax)=="col")] = "grey20"

emmax[emmax$chr == 'A04',which(colnames(emmax)=="col")] = "grey40"

```
emmax[emmax$chr == 'A05',which(colnames(emmax)=="col")] = "grey20"
emmax[emmax$chr == 'A06',which(colnames(emmax)=="col")] = "grey40"
emmax[emmax$chr == 'A07',which(colnames(emmax)=="col")] = "grey20"
emmax[emmax$chr == 'A08',which(colnames(emmax)=="col")] = "grey40"
emmax[emmax$chr == 'A09',which(colnames(emmax)=="col")] = "grey20"
emmax[emmax$chr == 'A10',which(colnames(emmax)=="col")] = "grey40"

emmax[emmax$chr==unique(emmax$chr)[1],which(colnames(emmax)=="manhattan")]=
  emmax[emmax$chr==unique(emmax$chr)[1],which(colnames(emmax)=="pos")]
for (i in 2:length(unique(emmax$chr))){
  emmax[emmax$chr==unique(emmax$chr)[i],which(colnames(emmax)=="manhattan")]=

emmax[emmax$chr==unique(emmax$chr)[i],which(colnames(emmax)=="pos")]+max(emmax[emma
x$chr==unique(emmax$chr)[i-1],which(colnames(emmax)=="manhattan")])
}

deb_emmax=min(emmax$manhattan)
fin_emmax=max(emmax$manhattan)

### plot it
tiff(paste("./Manhattan_local_score_", trait,"_xsi3.tiff",sep=""), res=600, width=25, height=15,
unit="cm",compression="lzw")

m=matrix(1:5,5,1)
layout(m,c(5),c(1.5,10,1.5,10,3))
layout.show(5)

par(mar=c(0,0,0,0), ps=8)
  plot(0,0,xlim=c(0,0.1),ylim=c(0,1),type="n",axes=FALSE,frame.plot=FALSE)
  text(0.05,0.2,paste(trait, sep=""),cex=1.5)
par(mar=c(0,2,0.7,0.5),ps=8,mgp=c(0.6,0.3,0))

plot(emmax$manhattan,emmax$BFdB,xlab="",ylab="",frame.plot=FALSE,xlim=c(deb_emmax,fin_e
mmax),axes=FALSE, cex =0.3 ,  col=emmax$col ,pch=16)
```

```
    axis(1,cex.axis=1,lwd=0.5,xaxp=c(deb_emmax,fin_emmax,10),labels=F, tck=0)
    axis(2,cex.axis=1,lwd=0.5,tck=0.03)
    box(lwd=0.5)
    mtext("BFdB", side=2, line=1, cex=0.8)
  par(mar=c(0,0.7,0.7,0.5),ps=8,mgp=c(0.6,0.3,0))
    plot(0,0,xlim=c(0,1),ylim=c(0,1),type="n",axes=FALSE,frame.plot=FALSE)
  par(mar=c(0,2,0.7,0.5),ps=8,mgp=c(0.6,0.3,0))
    plot(emmax$manhattan, emmax$lindley, xlab="", ylab="", frame.plot=FALSE, xlim=c(deb_emmax, fin_emmax), axes=FALSE, cex=0.5,
        col=ifelse(
        (emmax$chr == "A07" & (emmax$pos >= 24921794 & emmax$pos <= 24924137))
        , "red", emmax$col), pch=16)
    axis(1,cex.axis=1,lwd=0.5,xaxp=c(deb_emmax,fin_emmax,10),labels=F, tck=0)
    axis(2,cex.axis=1,lwd=0.5,tck=0.03)
    abline(h=mean(emmax$th), col="grey", lty=2, las=1)
    box(lwd=0.5)
    mtext("lindley score", side=2, line=1, cex=0.8)
  par(mar=c(0,0,0,0), ps=8)
    plot(0,0,xlim=c(0,0.1),ylim=c(0,1),type="n",axes=FALSE,frame.plot=FALSE)

  dev.off()
}
```

**###### Script to compute enrichment for genetic differentiation index XtX #####**

**### AT RH ###**

```
XtX=read.table("./dataset16_AT_RH_XtX.txt",h=T,na.strings=".")
chrom=as.data.frame(matrix(NA, ncol=2, nrow=10))
colnames(chrom)=c("scaffold", "chrom")
chrom$scaffold=unique(XtX$chr)
chrom$chrom=c(1:10)
XtX$chromo=chrom[match(XtX$chr, chrom$scaffold),which(colnames(chrom)=="chrom")]
XtX$SNP_ID=paste(XtX$chromo,XtX$pos,sep="_")
```

```
#i = nb of top XtX
N=round(0.0005*nrow(XtX),0)

XtX$rank=rank(-(XtX$mean_M_XtX), na.last =T, ties.method = "average")
top=XtX[XtX$rank<=N,]

# read results file
data=read.table("/groups/holoe2plant/Loeiz/R_rpob/BLUPS_RH_AT.txt", h=T)
RE=as.data.frame(matrix(NA, ncol=4, nrow=ncol(data[-1])))
colnames(RE)= c("Traits", "ntops", "Enrichment", "pvalue")
RE$Traits=unique(colnames(data[-1]))

terms=as.factor(RE$Traits)

#define threshold top XtX
threshold=round(0.0005*nrow(XtX),0)
# define number of permutation
perm=10000

for (j in terms){
  data=read.table(paste("./Dataset17_",j,".txt",sep=""), h=T, na.strings=".")
  data$chromo=chrom[match(data$chr, chrom$scaffold),which(colnames(chrom)=="chrom")]
  data$SNP_ID=paste(data$chromo,data$pos,sep="_")
  data$rank=rank(-(data$lindley), na.last =T, ties.method = "average")
  data1=data[data$rank<=threshold,]

  Z=append(top$SNP_ID,data1$SNP_ID)
  U=length(unique(Z))
  E=((length(Z)-U)/nrow(top))/(nrow(data1)/nrow(data))

  rand=c()##set up the vector to accumulate the results from permutations
```

```
pb = txtProgressBar(min = 0, max = perm, style = 3)

  for (chr in 1:10){
  assign(paste("pv",chr,sep=""),data$rank[data[,which(colnames(data)=="chromo")]==chr])
}
for(i in 1:perm){
  X=sample(1:10,10)
  Y=sample(c(T,F),10,replace=T)
  for (n in 1:length(X)){
    a=get(paste("pv",X[n],sep=""))
    if(Y[n]==T){a=rev(a)}
    assign(paste("r",n,sep=""),a)
  }
  rrank=c(r1, r2, r3, r4, r5, r6, r7, r8, r9, r10)

  W=sample(1:length(rrank),1)
  rrr=rrank[c(W:length(rrank), 1:(W-1))]

  data1rand=data[rrr<=nrow(data1),]

  Zrand=append(top$SNP_ID,data1rand$SNP_ID)
  Urand=length(unique(Zrand))

  Erand=((length(Zrand)-Urand)/nrow(top))/(nrow(data1)/nrow(data))
  rand=c(rand, Erand)
  # update progress bar
  setTxtProgressBar(pb, i)
}
close(pb)

out=list()
out$ntop=length(Z)-U
```

```
  out$E=E
  out$pvalue=length(which(rand > E))/perm

  RE[match(j,RE$Traits),c(2,3,4)]=c(out$ntop,out$E,out$pvalue)
}

write.table(RE,paste("./Dataset19_AT_RH.txt",sep=""),sep="\t",row.names=F,col.names=T,quote=F)
```

**########### Top SNP ##########**

```
XtX=read.table("./dataset16_AT_RH_XtX.txt",h=T,na.strings=".")
top_XtX = head(XtX[order(XtX$mean_M_XtX, decreasing=TRUE),],round(0.0005*nrow(XtX),0))

write.table(top_XtX,file="Top_XtX_rpoB_AT_RH",quote=F, sep="\t",row.names=F,)


XtX=read.table("./dataset16_SP_RH_XtX.txt",h=T,na.strings=".")
top_XtX = head(XtX[order(XtX$mean_M_XtX, decreasing=TRUE),],round(0.0005*nrow(XtX),0))

write.table(top_XtX,file="Top_XtX_rpoB_SP_RH",quote=F, sep="\t",row.names=F,)
```

**The following scripts have been used for genome mapping and allele frequency matrix estimation**

```
include: 'config.py'

sample=[]
for x in config['SAMPLES']:
    tempo = x.split('/')[-1]
    if tempo.split('_')[0] not in sample:
        sample.append(tempo.split('_')[0])
```

```
sample_dir = config['SAMPLE_DIR']
genome = config['GENOME'].split('/')[-1]
genome_dir = config['GENOME_DIR']
MAPQ = config['QUALITY']
specie = config['SPECIE']
centro = config['CENTRO']
dico = '.'.join(genome.split(".")[:-1])

rule all:
    input:
        expand("reports/{specie}/{sample}.fastp.html", sample=sample, specie=specie),
        expand("reports/{specie}/{sample}.fastp.json", sample=sample, specie=specie),
        expand("wgscoverageplotter/"+str(specie)+"/{genome}/{sample}_coverage_plot.svg", sample=sample, genome=genome),
        expand("mapping/stats/{specie}/{genome}/{sample}/{sample}.bam.stats", sample=sample, genome=genome, specie=specie),
        expand("mapping/stats/{specie}/{genome}/{sample}/{sample}_dedup.bam.stats", sample=sample, genome=genome, specie=specie),
expand("VCF/varscan/{specie}/{genome}/pool_varscan_{specie}_{genome}_baSNP_contig_MAF_filtered_scaff_annotated.vcf", genome=genome, specie=specie),
        expand("plots/{specie}/{genome}/ADP.png", genome=genome, specie=specie),
        expand("plots/{specie}/{genome}/ADP.png", genome=genome, specie=specie),
        expand("plots/{specie}/{genome}/ADP_stats.txt", genome=genome, specie=specie),
        expand("plots/{specie}/{genome}/SD_stats.txt", genome=genome, specie=specie),
        expand("centro_number_SNPs_{specie}_{genome}.txt", genome=genome, specie=specie),
expand("VCF/varscan/{specie}/{genome}/pool_varscan_{specie}_{genome}_baSNP_contig_MAF.vcf", genome=genome, specie=specie),
        expand("cleaned_reads/{specie}/{sample}.clean_R1.fq.gz", sample=sample, specie=specie),
        expand("cleaned_reads/{specie}/{sample}.clean_R2.fq.gz", sample=sample, specie=specie),
        expand("index/{genome}.bwt", genome=genome),
        expand("index/{genome}.fai", genome=genome),
```

```
        expand("index/"+str(genome)+"_contig.txt", genome=genome),
        expand("index/"+str(dico)+".dict", genome=genome),
        expand("mapping/bam/{specie}/{genome}/{sample}/{sample}.bam", sample=sample, genome=genome, specie=specie),
        expand("mapping/bam/{specie}/{genome}/{sample}/{sample}_dedup.bam", sample=sample, genome=genome, specie=specie),
        expand("mapping/bam/{specie}/{genome}/{sample}/{sample}_dedup.bam.bai", sample=sample, genome=genome, specie=specie),
        expand("VCF/varscan/{specie}/{genome}/pool_varscan_{specie}_{genome}.vcf", genome=genome, specie=specie),
        expand("VCF/varscan/{specie}/{genome}/pool_varscan_{specie}_{genome}_baSNP.vcf", genome=genome, specie=specie),
expand("VCF/varscan/{specie}/{genome}/pool_varscan_{specie}_{genome}_baSNP_contig.vcf", genome=genome, specie=specie),
expand("VCF/varscan/{specie}/{genome}/pool_varscan_{specie}_{genome}_baSNP_contig_MAF_filtered.recode.vcf", genome=genome, specie=specie),
expand("VCF/varscan/{specie}/{genome}/pool_varscan_{specie}_{genome}_baSNP_contig_MAF_filtered_scaff.recode.vcf", genome=genome, specie=specie)

rule fastp:
    input:
        read1=str(sample_dir)+"{sample}_R1.fastq.gz",
        read2=str(sample_dir)+"{sample}_R2.fastq.gz"
    output:
        read1=temp("cleaned_reads/{specie}/{sample}.clean_R1.fq.gz"),
        read2=temp("cleaned_reads/{specie}/{sample}.clean_R2.fq.gz"),
        report_html="reports/{specie}/{sample}.fastp.html",
        report_json="reports/{specie}/{sample}.fastp.json"
    params:
        threads="5",
        memory = "3G",
        job_name="fastp"
```

```
    log:
        err="logs/fastp_sample/{specie}/{sample}.log"
    conda:
        "envs/fastp"
    shell:
        "fastp -i {input.read1} -I {input.read2} -o {output.read1} -O {output.read2} -w {params.threads} -h {output.report_html} -j {output.report_json} 2> {log.err}"

rule index:
    input:
        str(genome_dir)+str(genome)
    output:
        genome="index/"+str(genome),
        fai="index/"+str(genome)+".fai",
        index="index/"+str(genome)+".bwt",
    params:
        index="./index/"+str(genome),
        threads="5",
        memory = "3G",
        job_name="index"
    log:
        err="logs/index/"+str(genome)+".log"
    shell:
        "mkdir -p index/;"
        "cp {input} ./index/;"
        "samtools faidx {params.index} -o {output.genome} 2> {log.err};"
        "bwa index -p {params.index} {output.genome} 2> {log.err};"

rule picard_dict:
    input:
        "index/" + str(genome)
    output:
```

```
        "index/"+str(dico)+".dict"
    params:
        threads="5",
        memory = "3G",
        job_name="picard_dict"
    log:
        err="logs/picard_dict/" + str(dico) + ".log"
    shell:
        "picard CreateSequenceDictionary -R {input} -O {output} 2> {log.err};"

rule map:
    input:
        genome="index/{genome}",
        read1="cleaned_reads/{specie}/{sample}.clean_R1.fq.gz",
        read2="cleaned_reads/{specie}/{sample}.clean_R2.fq.gz"
    output:
        temp("mapping/bam/{specie}/{genome}/{sample}/{sample}.bam")
    params:
        qual=MAPQ,
        threads="30",
        memory="2G",
        job_name="map",
        RG = '\"@RG\\tID:{sample}\\tPL:ILLUMINA\\tLB:LIB-{sample}\\tSM:{sample}\"'
    log:
        err="logs/map/{specie}/{genome}/{sample}.log"
    shell:
        "set +u;"
        "bwa mem -M -R {params.RG} -t {params.threads}  {input.genome} {input.read1} {input.read2} | samtools sort -T {wildcards.sample} -@ {params.threads} -O bam -o {output} 2> {log.err};"
        "set -u;"

rule fixmate_sort_n:
```

```
    input:
        "mapping/bam/{specie}/{genome}/{sample}/{sample}.bam"
    output:
        temp("mapping/bam/{specie}/{genome}/{sample}/{sample}_nsorted.bam")
    params:
        threads="5",
        memory="3G",
        job_name="fixmate_sort_n",
        dir="mapping/bam/{specie}/{genome}/{sample}/"
    log:
        err="logs/fixmate_sort/{specie}/{genome}/{sample}.log"
    shell:
        "samtools sort -n -T {wildcards.sample} -@ {params.threads} {input} -O BAM -o {params.dir}{wildcards.sample}_nsorted.bam 2> {log.err}"

rule fixmate:
    input:
        "mapping/bam/{specie}/{genome}/{sample}/{sample}_nsorted.bam"
    output:
        temp("mapping/bam/{specie}/{genome}/{sample}/{sample}_fm_temp.bam")
    params:
        threads="12",
        memory="4G",
        job_name="fixmate",
        dir="mapping/bam/{specie}/{genome}/{sample}/"
    log:
        err="logs/fixmate/{specie}/{genome}/{sample}.log"
    shell:
        "samtools fixmate -@ {params.threads} -rpcm {input} {output} 2> {log.err}"

rule fixmate_sort_coor:
    input:
```

```
        "mapping/bam/{specie}/{genome}/{sample}/{sample}_fm_temp.bam"
    output:
        temp("mapping/bam/{specie}/{genome}/{sample}/{sample}_fm.bam")
    params:
        threads="5",
        memory="3G",
        job_name="fixmate_sort_coor",
        dir="mapping/bam/{specie}/{genome}/{sample}/"
    log:
        err="logs/fixmate_sort/{specie}/{genome}/{sample}.log"
    shell:
        "samtools sort -T {wildcards.sample} -@ {params.threads} {input} -O BAM -o {params.dir}{wildcards.sample}_fm.bam 2> {log.err}"


rule markdup:
    input:
        "mapping/bam/{specie}/{genome}/{sample}/{sample}_fm.bam"
    output:
        temp("mapping/bam/{specie}/{genome}/{sample}/{sample}_dedup.bam")
    params:
        metrics="mapping/bam/{specie}/{genome}/{sample}/{sample}_MD_metrics.txt",
        threads="12",
        memory="4G",
        job_name="MarkDuplicates"
    log:
        err="logs/markdup/{specie}/{genome}/{sample}.log"
    shell:
        "samtools markdup -@ {params.threads} -r {input} {output} 2> {log.err};"


rule samtools_index:
    input:
        "mapping/bam/{specie}/{genome}/{sample}/{sample}_dedup.bam"
```

```
    output:
        temp("mapping/bam/{specie}/{genome}/{sample}/{sample}_dedup.bam.bai")
    params:
        threads="12",
        memory="4G",
        job_name="sam_index"
    log:
        err="logs/samtools_index/{specie}/{genome}/{sample}.log"
    shell:
        "samtools index {input} > {output} 2> {log.err};"


rule wgscoverageplotter:
    input:
        bam="mapping/bam/{specie}/{genome}/{sample}/{sample}_dedup.bam",
        bai="mapping/bam/{specie}/{genome}/{sample}/{sample}_dedup.bam.bai",
        index="index/"+str(genome),
        picard_index="index/"+str(dico)+".dict"
    output:
        "wgscoverageplotter/{specie}/{genome}/{sample}_coverage_plot.svg"
    params:
        sample="{sample}",
        threads="12",
        memory="4G",
        job_name="wgscoverageplotter"
    log:
        err="logs/wgscoverageplotter/{specie}/{genome}/{sample}.log"
    shell:
        'wgscoverageplotter.py --dimension 1500x500 -C -1 --clip -R {input.index} {input.bam} -I "..." --
percentile median > {output}'


rule stats:
    input:
```

```
        bam="mapping/bam/{specie}/{genome}/{sample}/{sample}.bam",
        markdup="mapping/bam/{specie}/{genome}/{sample}/{sample}_dedup.bam"
    output:
        bam="mapping/stats/{specie}/{genome}/{sample}/{sample}.bam.stats",
        markdup="mapping/stats/{specie}/{genome}/{sample}/{sample}_dedup.bam.stats"
    params:
        threads="5",
        memory="3G",
        job_name="stats"
    log:
        err="logs/stats/{specie}/{genome}/{sample}.log"
    shell:
        "samtools stats -@ {params.threads} {input.bam} > {output.bam} 2> {log.err};"
        "samtools stats -@ {params.threads} {input.markdup} > {output.markdup} 2> {log.err};"


rule sample_list:
    output:
        "sample_list.txt"
    params:
        threads="1",
        memory="1G",
        job_name="sample_list"
    log:
        err="logs/sample_list/sample_list.log"
    run:
        fichier = open("sample_list.txt", "w+")
        for x in sample:
            fichier.write(str(x) + "\n")

rule mpileup:
    input:
```

```
        sample_list="sample_list.txt",
        fasta="index/{genome}",
        bam=expand("mapping/bam/{specie}/{genome}/{sample}/{sample}_dedup.bam", sample=sample, genome=genome, specie=specie)
    output:
        temp("VCF/varscan/{specie}/{genome}/pool_varscan_{specie}_{genome}.vcf")
    params:
        threads="30",
        memory="2G",
        job_name="mpileup"
    log:
        err="logs/mpileup/{specie}/{genome}.log"
    shell:
        """
        set +e
        samtools mpileup -q 30 -B -A -d 1000 -f {input.fasta} {input.bam} | varscan mpileup2cns --output-vcf 1 --min-avg-qual 18 --vcf-sample-list {input.sample_list} > {output} 2> {log.err};
        exitcode=$?
        if [ $exitcode -eq 1 ]
        then
            exit 1
        else
            exit 0
        fi
        """


rule bcftools:
    input:
        "VCF/varscan/{specie}/{genome}/pool_varscan_{specie}_{genome}.vcf"
    output:
        temp("VCF/varscan/{specie}/{genome}/pool_varscan_{specie}_{genome}_baSNP.vcf")
    params:
```

```
        threads="12",
        memory="2G",
        job_name="bcftools"
    log:
        err="logs/bcftools/{specie}/{genome}.log"
    shell:
        "bcftools view {input} --types snps -m 2 -M 2 -o {output}"

rule awk_get_contig:
    input:
        "index/"+str(genome)+".fai"
    output:
        "index/"+str(genome)+"_contig.txt"
    params:
        threads="12",
        memory="2G",
        job_name="awk_get_contiig"
    log:
        "logs/awk_contig/"+str(genome)+".log"
    shell:
        """
        awk '{{printf("##contig=<ID=%s,length=%d>\\n",$1,$2);}}' {input} > {output}
        """

rule awk_insert_contig:
    input:
        contig="index/"+str(genome)+"_contig.txt",
        vcf="VCF/varscan/{specie}/{genome}/pool_varscan_{specie}_{genome}_baSNP.vcf"
    output:
        temp("VCF/varscan/{specie}/{genome}/pool_varscan_{specie}_{genome}_baSNP_contig.vcf")
    params:
        threads="12",
```

```
        memory="2G",
        job_name="awk_insert_contiig"
    log:
        "logs/awk_insert_contig/{specie}/{genome}/.log"
    run:
        contig = []
        with open(input.contig, "r") as f:
            for x in f:
                contig.append(x)
            fichier = open(str(output), "w+")
            with open(input.vcf, "r") as g:
                for x in g:
                    if x[:6] == '#CHROM':
                        for y in contig:
                            x = y + x
                    fichier.write(str(x))
        #shell("awk '/^#CHROM/ {{ printf(contig);}} {{print;}}' {input.vcf} > {output}")

rule bcftools_MAF:
    input:
        "VCF/varscan/{specie}/{genome}/pool_varscan_{specie}_{genome}_baSNP_contig.vcf"
    output:
        "VCF/varscan/{specie}/{genome}/pool_varscan_{specie}_{genome}_baSNP_contig_MAF.vcf"
    params:
        threads="12",
        memory="2G",
        job_name="bcftools_MAF"
    log:
        err="logs/bcftools_MAF/{specie}/{genome}.log"
    shell:
        "bcftools +fill-tags {input} -o {output}"# -- -t MAF"
```

```
rule coverage_MAF_plots:
    input:
        "VCF/varscan/{specie}/{genome}/pool_varscan_{specie}_{genome}_baSNP_contig_MAF.vcf"
    output:
        ADP="plots/{specie}/{genome}/ADP.png",
        MAF="plots/{specie}/{genome}/MAF.png",
        adp="plots/{specie}/{genome}/ADP_stats.txt",
        sd="plots/{specie}/{genome}/SD_stats.txt"
    params:
        ADP="plots/{specie}/{genome}/ADP",
        MAF="plots/{specie}/{genome}/MAF",
        threads="10",
        memory="1G",
        job_name="coverage_MAF_plots"
    log:
        err="logs/coverage_MAF_plots/{specie}/{genome}.log"
    shell:
        "python3.10 script_coverage_MAF.py -v {input} -m {params.MAF} -c {params.ADP} -s
{output.sd} -a {output.adp}"



rule vcftools_filter:
    input:

vcf="VCF/varscan/{specie}/{genome}/pool_varscan_{specie}_{genome}_baSNP_contig_MAF.vcf",
        adp="plots/{specie}/{genome}/ADP_stats.txt",
        sd="plots/{specie}/{genome}/SD_stats.txt"
    output:

temp("VCF/varscan/{specie}/{genome}/pool_varscan_{specie}_{genome}_baSNP_contig_MAF_filt
ered.recode.vcf")
    params:
```

```
name="VCF/varscan/{specie}/{genome}/pool_varscan_{specie}_{genome}_baSNP_contig_MAF_filtered",
        threads="12",
        memory="2G",
        job_name="vcftools"
    log:
        err="logs/vcftools_filter/{specie}/{genome}.log"
    shell:
        """
        DP=$(< {input.adp});
        SD=$(< {input.sd});
        min=$(echo | awk "{{ print $DP - $SD }}");
        max=$(echo | awk "{{ print $DP + $SD }}");
        vcftools --vcf {input.vcf} --min-meanDP $min --max-missing-count 2 --max-meanDP $max --recode --out {params.name}
        """

rule vcftools_scaff:
    input:
"VCF/varscan/{specie}/{genome}/pool_varscan_{specie}_{genome}_baSNP_contig_MAF_filtered.recode.vcf",
    output:
temp("VCF/varscan/{specie}/{genome}/pool_varscan_{specie}_{genome}_baSNP_contig_MAF_filtered_scaff.recode.vcf")
    params:
name="VCF/varscan/{specie}/{genome}/pool_varscan_{specie}_{genome}_baSNP_contig_MAF_filtered_scaff",
        threads="12",
        memory="2G",
        job_name="vcftools_scaff"
    log:
```

```
        err="logs/vcftools_scaff/{specie}/{genome}.log"
    shell:
        "vcftools --vcf {input} --maf 0.05 --chr A01 --chr A02 --chr A03 --chr A04 --chr A05 --chr A06 --chr A07 --chr A08 --chr A09 --chr A10 --recode --out {params.name}"



rule snpeff:
    input:
        "VCF/varscan/{specie}/{genome}/pool_varscan_{specie}_{genome}_baSNP_contig_MAF_filtered_scaff.recode.vcf"
    output:
        temp("VCF/varscan/{specie}/{genome}/pool_varscan_{specie}_{genome}_baSNP_contig_MAF_filtered_scaff_annotated.vcf")
    params:
        threads="2",
        memory="20G",
        job_name="snpeff"
    log:
        err="logs/snpeff/{specie}/{genome}.log"
    shell:
        "java -Xmx8g -jar ~/snpEff/snpEff.jar Z1 {input} > {output}"


rule bedtools_subtract:
    input:
        vcf="VCF/varscan/{specie}/{genome}/pool_varscan_{specie}_{genome}_baSNP_contig_MAF_filtered_scaff_annotated.vcf"
    output:
        "VCF/varscan/{specie}/{genome}/pool_varscan_{specie}_{genome}_baSNP_contig_MAF_filtered_scaff_annotated_centro.vcf"
    params:
        threads="12",
```

```
        memory="2G",
        job_name="bedtools_subtract"
    log:
        err="logs/bedtools_subtract/{specie}/{genome}.log"
    shell:
        "bedtools subtract -a {input.vcf} -b "+str(centro)+" > {output}"


rule count_lines:
    input:

raw="VCF/varscan/{specie}/"+str(genome)+"/pool_varscan_{specie}_"+str(genome)+"_baSNP.vcf",

filtered="VCF/varscan/{specie}/"+str(genome)+"/pool_varscan_{specie}_"+str(genome)+"_baSNP_c
ontig_MAF_filtered.recode.vcf",

maf="VCF/varscan/{specie}/"+str(genome)+"/pool_varscan_{specie}_"+str(genome)+"_baSNP_conti
g_MAF_filtered_scaff.recode.vcf",

centro="VCF/varscan/{specie}/"+str(genome)+"/pool_varscan_{specie}_"+str(genome)+"_baSNP_co
ntig_MAF_filtered_scaff_annotated_centro.vcf"
    output:
        raw="raw_number_SNPs_{specie}_"+str(genome)+".txt",
        filtered="filtered_number_SNPs_{specie}_"+str(genome)+".txt",
        maf="maf_number_SNPs_{specie}_"+str(genome)+".txt",
        centro="centro_number_SNPs_{specie}_"+str(genome)+".txt"
    params:
        threads="12",
        memory="2G",
        job_name="count_lines"
    log:
        err="logs/count_lines/{specie}/"+str(genome)+".log"
    shell:
        "grep -v '##' {input.raw} | wc -l > {output.raw};"
        "grep -v '##' {input.filtered} | wc -l > {output.filtered};"
```

```
"grep -v '##' {input.maf} | wc -l > {output.maf};"
"grep -v '##' {input.centro} | wc -l > {output.centro};"
```
